# Neocortical temporal patterning by a two-layered regulatory network

**DOI:** 10.64898/2025.12.18.695250

**Authors:** Guohua Yuan, Zhe Zhao, Xiaoxiang Dong, Xu-Wen Wang, Qiangqiang Zhang, Xiangyu Yu, Ming Zhu, Jiawei Yu, Dan Zhang, Nantao Zhang, Zhiheng Xu, Lei Dai, Yang-Yu Liu, Song-Hai Shi, Yinqing Li

## Abstract

In the developing neocortex, a diverse array of neurons with defined types and abundances are systematically generated by a limited population of radial glial progenitors (RGPs) as they undergo successive fate changes. The molecular regulation behind this intricate temporal patterning remains elusive. We undertook in-depth single-cell multi-omics analyses, discovering a two-layered regulatory framework at the core of this process. Central to this are global temporal regulators positioned above a temporal network, consisting of series of transcriptional factor (TF) hub groups under sequential state transitions. This temporal network operates not by restricting TF expressions to discrete temporal windows, but through coordinated transcriptional and chromatin-accessibility dynamics that modulate transient TF regulatory activity. Moreover, global temporal regulators specify the duration of each cascading stage and, consequently, the number of progenies generated at each stage. Loss of global temporal regulators protracts RGP lineage progression, whereas their increased activity accelerates it. These findings suggest a two-layer temporal regulatory system controlling RGP lineage progression and neural progeny output duality in mammalian neocortical development.

## Background

In the developing neocortex, radial glial progenitors (RGPs) in the ventricular zone (VZ) serve as the primary progenitor cells^1^. They undergo a precisely timed sequence of stages: self-amplification, deep-layer neurogenesis, and then superficial-layer neurogenesis^2–5^. This sequence, termed temporal patterning, is integral for producing neurons with specific types and quantities corresponding to their stages of origin^6–9^. Both *in vivo* and *in vitro* studies underscore this patterning^3,10,11^, hinting at a robust intrinsic program^6,7^. Furthermore, brain cell atlases have revealed comparable cellular compositions across mammalian brains, despite their significant differences in size^12,13^. This implies that the temporal patterning is not only robust and conserved, but also flexible enough to be tuned differently across mammals. How these two seemingly paradoxical properties are achieved is unknown.

In the classical model of developing *Drosophila* nervous system, temporal patterning is orchestrated by a series of transcription factors (TFs), each with a defined temporal expression window^14–17^. This on-and-off expression regulation of TFs establishes developmental stages in the *Drosophila* ventral nerve cord and optic lobe^15,17–21^. Yet, in mammals, despite extensive genetic research^22–26^ and single-cell atlases^8,9,27–33^ profiling the transcriptional details of the developing neocortex, no analogous temporal TF expression window has been identified^34^. Consequently, the molecular regulators dictating temporal patterning in the developing mammalian cortex remain elusive^14–17,34^.

Here, we introduce an experimental and analytical framework aimed at assessing multilayered gene regulation within single cells to decode the RGP multi-state progression. By focusing on the synchronous dynamics of both gene expression and chromatin accessibility, we identified and functionally characterized a set of TF groups that govern sequential expression programs crucial for every RGP stage. We termed these TFs transition-hub TFs and found that they act within a temporal network. This temporal network operates not by bounding transition-hub TF to temporal expression windows, but instead, by coordinated transcriptional and chromatin dynamics that modulate TF regulatory action at their cis-regulatory elements. Analysis of factors acting upstream of the transition-hub TF groups revealed global temporal regulators determines the duration of each RGP fate shift across the entire RGP progression, consequently modulating the quantity of each specific progeny outputs. This two-layered regulatory network consisting of transition-hub TFs and global temporal regulators provides an analytical foundation and a direct means for experimental interrogation of temporal patterning, opening up new possibilities for uncovering molecular regulators of cell-state specification in diverse transitioning systems.

### In-depth mapping of TFs and regulatory elements with single-cell multi-omics

To explore the regulatory programs underlying RGP lineage progression and neocortical development, we employed in-depth multi-omics approaches to systematically analyze gene expression and regulation in RGPs. We first developed a highly sensitive technology for joint profiling of ATAC and RNA expression in the same single cells (scAnR-seq). Our approach prioritized in-depth coverage rather than scalability, in contrast to previous co-assays^35–37^, to enable deep characterization of regulatory programs more effectively. scAnR-seq splits single cell nuclear and cytoplasmic content using magnetic beads, allowing independent optimization for ATAC-seq and RNA-seq (**Figure S1-1a and Figure 1a**). We comprehensively screened a total of 120 parameters (**Figure S1-1 b-h and Table S1**), arriving at a robust, reliable protocol with unparalleled sensitivity (e.g. 5.8E+04 unique fragments and 4.8E+03 unique genes in single embryonic cells) (**Figure S1-2**), surpassing even dedicated stand-alone single-cell assays^38,39^(**Figure S1-2 h, k**). Critically, the high-quality data generated by scAnR-seq unveils key regulatory components previously obscured by data sparsity^35–37^ (**Figure 1b,c**) such as low-abundance TFs and distal regulatory elements, both crucial in shaping dynamic expression programs across cell states^40–44^. Notably, distal CREs have particularly been found in various developmental contexts to provide flexible, long-range control of gene expression, supporting finer regulatory tuning of developmental processes than what promoter regions alone can achieve^36^.

**Figure 1:**
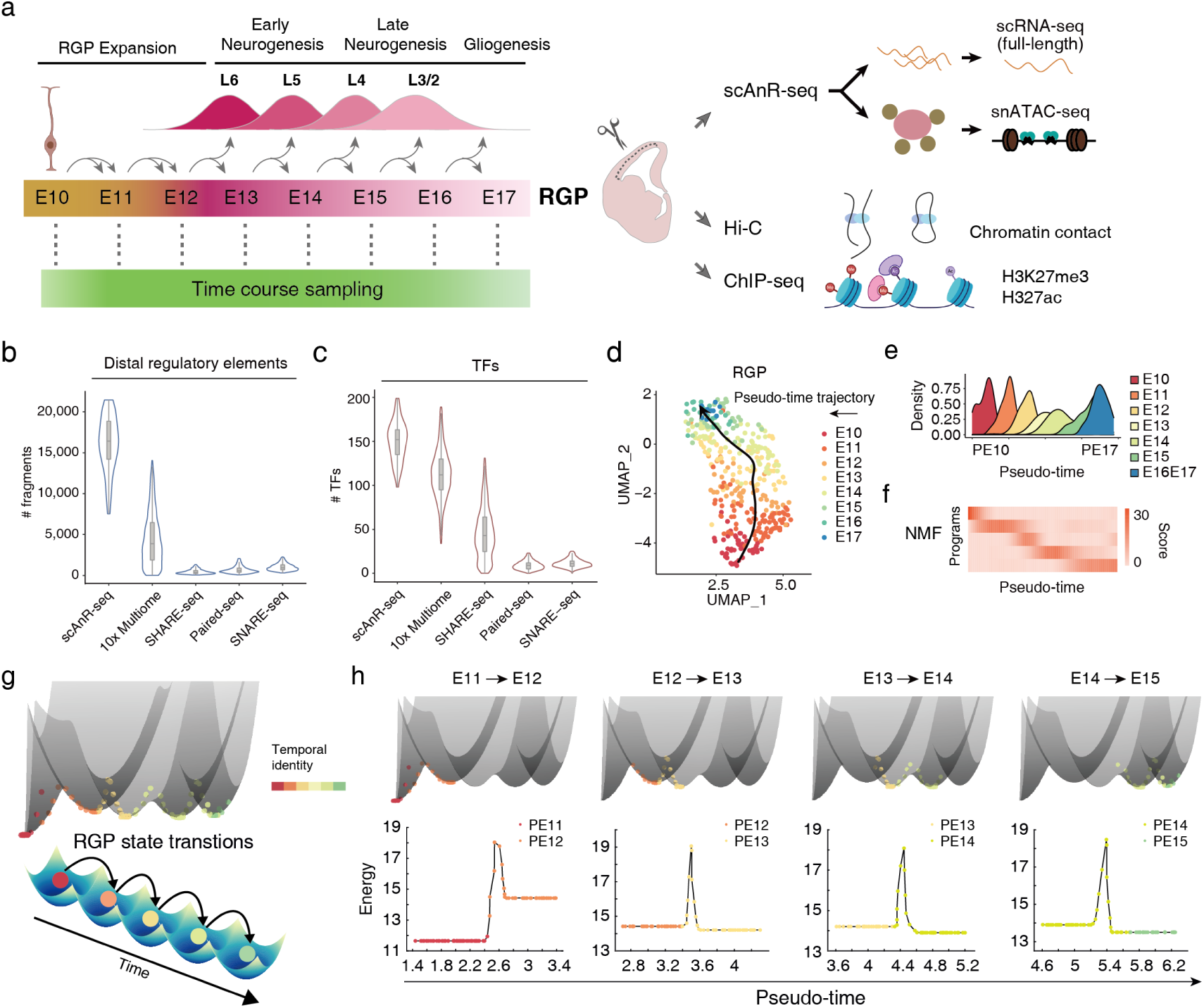
Multi-omics profiles along pseudo-time recapitulate RGP multi-state transitions. **(a)** Schematic of time-series sample collection covering cortex development. The scAnR-seq data was generated systematically at 24-hour intervals between embryonic day (E) 10 and E17. This involved collecting 4 mouse brains at each interval. During the late stage (E13-E17), when RGPs are less predominant in the VZ, we supplemented sampling with an additional 2∼3 mouse brains at each time point. H3K27ac, H3K27me3, and HiC data was generated for every two days. **(b and c)** Quantification of detected TFs (b) and detected fragments in distal regulatory regions (c) in cells derived from brain tissues by scAnR-seq (this study), 10x Multiome, SHARE-seq^36^, Paired-seq^37^ or SNARE-seq^35^. Box inside violin plot denotes 25^th^, 50^th^, and 75^th^ percentiles; the whisker length represents 1.53 interquartile range. **(d)** UMAP showing pseudo-time trajectory of RGPs. Cells are colored by sampling time. **(e)** Density plot showing the spread of sampling time along pseudo-time trajectory. **(f)** Gene program score from NMF along pseudo-time**. (g)** A series of energy-potential wells resonated with RGP temporal identities along pseudo-time. Top: Energy landscape of the RGP lineage progression, determined on scAnR-seq profiles using a statistical physics method^48^. In analogy to the Boltzmann distribution, lower energy states correspond to configurations with more cells of similar scAnR-seq profiles, representing stable states in the lineage progression, and higher energy states represent configurations with fewer cells of similar scAnR-seq profiles, corresponding to transition between stable states. Each dot represents a single cell. Bottom: An energy-barrier crossing model along RGP lineage progression. Cells are colored by their temporal identity along pseudo-time trajectory. **(h)** Projection of energy landscape in Fig. 1g corresponding to each transition along pseudo-time. Cells are colored by their temporal identity.

### Multi-omics profiles along pseudo-time recapitulate RGP multi-state transitions

We applied scAnR-seq to characterize the regulatory basis of RGP progression. We harvested the VZ tissue, which contains the majority of RGPs, every 24 hours, capturing all phases from RGP self-expansion to superficial-layer neurogenesis (**Figure 1a**). This involved collecting 4 mouse brains at each interval (8 key time points in total over E10-E17). During the late stage (E13-E17), when RGPs are less predominant in the VZ, we supplemented sampling with an additional 2∼3 mouse brains at each time point, employing FlashTag labeling to enhance RGP capture^45^. Notably, clustering of scAnR-seq profiles^46^ yielded much clearer cell-type differentiation compared to separate ATAC or RNA analysis (**Figure S1-3**), highlighting the synergistic value of integrating chromatin accessibility and gene expression data.

Pseudo-time analysis of our scAnR-seq profiles closely paralleled (R > 0.9) their experimental sampling time (**Figure 1d, e**). RGPs from consecutive days substantially overlapped in pseudo-time and formed a continuum of developmental trajectory, indicating that intermediate temporal stages of RGP development are well sampled in the dataset. This alignment enabled a direct correlation between pseudo-time and real-time, assigning a temporal identity to each RGP profile (**Figure 1e**). This is corroborated by differentially expressed (DE) genes over pseudo-time, which were enriched in genomic regions exhibiting dynamic epigenetic shifts, as highlighted by changes in H3K27ac, H3K27me3 ChIP-seq, and Hi-C topological contacts (**Figure S1-4 a-f**). Importantly, our pseudo-time analysis encompassed all RGP states observed from a comprehensive atlas study^27^ (**Figure S1-4 g**). In sum, RGP progression was adequately sampled and encapsulated by scAnR-seq profiles along pseudo-time.

Using scAnR-seq profiles and pseudo-time, we examined the temporal heterogeneity of RGP in molecular programs. An unbiased analysis of genes across RGP pseudo-time via non-negative matrix factorization (NMF)^47^ and principal component analysis (PCA) consistently revealed temporally specific, correlated gene expression shifts (**Figure 1f and Figure S1-4 h**). These shifts occurred in a wave-like pattern, with RGP progression seen as sequential cell states, each marked by a stable phase in a program, interspaced by transitions between adjacent programs. Notably, these RGP state transitions aligned with shifts in their developmental stages. To ensure the reproducibility of our transition analysis, we employed short-time Fourier transformation (STFT) on PCA-identified programs to identify prominent time-series changes, which allows for the detection of the high-frequency components reflecting how quickly gene expression patterns are shifting over pseudo-time. STFT robustly uncovered multiple transitions aligning with program shifts (**Figure S1-4 h**).

To further interrogate the state transitions, we examined the developmental trajectory of RGP progression via a statistical physics approach^48^, quantifying transition entropy to discern between stable states and transitions. The result revealed a series of energy-potential wells resonated with RGP temporal identities along pseudo-time (**Figure 1g,h**), illustrating the RGP progression as a barrier-crossing model in a multi-stable potential landscape. This landscape of barriers and potentials manifest from intricate biological interactions, such as gene expression regulation. We quantified cell density along pseudo-time to distinguish between stable developmental states represented by densely populated regions and transitions between these states represented by sparsely populated regions (**Figure 1g,h**). This distinction highlights the periods of stability where RGPs maintain their developmental state and temporal identity versus the periods of transition where RGPs undergo more significant regulatory changes and move from one developmental state to another developmental state. We identified multiple temporal transitions which represent dynamic regulatory events that regulate RGPs from one developmental state to the next.

### Temporal network underlying RGP transitions: TF regulatory activities are asynchronous and transitory

How is the progression through multiple transitions in the broad concerted developmental pathway governed? To investigate the regulation overarching the RGP progression, we first examined the relation between chromatin accessibility and gene expression across pseudo-time, focusing on two key metrics: (i) Dynamics of TF activity vs. expression and (ii) Dynamics of cis-regulatory element (CRE) accessibility vs. target gene expression.

For TF dynamics, we found a global coordination between TF expression and activity, as captured by TF-motif accessibility via ChromVAR^49^ (**Figure S2-1 a**). This was evident in known regulators such as *Hmga2* and *Neurog2*, which displayed dynamic changes in the early and mid-stages (**Figure S2-1 b**), reflecting their roles in self-renewal^26,50^ and neurogenesis initiation^51–53^, respectively. These expression changes were broadly confirmed using in situ hybridization over time (**Figure S2-2**).

For the CRE dynamics, we associated distal CREs and proximal CREs with target genes using their statistical linkage^54–56^ (**Figure 2a and Figure S2-3 a**). Given that CREs and target genes often interact through chromatin folding^57,58^, we refined this association using Hi-C data, leading to 58,383 CREs and 110,465 CRE-gene associations. A significant fraction of these CREs were recognized as brain enhancers in ENCODE^59^, overlapped with regulatory features such as RGP H3K27ac and H3K27me3 peaks, and featured dense associations at key developmental regulators^36^ (**Figure 2 b,c and Figure S2-3 b-d**). Remarkably, during RGP progression, distal CREs, with their deep representation in our scAnR-seq data, particularly stood out in their numbers and correlation strength with gene expression (**Figure 2e and Figure S2-3 d**), emphasizing their pivotal role in cell state determination. This correlation, together with a slight lead in CRE dynamics over expression dynamics (**Figure 2d,f and Figure S2-3 e**), enabled prediction of a cell’s impending transcriptional state given its current CRE state via chromatin potential^36^, revealing a developmental path readily aligned with our pseudo-time estimations (**Figure 2g**). Collectively, these analyses point to correlated changes in key regulatory elements, i.e. TF and CREs, suggesting coordinated regulatory programs guiding developmental transitions.

**Figure 2:**
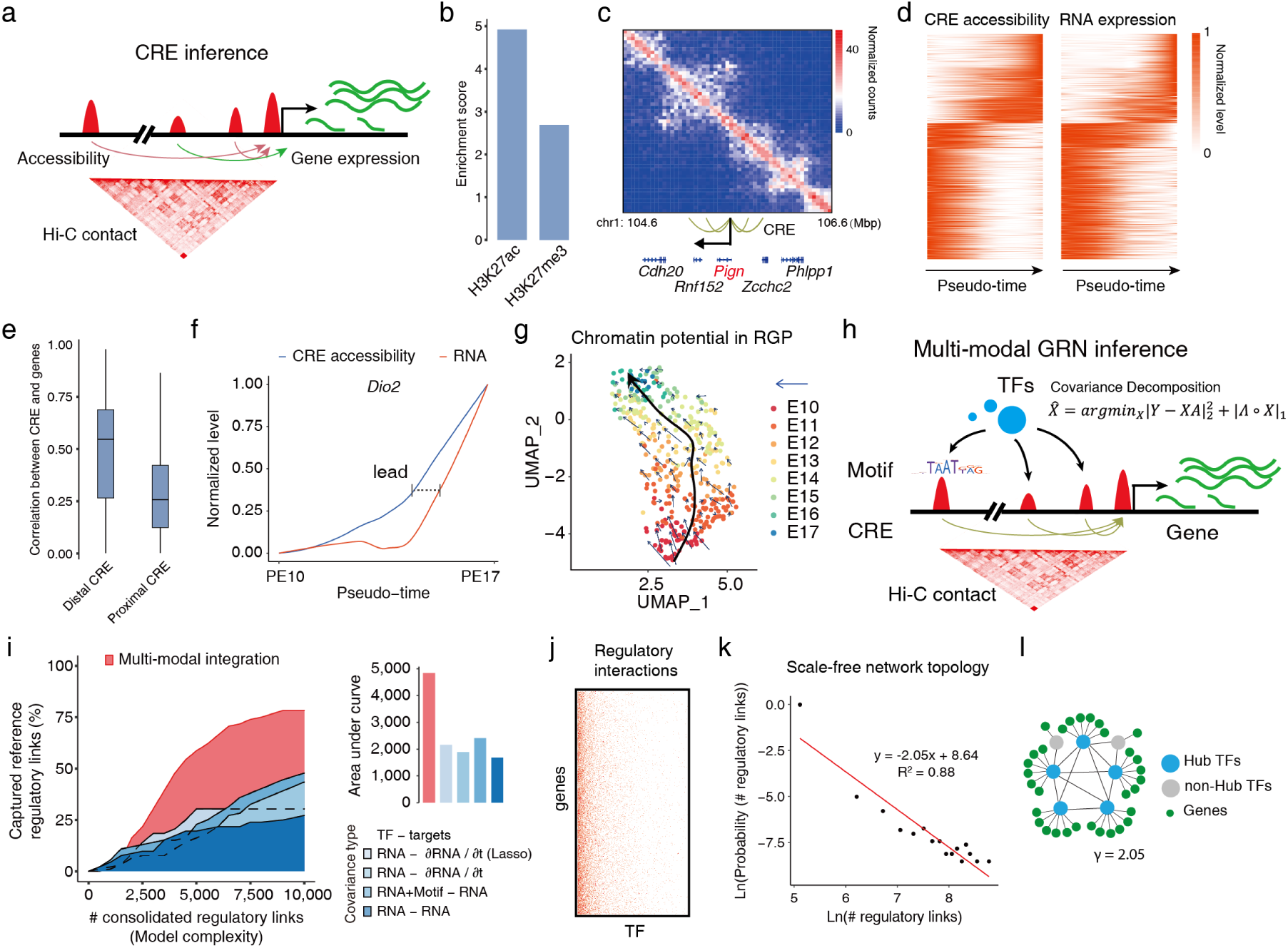
Global chromatin and gene expression dynamics determine TF-target regulatory links. **(a)** High-fidelity CRE inference based on peak-to-gene associations derived from covariation between chromatin accessibility (ATAC-seq) and gene expression (RNA-seq) and incorporating Hi-C chromatin contact information. **(b)** Enrichment score (odds ratio) of CREs in H3K27ac and H3K27me3 peaks. **(c)** Representative locus showing the *Pign* CREs locate in regions Hi-C contact. **(d)** Heatmaps showing single-cell gene expression and CRE accessibility dynamics over pseudo-time. Single cells are ordered by pseudo-time. **(e)** Boxplot showing correlation between gene expression and distal versus proximal CREs. Box denotes the 25th, 50th, and 75th percentiles. **(f)** CRE accessibility (blue) and gene expression (red) dynamics of *Dio2* during its up-regulation, showing a modest lead in the CRE dynamics. **(g)** Chromatin potential aligned with pseudo-time. The direction of the arrow links a cell and its predicted most similar cell. **(h)** Model for single-cell ATAC and RNA data covariance. TFs bind to CREs, regulating target gene expression based on TF activity, abundance, or CRE accessibility. To infer regulatory interactions between TFs and target genes, we utilized a multi-modal inference approach, which incorporates the TF motifs in CREs as prior information and quantifies the statistical link strength among TFs, CREs, and genes dynamics. **(i)** Left: rarefaction analysis showing percentage of reference regulatory links captured versus number of consolidated regulatory links. The regulatory links were consolidated in descending statistical significance into the overall regulatory relation. Right: area under the curve for each rarefaction analysis. **(j)** The adjacency matrix showing inferred regulatory interactions between TFs (x axis) and target genes (y axis). **(k)** Distribution of numbers of linked target genes per TF. The distribution of numbers of regulatory links per TF followed a power law distribution. **(l)** illustration of the hub structure in the overall regulation relation.

TFs bind to CREs, regulating target gene expression based on TF activity, abundance, or CRE accessibility (**Figure 2h**). To infer regulatory links between TFs and target genes, we utilized a multi-modal inference approach^60,61^, which incorporates the TF motifs in CREs as prior information and quantifies the statistical link strength among TFs, CREs, and genes dynamics (**Figure S2-4 a**). We then consolidated these inferred links in descending statistical significance, forming an overall TF-target regulatory relation. For validation, we obtained a reference set comprising 1,722 pairs from 9 TF overexpression experiments in RGP and another 364 experiment-backed pairs from literature (**Table S2**). Our overall regulatory relation showed significant enrichment from TF overexpression datasets and captured 79.8% of the reference pairs (**Figure 2i and Figure S2-4 b, c**). Notably, this enrichment was not attributed to false positives, as rarefaction analysis showed the reference-pair coverage rose quickly as the inferred regulatory links consolidated (**Figure 2i**). And, when randomizing the statistical covariance in our scAnR-seq data, this enrichment vanished, further affirming the validity of our approach (**Figure S2-4 d, e**). Compared to conventional inferences that mostly reply on RNA co-expression alone^60–64^, the overall regulatory relation, grounded in multi-modal inferences, demonstrated a markedly higher reference capture rate (**Figure 2i**). Within this regulatory relation, we found the distribution of regulatory links per TF conformed to a scale-free topology with power-law exponent γ=2.05, typical of gene regulatory networks^65,66^ (**Figure 2 j-l**), highlight hub-TFs as the core in the regulatory network.

To understand regulatory activities across developmental transitions, we next segmented pseudo-time into consecutive stages and quantified the activity of each TF-target regulatory link based on the strength of concurrent changes in the linked target and TF within each stage, reasoning that if a target were primarily driven by a TF during a specific phase, any significant changes in target expression should coincide with salient shifts in TF expression or motif accessibility within the same time frame. Our analysis revealed that the regulatory link activities were highly transitory (**Figure 3a, b**). A substantial fraction (72.3%) were mostly active within individual time stages (**Figure 3a**), on average accounting for nearly three-quarters (77.8%) of regulatory links per TF. In addition, the distribution of active regulatory links varied greatly across different time stages and TFs. For instance, “early-stage” links were prominent in *Mybl2* and *Hbp1* (11.1%, 7.1%), whereas both “mid-stage” and “late-stage” links were more balanced in *Sox9*, demonstrating the transitory and asynchronous nature of activities among regulatory links by different TFs (**Figure 3b**). Importantly, the activities of regulatory links showed minimal dependency on the absolute expression levels of either TFs or targets (**Figure S3-1 a-c**), suggesting that this analysis is robust against potential expression-level biases in single-cell omics data^67^.

**Figure 3:**
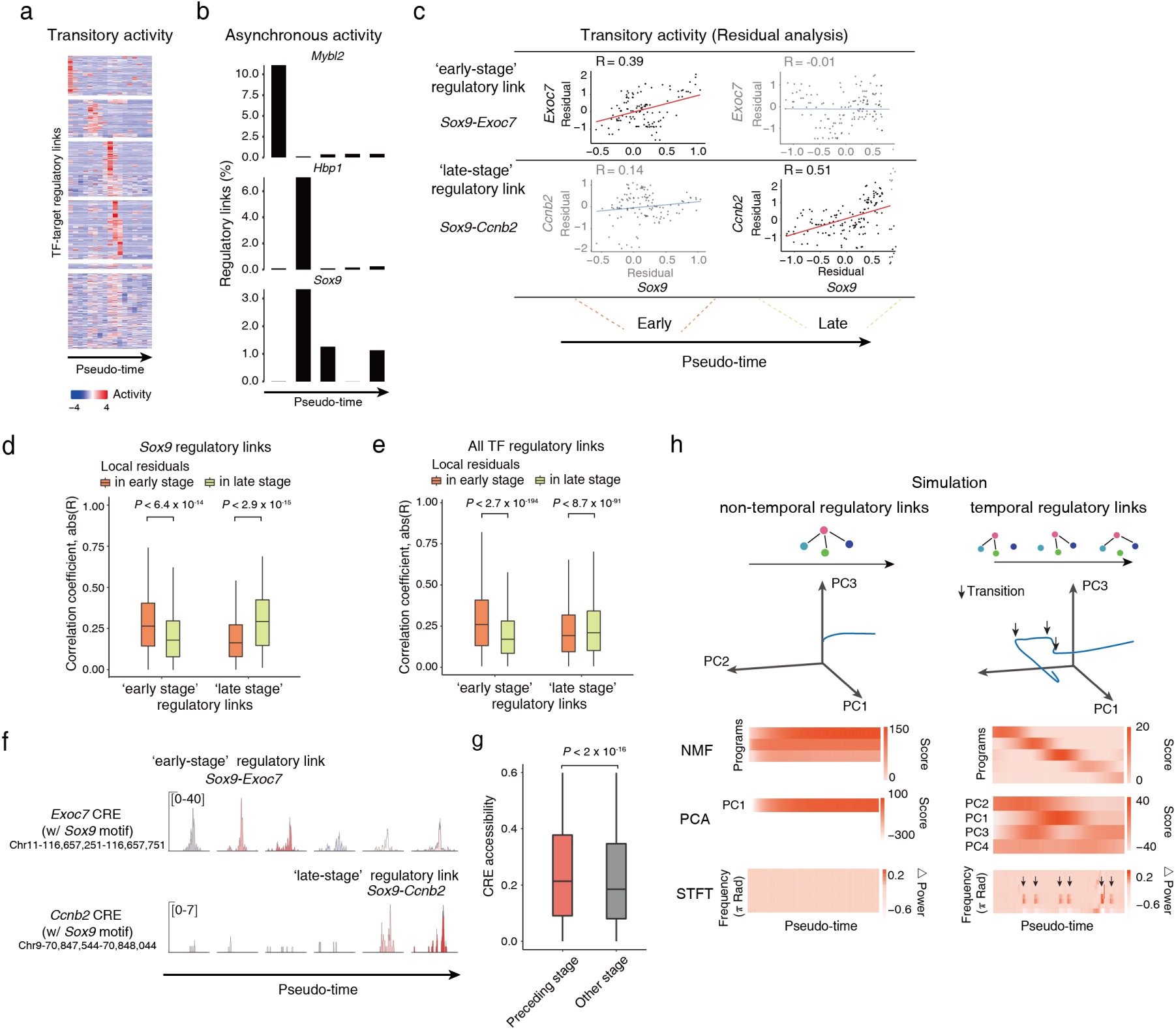
Activity of regulatory links is asynchronous and highly transitory. **(a)** Cluster analysis of TF-gene regulatory links by activity along pseudo-time (Methods). The pseudo-time was divided into 17 consecutive time bins using sliding windows, the activity of each regulatory link was quantified. Horizontal clusters of regulatory links were computed using principal components of the activity of regulatory links. Heatmap showing activity of each regulatory link along pseudo-time. **(b)** Percentage of regulatory links of indicated TFs by stages across pseudo-time. The temporal boundaries that differentiate activities of regulatory links of neighboring clusters in (a) were obtained by maximizing likelihood ratio and were used to partition stages along pseudo-time. Counting only regulatory links with strong activity in each stage (Methods). **(c)** Scatter plots showing single-cell residuals of *Sox9* and targets of two regulatory links with different temporal activity: *Sox9* and *Exoc7*, mostly active in early stage (top); and *Sox9* and *Ccnb2*, mostly active in late stage (bottom). Line represents linear regression fitting. **(d)** Box plot showing higher correlation between single cell residuals of *Sox9* and target when the regulatory link exhibits strong activity. P < 6.4 × 10^-14^ in early stage; P < 2.9 × 10^-15^ in late stage. Wilcoxon rank-sum test. **(e)** Extension of the analysis in (d) to all TFs. Box plot showing higher correlation between single cell residual variability of a given TF and target when the regulatory link exhibits strong activity. P < 2.7 × 10^-194^ in early stage; P < 8.7 × 10^-91^ in late stage. Wilcoxon rank-sum test. **(f)** IGV track showing accessibility increase at a CRE that specifically links *Sox9* and target precedes strong activity of the regulatory link. Peaks are colored by activity strength of the regulatory link as in (a). **(g)** Extension of the analysis in (f) to all TFs. Box plot showing higher accessibility at CRE that specifically links a TF-target precedes strong transitory activity of the corresponding TF-target regulatory link. P < 2 × 10^-16^. Two-sided unpaired Wilcoxon rank-sum test. **(h)** Ordinary differential equation (ODE) simulation of dynamics without (left) and with link temporality (right).

The transitory activity of regulatory link necessitated interrogation with matched temporal resolution, thus we leveraged the time series single-cell transcriptomes and regressed them against pseudo-time to obtain local “residuals” (defined as the difference between single cell and pseudo-time dependent expression level), which represent low yet detectable perturbations in TF and responses in target expression. We found that the correlation of single-cell local residuals of TF and target was significantly higher when the regulatory links were active (**Figure 3c-e and Figure S3-1 c**), confirming their temporally selective regulation.

Furthermore, the transitory regulatory activities were preceded by changes in the chromatin accessibility at the individual CREs that specifically linked the TF and target pairs (**Figure 3f, g and Figure S3-1 d**), suggesting that epigenetic changes could contribute to their priming.

The asynchronous and transitory activities of regulatory links across RGP progression suggested their potential governance by a temporal network, i.e., a time-varying regulatory structure consisting of links that exist only intermittently^68^. Such a network allows a complex system to be steered within sequential subspaces confined by temporal links^68–70^, facilitating sequential transitions through its stage-wise dynamics. To investigate this principle in RGP progression, we created a network model with degree distributions sampled from the RGP overall regulatory relation and applied ordinary differential equations (ODEs)^71^ to simulate its dynamics. Additionally, the link temporality was modeled with explicit activation and deactivation of time-varying subsets of links. We found that the multiplicity of sequential transitions was recapitulated in simulations with asynchronous link activity, as indicated by consecutive programs in NMF and PCA, as well as by multiple sequential spikes in the STFT analysis (**Figure 3h right**). In contrast, the STFT spike was entirely absent in simulations without link temporality (**Figure 3h, left**). Hence, the dynamics of multiple transitions are consistent with asynchronous activities of regulatory links. Together, these results reveal temporally selective regulatory activities as a prominent feature of RGP lineage progression and enables complex systems to transition through sequential states.

### Temporal network comprises transition-wise hub TF groups

Our identification of temporally selective regulatory interactions suggested that RGP lineage progression is governed by a time-varying gene regulatory network where interactions between TF and target genes are active at specific times, forming a series of temporal subnetworks.

Therefore, we analyzed the subnetworks of TFs and target genes with active regulatory interactions during each RGP developmental state transition (**Figure 4a-e and Figure S4-1 a-c**). Within these subsets, we ranked TFs by the number of linked target genes. While there was a consistent TF hierarchy, a large fraction (23.6%) of TFs, including *Hes1*, *Prdm16*, and *Lef1*, were markedly (> 5 fold) promoted to top-ranked TFs in certain transitions, altering top TF list for each transition (**Figure 4b and Figure S4-1 b, c**). This transition-wise analysis revealed a changing regulatory hierarchy over time (**Figure 4b and Table S3**), e.g., *Hmga2*, *Sox9* and *Hbp1* in early and middle transitions, and *Lhx2* and *Lef1* in late transitions, orchestrating dynamic gene expression in sequence throughout RGP progression. We refer to these TFs as transition-hub TFs. The regulatory hierarchy proved robust, as the transition-wise TF rankings persisted even when 40% of the regulations were randomly shuffled (R = 0.96; **Figure S4-1 d**). Moreover, targets of transition-wise top-ranked TFs were significantly enriched in the same topologically associating domains (TADs), which delineate co-regulatory domains^58,72^, indicating the involvement of chromatin organization in hub regulation (**Figure 4d, e and Figure S4-1 e**).

**Figure 4:**
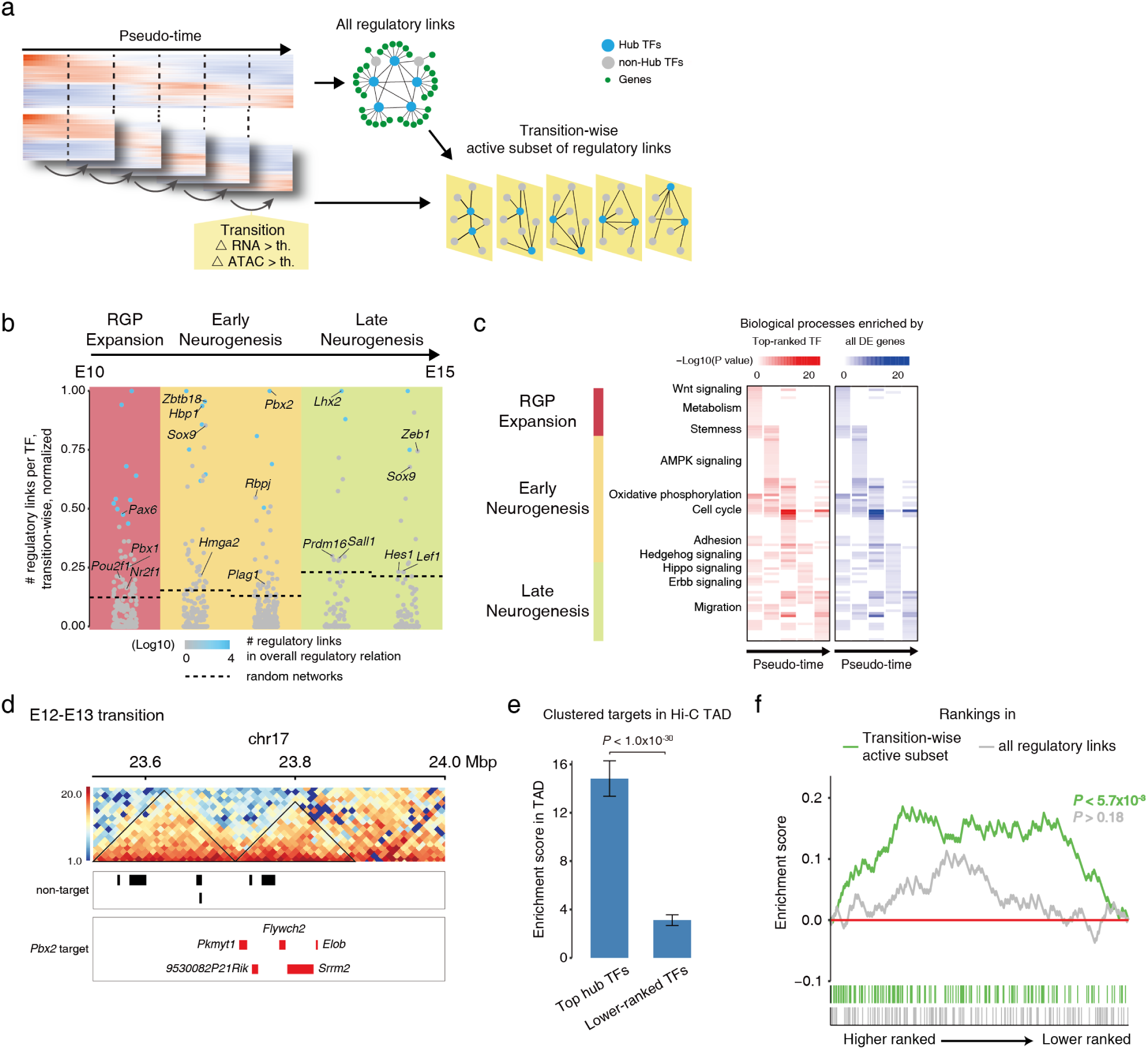
Temporal network comprises transition hub TFs. **(a)** Overview of transition-wise analysis of regulatory links. Thresholds for active subset was set based on differential motif accessibility or differential expression with P value < 0.05, Wilcoxon rank-sum test. **(b)** Transition hub TFs associated with a significantly higher number of regulatory links in transition-wise analysis, compared to TFs in random networks without hub structure. Numbers of regulatory links normalized as fractions of the maximum in transition-wise analysis. **(c)** Heatmaps showing enrichment of KEGG and GO biological processes using direct targets of top-ranked TFs (red) or all differentially expressed genes (blue) in each transition. Representative processes are provided for each transition. **(d)** Hi-C contact matrix (top) and gene model (bottom) showing enrichment of *Pbx2* target genes in topologically associating domains (TADs) (black line) in the E12-E13 transition. Red bars denote *Pbx2* target genes. Black bars denote *Pbx2* non-target genes. **(e)** Bar plot showing enrichment of target genes of top transition-hub TFs in Hi-C TAD. Enrichment score estimated by comparing targets in a TAD and in randomized TADs, one-sample Wilcoxon rank-sum test. Data are shown as mean ± std. P < 1.0 × 10^-30^, one-sided unpaired Wilcoxon rank-sum test. **(f)** GSEA running score showing TF enrichment in gene sets of regulators involved in cortical development (Table S4). Comparing TF ranks in the active subset (green) or all regulatory links (grey). P < 5.7 × 10^-3^ for active subset, P > 0.18 for overall regulatory relation, Kolmogorov-Smirnov test.

We next examined how the transition-wise regulatory hierarchy corresponded with dynamically changing biological processes across transitions. The processes governed by these transition-wise top-ranked TFs closely recapitulated those identified by all differentially expressed (DE) genes during respective transitions (**Figure 4c**). For instance, early transition TF targets were enriched for Wnt signaling and stemness, while later ones were enriched for oxidative phosphorylation, cell cycle, ErbB signaling, and migration, reflecting the temporal shift from self-amplification to deep-layer and then upper-layer neurogenesis (**Figure 4c**).

To systematically assess the roles of hub TFs, we employed GeneFormer, a foundational biological model trained on 30 million single-cell transcriptomics profiles^73^. This deep neural network model captures comprehensive complex gene regulatory relationships, enabling us to probe temporal subnetworks beyond traditional experimental capabilities. Focusing on the hub TFs from each temporal subnetwork, we upregulated their expression in silico during different RGP developmental state transitions and examined the effects on gene expression programs (**Figure S4-2 a**). Our analysis revealed that perturbing these hub TFs led to significantly greater changes in gene expression within their corresponding state transitions when they were hub TFs, compared with within other state transitions when they were not (**Figure S4-2 b**). This temporal specific effect was consistently observed across different developmental state transitions (**Figure S4-2 b**), supporting the notion that hub TFs regulate temporally selective gene expression programs. To further assess the role of hub TFs, we curated a list of regulators known to affect RGP lineage development when experimentally manipulated based on the previous studies and found a significant enrichment of hub TFs among these critical RGP lineage regulators (**Figure 4f and Table S4**). Notably, this enrichment substantially surpassed the overall regulatory relation (**Figure 4f and Figure S4-1 f**), highlighting the critical role of highly-promoted TFs in specific transitions. Collectively, these findings highlight the critical role of temporal subnetworks and their corresponding hub TFs in coordinating RGP state-specific gene expression programs and lineage transitions.

The distribution of TF targets was not random concerning transition-wise TF ranking. TFs with more targets were linked to broader functions, while those with fewer targets had specific roles, reflecting a functional hierarchy during RGP transitions (**Figure 5a and Figure S5-1**). In the early-stage transition (E11-E12), RGPs switch from symmetric to asymmetric cell division, producing distinct daughter cells: one remains attached while the other departs from the VZ surface^74^. We identified *Foxk1*, a previously less-known TF, prominently promoted in rankings during this E11-E12 transition and is directly regulated by a top-ranked TF *Hbp1* (**Figure 5a-c and Figure S5-2 a**). Both were ascribed to cell adhesion processes (**Figure 5a**).

**Figure 5:**
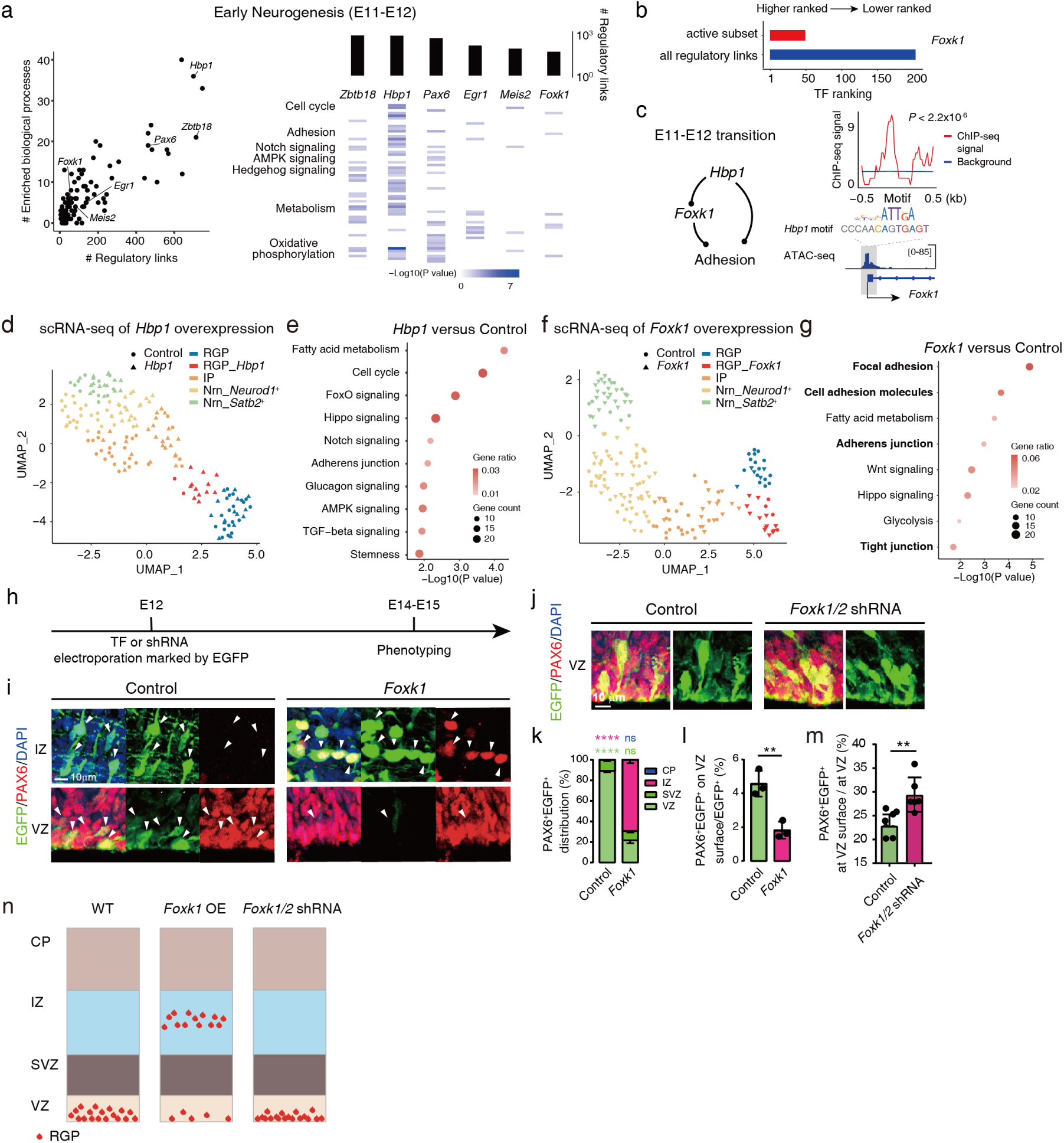
Hierarchical functional coordination of key processes by transition hub TFs. **(a)** Left: scatter plot showing the number of enriched KEGG and GO biological processes by TF direct targets versus the number of regulatory links in E11-E12 transition. Right: heatmap showing the enrichment score of representative KEGG and GO biological processes by direct targets of indicated TFs. Top: bar plots showing the number of regulatory links in E11-E12 transition. **(b)** Bar plots showing *Foxk1* promoted in TF ranking in E11-E12 transition. **(c)** Left: schematic showing the role of *Hbp1* and its downstream TF *Foxk1* in regulating adhesion in the E11-E12 transition. Right: ATAC-seq signal at the *Foxk1* promoter. Grey box indicates an identified *Hbp1* motif occurrence in the *Foxk1* promoter. The identified motif occurrence is a valid *Hbp1* binding site, as shown by enrichment of HBP1 ChIP-seq signal (GSE104247)^106^ in a cell line. **(d)** UMAP visualization of scRNA-seq from control and *Hbp1* overexpression samples. Cell shape indicates experimental conditions, and color indicates cell type assignment. **(e)** KEGG biological processes associated with differentially expressed genes in RGPs overexpressing *Hbp1*. **(f)** UMAP visualization of scRNA-seq from control and *Foxk1* overexpression samples. Cell shape indicates experimental conditions, and color indicates cell type assignment. **(g)** KEGG biological processes associated with down-regulated genes in RGPs overexpressing *Foxk1*. **(h)** Experimental timeline for *Hbp1* and *Foxk1* manipulation and phenotyping. **(i)** Representative coronal sections of dorsolateral telencephalon triple stained using anti-GFP (green), anti-PAX6 (red) antibodies, and a DNA dye (blue), following the experimental scheme shown in (h). Zoomed in view of VZ and IZ showing substantial discharging from the VZ and repositioning to the IZ. **(j)** Representative coronal sections of dorsolateral telencephalon triple stained using anti-GFP (green), anti-PAX6 (red) antibodies, and a DNA dye (blue), following the experimental scheme shown in (h). Zoomed in view of VZ and IZ. **(k-l)** Quantification of the distribution of PAX6^+^EGFP^+^ cells (k), and percentage of PAX6^+^EGFP^+^ cell on the ventral surface in EGFP+ cells (l). Data are shown as mean ± SEM. (j) **** P<0.0001, ***P=0.0004, **P=0.007; (k) **** P<0.0001; (l) **P=0.0067; Student’s t-test. N.S., not significant. **(m)** Quantification of distribution of percentage of PAX6^+^EGFP^+^ cell on the ventral surface in PAX6^+^EGFP^+^ cells in VZ. Data are shown as mean ± SEM. **P=0.008; Student’s t-test; N.S., not significant. **(n)** Summary of experimental results in the perturbation of *Foxk1*.

Overexpression of *Hbp1* influenced multiple pathways like cell cycle, Notch signaling, and adherens junction (**Figure 5d, e and Figure S5-2 b, c**), aligning with disrupted phenotypes seen in *Hbp1*-altered animals^75^. On the other hand, overexpressing *Foxk1* specifically impacted adhesion-related genes (**Figure 5f, g and Figure 5-2 d**). Supporting this, RGPs overexpressing *Foxk1* showed substantial discharge from the VZ (89% down to 20%), repositioning to the intermediate zone (IZ indicating *Foxk1* role in attachment destabilization (**Figure 5h,i,k,l**).

Conversely, knocking down *Foxk1* or together with its homolog *Foxk2* resulted in increased attachment to the VZ (**Figure 5h,j,m and Figure S5-2 e-j**). Our analysis suggests that adhesion destabilization, delicately modulated in hierarchy by transition-hub TFs, could be a key regulation for this transition (**Figure 5n**).

### Global temporal regulators regulate the pace of RGP progression

Within the temporal network, the activity of each TF’s regulatory links is largely suppressed outside its designated hub stage, eliminating the need for simultaneous silencing of its RNA expression (**Figure S6-1 a**). This stood in stark contrast to the *Drosophila* temporal TF chain, where chained singular TFs toggle between on-and-off expressions within successive temporal windows. In line with this distinction, even though some mouse orthologues of *Drosophila* cascade TFs, such as *Pax6* (orthologue to *Ey*) and *Pou2f1* (orthologue to *Pdm*), play roles in early transitions, their expression is not confined to strict temporal windows (**Figure S6-1 b,** c)^17,20^.

Across RGP progression, cross-regulation on the regulatory link activity occurs at both target CREs and TFs, resulting in target genes being dynamically regulated by different TFs at different times, as evidenced by shifted or attenuated target-expression dynamics across transitions (**Figure S6-1 d-f**). For instance, *Zbtb18* and *Zfx* were transition-hub TFs in the E11-E12 transition, while *Elk3* and *Rfx2* were transition-hub TFs in the next transition. Shifted target-expression dynamics were observed at *Pygo2*, a common target of *Zbtb18* and *Elk3*. Initially, *Pygo2* expression increased alongside *Zbtb18* expression. In the next transition, *Pygo2* expression continued to increase, while *Zbtb18* expression did not. Concurrently, *Elk3* expression increased as did accessibility of its cognate CRE at *Pygo2*, suggesting *Pygo2* regulation shifted from *Zbtb18*-mediated to *Elk3*-mediated regulation (**Figure S6-1 e**). At *Nek6*, a common target of *Zfx* and *Rfx2*, we observed attenuated target-expression dynamics. *Nek6* and *Zfx* expression both increased initially but were then decoupled in the subsequent transition when the *Rfx2*-specific CRE at *Nek6* gained increased accessibility. This chromatin change likely counteracted the down-regulation of *Zfx2*, leading to attenuated dynamics and hence a stable level of *Nek6* expression (**Figure S6-1 f**). The overlapping regulation by cascading transition-hub TFs observed for *Pygo2* and *Nek6* represents a major feature in RGP progression. In a median of 94.4% of cases with shifted expression dynamics and 70.7% of cases with attenuated expression dynamics, such overlapping regulation was found, including transition-hub TFs as target genes outside of their hub stages (**Figure S6-1 g-i**). These findings suggest that coordinated transcriptional and chromatin interactions are associated with regulatory link temporal activity, enabling dynamic regulation without the need for toggling TF expression.

In the network landscape, extensive cross-regulation foster emergence of attractor states, which parallel the potential landscape observed along RGP pseudo-time (**Figure 1h**), evoking the model of energy barriers. These barriers signify gatekeeping stages in the regulatory network^76,77^, implying the existence of driving forces that navigate these barriers to guide global temporal coordination^17,78^. To identify molecular programs that may act upon the entire cascade progression, we searched for common upstream TFs whose targets were over-represented among transition-wise top-ranked TFs across all transitions (**Figure 6a-c and Figure S6-2 a, b**). This analysis uncovered significantly and robustly enriched common upstream TFs that exhibited large variance across the entire progression (**Figure 6a, c and Figure S6-2 a**), which we collectively refer to as global temporal regulators. Interestingly, members of the NFI family (*Nfia*, *Nfib*, and *Nfix*) emerge as the most dominant TFs ^79–82^. These global temporal regulators exhibit a continuously and progressively increasing expression patterns, suggesting a dosage-dependent global temporal control.(**Figure 6b and Figure S6-2 c,d**; Zhang et al).

**Figure 6:**
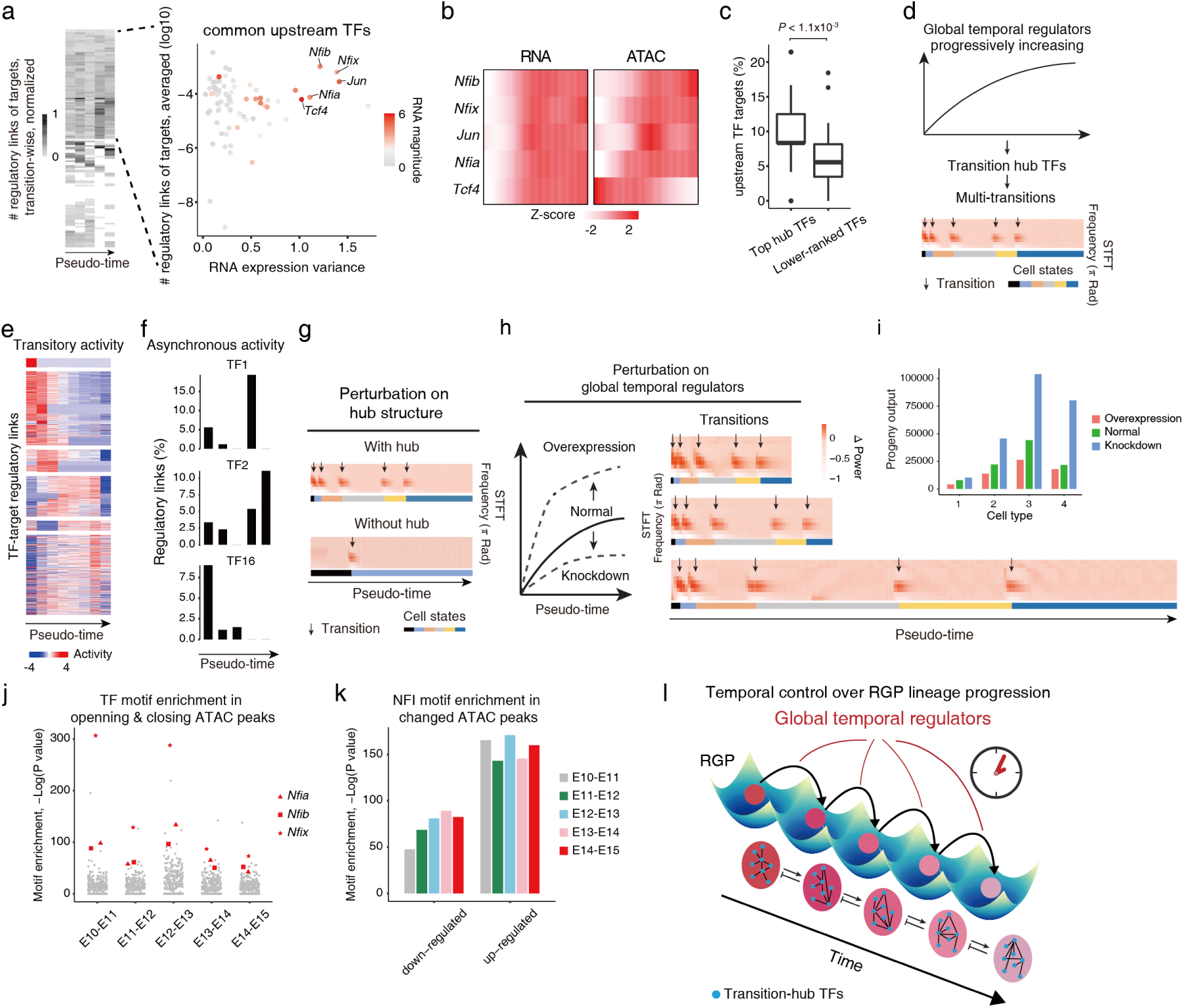
Global temporal regulators regulate the pace of RGP progression. **(a)** Left, heatmap showing the average number of regulatory links of targets of upstream TFs per transition. The numbers of regulatory links of targets by each common upstream TF were normalized as fractions of the maximum and averaged in transition-wise analysis. Common upstream TFs that have targets in all transitions were kept for analysis in scatter plot. Right, scatter plot showing the average number of regulatory links of targets versus the variance of RNA expression of common upstream TFs along pseudo-time. The numbers of regulatory links of targets by each common upstream TF were normalized as fractions of the maximum in transition-wise analysis and averaged across all transitions. **(b)** Heatmap showing progressively increasing dynamics of RNA expression (left) and motif accessibility (right) of TF highlighted in (A) along pseudo-time. **(c)** Common upstream TFs regulate a higher percentage of top transition-hub TFs compared to lower-ranked TFs. P < 1.1 × 10^-3^. One-sided paired Wilcoxon rank-sum test. Box plots denote 25th, 50th, and 75th percentiles. **(d)** Ordinary differential equation (ODE) simulation of dynamics on hub structure induced by progressively increasing expression pattern of a global temporal regulator (top) displayed multiple transitions (bottom). The global temporal regulator was identified based on its broad connections with top-ranked TFs in the network, as TFs top-ranked in the overall network were mostly also transition-wise top-ranked. **(e)** Cluster analysis of TF-gene regulatory links by activity along pseudo-time in simulation (Methods). Heatmap showing activity of each regulatory link along pseudo-time. **(f)** Percentage of regulatory links of indicated TFs by stages across pseudo-time. Counting only regulatory links with strong activity in each stage (Methods). **(g)** ODE simulation of dynamics induced by progressively increasing expression pattern with or without hub structure. STFT analysis characterizes the transitions along pseudo-time. Arrows point to middle of transitions. Colors indicate cell states interspaced by transitions. **(h)** ODE simulation of dynamics in response to increased or decreased slope of the progressively increasing expression pattern of a global temporal regulator. Left: increased slope (emulating overexpression) or decreased slope (emulating knock-down). Right: STFT analysis characterizes the transitions along pseudo-time. Arrows point to middle of transitions. Colors indicate cell states interspaced by transitions. **(i)** Bar plot showing the fold changes in the abundance of different cell types, calculated based on the simulations in (h). The abundance of progeny output is proportional to the duration of each state. **(j)** Scatter plot showing the enrichment of TF motifs in the sets of changing ATAC peaks at each transition. NFI motifs are highlighted in red, showing their significant enrichment across various transitions. **(k)** Scatter plot showing the enrichment of NFI motifs in the sets of opening and closing ATAC peaks at each transition. **(l)** Model of energy-barrier crossing, showing transition hub structure and global temporal regulators in regulation of sequential transitions along RGP lineage progression.

We next sought to gain further insight into the relation between the global temporal regulator, temporal network, and RGP multi-transition progression. However, the inherent complexity of the temporal network, characterized by extensive cross-regulation and feedback looks, makes any bottom-up experimental approach impractical. To circumvent these limitations, we opted for a computational strategy and constructed a network model, drawing degree distributions from the RGP overall regulatory relation (**Methods**).

We incorporated global regulators with broad connections to top-ranked TFs into a two-layer regulatory model and simulated the network dynamics (**Figure 6d**). Strikingly, the dynamics exhibited an emergent temporality of regulatory links (**Figure 6e,f**) and recapitulated multiple sequential transitions in consistent with RGP progression (**Figure 6d**). Perturbing the hub structure in simulation, we found it is necessary in transforming the regulator’s continuous pattern into the network’s staged temporal patterns via cross-regulation (**Figure 6g**).

Furthermore, we simulated the effect of downregulation or overexpression of global temporal regulator on sequential state transitions (**Figure 6h,i**). Interestingly, the simulation showed that overexpression promoted state transitions by enhancing regulatory changes required for transitions, whereas downregulation delayed state transitions by diminishing regulatory changes required for transitions. Moreover, altering this gradient solely affected the time interval between transitions, neither resulting in the loss of a state nor introducing new ones. These analyses indicate that global temporal regulators modulates the protraction or acceleration of lineage progression in its entirety.

In agreement with these computational analyses, a companion study (Zhang et al.) took a top-down experimental approach and offered empirical evidence: experimental removal of members of global temporal regulators (e.g. NFI family) extended the duration of each RGP stage and the corresponding production of stage-specific progenies (Zhang et al.). These changes were consistently observed throughout lineage progression in both a mouse model and a human cerebral organoid system (Zhang et al.). Conversely, overexpressing these regulators had the opposite effects.

In addition, our single-cell multiome data showed that NFI RNA expression and NFI motif accessibility increase in close synchrony along the RGP trajectory (**Figure 6b**).

Interestingly, NFIs are the most strongly enriched TFs in dynamic CREs across all transitions, with particularly prominent enrichment in opening CREs (**Figure 6j,k**), although they also appear in closing CREs. This pattern suggests that rising NFI levels may progressively increase chromatin accessibility, facilitating the activation of stage-specific regulatory programs.

Consistent with reports that NFIs exhibits chromatin opening ability in a context dependent manner^83–86^. This suggests that increasing NFI abundance may potentiate TF regulatory activity through dynamic chromatin accessibility, thereby coordinating state transitions.

Of note, searching for other common upstream factors highlighted miRNA and several signaling pathways (**Figure S6-3**), including recognized RGP regulators like let-7^87^ and the Wnt, Notch, and TGF-beta signaling pathways^79,88–90^, suggesting additional intrinsic and extrinsic regulators that fine-tune temporal progression. Taken together, these results suggest that neocortical development is regulated by a global timing program, featuring a transition hub regulatory network and global temporal regulators, which coordinate cascade progression (**Figure 6l**).

## Discussion

We aimed to elucidate the regulatory programs of neocortical development using the scAnR-seq co-assay. This assay outperformed other technologies, especially in detecting crucial regulatory elements like TFs and CREs. Notably, distal enhancers, despite being less accessible, were detected much more effectively (**Figure S2-3 f-h**). We found that genes often have multiple distal CREs. This suggests a more flexible expression modulation, highlighting the importance of distal CREs in developmental gene regulation^36,91^. As the field is anticipated to shift from high-throughput cell typing to in-depth characterization of gene expression programs, insights from these detailed analyses will be pivotal in deciphering transcriptional regulation dynamics and cell fate transition principles.

Our analytical framework focused on transition-wise dynamics, revealing transition-hub TFs key to neocortical development. Instead of adhering to temporal expression windows, these TFs form a cascade achieved through modulating their regulatory activity. This gives rise to a regulatory network with emergent, asynchronous temporality. Integral to this is the cross-regulation and chromatin modulation, ensuring appropriate gene regulation without TF on/off toggling, echoing the roles of PRC2 and NuRD in RGP patterning^92–97^ and findings in interneuron development^98^. Thus, unlike certain invertebrates using temporally restricted TFs, mammals seem to adopt an adaptable TF strategy throughout cell-state transitions. In support of this, large-scale single-cell mammalian studies show that distinct expression patterns are predominant in mature cells, not progenitors^99–103^. It is possible that initial broad gene expressions in progenitors refine into specific patterns during maturation, demarcating terminal cell types^9,27^.

Our study uncovers a network involving intricate global coordination governed by the increasing gradient of global temporal regulators like the NFI TF family. Experimentally altering these regulators could tune the overall span of lineage progression, distinguishing them from previously reported regulators that alter cell fate^8^, drop alternative lineage^104^, or generate aberrant populations^27,105^. Large-scale brain atlases reveal conserved cell types between mouse and human neocortex^12,13^, and the NFI family’s steeper expression in mice might explain size variances (**Figure S6-2 d**). These findings propose a developmental framework comprising the temporal network of transition-hub TFs and global temporal regulators, unifying flexibility in the control of tissue size and stability in the control of cellular composition. Lastly, our study underscores the potential of global temporal regulators in bidirectionally probing temporal patterns across diverse biological contexts.

## Limitations of the study

We acknowledge the following limitations in the current study. First, determining the activity of a regulatory link requires accurate measurement of statistical covariance between a TF and the target gene. This approach may not be suitable for assessing whether a TF actively maintains the expression level of a target gene when both the TF and the target gene show little expression change. In such cases, direct analysis of TF regulation is further complicated by other potential acting regulations, including DNA methylation, histone modifications, and co-factors that can induce epigenetic landscapes responsible for maintaining stable expression levels. Although we utilized single-cell local residual to confirm the strong activity of regulatory links, the resolution of this analysis may be limited for low-level variabilities due to measurement noise in single-cell omics. Further improvement in single-cell sequencing technologies, in combination with perturbations that introduce synthetic variations in gene expression, will be necessary to more accurately elucidate regulatory activity involved in the maintenance of gene expression levels. Second, our study mainly focuses on profiling gene expression and chromatin accessibility and therefore misses out on orthogonal molecular information. For instance, comprehensive TF ChIP-seq experiments will help distinguish binding events by TFs that have high similarity in cognate motifs. High-resolution Hi-C data in time series will further support the characterization of CRE activity dynamics. Future studies that incorporate measurements of protein abundance, TF binding, histone modifications, and DNA methylation in the same single cells will greatly enhance our understanding of transcriptional and post-transcriptional regulation during development.

## Acknowledgements

We thank all members of the Y.L. laboratory for discussions and support, W. Yao in W. Xie lab for technical support on ChIP-seq, Y. Liu, J. Fang in the Technology Center for Protein Sciences at Tsinghua University for assistance with cells sorting and for sequencing library preparation, Drs. J. Gu, C. David, M. Chen, and D. Mi for helpful comments on the manuscript. The work in the laboratory of Y.L. was supported by the National Key R&D Program of China 2019YFA0904402, Beijing Natural Science Foundation Z210010, the National Key R&D Program of China 2019YFA0906700, the National Natural Science Foundation of China 32171448, Tsinghua University Spring Breeze Fund 2020Z99CFG006, Tsinghua University Initiative Scientific Research Program 2021Z11JCQ020, 2022Z11QYJ032. L.D. was supported by the National Key R&D Program of China 2019YFA0906700, National Natural Science Foundation of China 31971513, 32061143023. S.-H.S. was supported by grants from the Ministry of Science and Technology of China (2021ZD0202300), the National Natural Science Foundation of China (32021002), Beijing Outstanding Young Scientist Program (BJJWZYJH01201910003012), Beijing Municipal Science & Technology Commission (Z20111000530000 and Z211100003321001), Chinese Institute for Brain Research (Beijing) and the New Cornerstone Investigator Program.

## Author Contributions

Y.L., G.Y., S.-H.S., Z.Z. provided overall project design. For scAnR-seq, G.Y., M.Z., Y.L. established protocol, X.D., X.Y. prepared single cell suspension, G.Y. generated sequencing library, G.Y., Z.Z., Y.L, N.Z., S.-H.S. analyzed data. Z.D., X.Z. collected the 10x multiome data from spinal cord samples. For ChIP-seq and Hi-C, X.Y., X.D. prepared sample, Z.Z. generated sequencing library, Z.Z., Y.L., J.Y., G.Y. processed data. For TF overexpression and knockdown, G.Y., X.Y., Z.Z. made constructs, X.D. performed in utero electroporation, immunohistochemistry, and imaging, X.D., G.Y., Y.L., S.-H.S., Z.Z. analyzed imaging data, G.Y., Z.Z. generated transcriptomics sequencing library, G.Y., Z.Z., Y.L., S.-H.S., Q.Z. analyzed transcriptomics data. For transition regulation analysis, Z.Z., G.Y., Y.L., S.-H.S., Q.Z. designed, and performed analysis. For dynamics simulation, X.W.W. performed the simulation, G.Y., Z.Z., X.W., Y.L., Y.Y.L., and L.D. analyzed the simulation results. Y.L., G.Y., Z.Z., X.W.W., Y.Y.L. wrote the manuscript with inputs provided by all authors.

## Competing interests

The authors declare no competing interests.

## Data Availability

Sequencing data for this study will be available through the Gene Expression Omnibus with the accession number GSE206105 (All scAnR-seq data) and GSE204701 (All Hi-C, Chip-seq, and CUT&Tag data).

## Code Availability

Code for analyses will be made available through Github at https://github.com/yuangh16/RGP-Project.

## Methods Cell culture

HEK293T cells were cultured in Dulbecco’s Modified Eagle Medium (DMEM, Gibco, C11995500BT) supplemented with 10% FBS (Gemini, 900-108), and 1% Penicillin-Streptomycin (Gibco, 10378016). K562 cells were cultured in RPMI-1640 medium (Gibco, C22400500BT) supplemented with 10% FBS, and 1% Penicillin-Streptomycin. All cells were cultured at 37 °C with 5% CO_2_ and passaged every two days.

## Animal care

All animal procedures were approved by the Institutional Animal Care and Use Committee (IACUC). CD-1 mice were purchased from Vital River Laboratory Animal Technology Co., Ltd. (Beijing, China) and maintained in Tsinghua University. Cas9 knock-in mice were kindly provided by Dr. Min Peng. Genotyping was carried out using standard PCR protocols. For experiments involving timed pregnancy, plug date was designated as E0.

## Single-cell suspension preparation

Single-cell suspension was prepared from E10 to E17 CD-1 mouse embryos. Four brains were collected for each time point. For E10 and E11 time points, ventricular zone was dissected from the tip of the brain. For E12 to E17 time points, brain tissues were first embedded in 4% low melting agarose at 35-37 °C and sectioned at 200 μm thickness with a vibratome (Leica) in ice-cold and oxygen-bubbling artificial cerebrospinal fluid (ACSF) containing: 125 mM NaCl, 5 mM KCl, 1.25 mM NaH_2_PO_4_, 1 mM MgSO_4_, 2 mM CaCl_2_, 25 mM NaHCO_3_, and 20 mM glucose (pH 7.4). The brain sections were transferred to a 3.5 cm dish containing ice-cold hibernate-E medium, and then ventricular zone was dissected and picked up under a stereoscope.

The isolated VZ tissues were digested with papain using a commercial kit (Worthington, LK003150) at 37 °C for 30 min according to the manufacturer’s recommendations. Single cell suspension was generated by dissociating the digested tissues through pipetting, washed with DPBS to remove digesting enzymes, and stained with DAPI in ice-cold DPBS. To enrich RGPs at later time points, FlashTag labeling was performed as previously described^45^. Briefly, pregnant mice were anesthetized, injected with FlashTag into the lateral ventricle of the uterine horns, and housed for 30 min. Then, single-cell suspensions were prepared as described above.

## scAnR-seq

### Preparation of coated beads

To prepare magnetic beads with binding specificity to nucleus, 5 µL Dynabeads^®^ M-280 Sheep Anti-Rabbit IgG beads (Thermo Fisher Scientific, 11203D) were washed once in PBS (Thermo Fisher Scientific, AM9624), resuspended in 50 µL Anti-Nesprin1 primary antibody (Abcam, ab192234) solutions (1:50 diluted in PBS), and incubated at room temperature for 30 min with rotation. The coated beads were washed once in PBS and resuspended in 250 µL lysis buffer, containing 10 mM HEPES-KOH (pH 7.5, Gibco, 15630080), 2.5 mM MgCl_2_ (Invitrogen, AM9530G), 10 mM KCl (Invitrogen, AM9640G), 2 U/µL RNase Inhibitor (Takara, 2313B), 1 mM DTT (Invitrogen, P2325), 0.5% Triton (Sigma-Aldrich, T8787-250ML).

### Single cell separation of nucleus and cytoplasm

Single live cells (DAPI negative) were sorted into 5 µL lysis buffer containing coated beads in batches of 96 well plates and centrifuged at 700 g for 1 min. The plates were vortexed to resuspend beads and incubated on a thermo mixer at 4 °C with shaking at 300 rpm for 30 min. After incubation, samples were placed on a magnet for 3 min to separate the nucleic and cytoplasmic content. Supernatant containing cytoplasmic content was carefully transferred to new plates for scRNA-seq, and the beads containing nuclei in the original plates were resuspended in 5 µL tagmentation mix for snATAC-seq.

### snATAC-seq for separated nuclei

The tagmentation mix was prepared as follows: 1 µL 5 × TTBL (Vazyme, TD501), 0.5 µL TTE mix V50 (Vazyme, TD501), 1.65 µL PBS, 0.05 µL Tween-20 (10%, Sigma-Aldrich, 11332465001), 0.025 µL Digitonin (2%, Promega, G9441), 0.5 µL DMF (Sigma-Aldrich, D4551-500ML), 1.275 µL H_2_O (Invitrogen, 10977023). The left nuclei on beads were resuspended by 5 µL tagmentation mix and incubated at 37 °C for 30 min. Then 5 µL stop mix containing 0.5 µL SDS (10%, Invitrogen, AM9820), 1 µL EDTA (500 mM, Invitrogen, AM9260G), 0.5 µL Proteinase K (20 μg/μL, New England Biolabs, P8107S), 0.04 µL carrier RNA (1 μg/μL, Thermo Fisher Scientific, 4382878), 2.96 µL H_2_O was added into each reaction and incubated at 55 °C for 30 min to terminate reactions. The reactions were purified with 2.2 × VAHTS DNA Clean Beads (Vazyme, N411-03). The eluted DNA were transferred into 25 μL PCR reactions containing 5 μL 5 × KAPA HiFi Fidelity Buffer (KAPA Biosystems, KK2102), 0.75 μL dNTP mix (10 mM each, KAPA Biosystems, KK2102), 0.5 μL KAPA HiFi DNA Polymerase (KAPA Biosystems, KK2102), 1.25 μL custom Nextera PCR primers 1 (25 mM; Table S5), and 1.25 μL custom Nextera PCR primers 2 (25 mM; Table S5). PCR was performed to amplify the library for 22 cycles for single cell using the following PCR program: 72 °C for 5 min; 98 °C for 3 min; and thermocycling at 98 °C for 20 s, 63 °C for 30 s and 72 °C for 3 min; following by 72 °C for 5 min. After the PCR reaction, libraries were purified with 1.8 × VAHTS DNA Clean Beads.

### scRNA-seq for separated cytoplasm content

The cytoplasmic RNA content in the transferred supernatant was purified by 2.2 × VAHTS RNA Clean Beads (Vazyme, N412-03) and used as input to a modified SMART-seq2 protocol for scRNA-seq library preparation^117,118^. Briefly, RNA was eluted into 4 µL elution mix containing 1 µL RT primer (10 µM), 1 µL dNTP mix (10 mM each, New England Biolabs, N0447L), 1 µL RNase Inhibitor (4 U/µL), and 1 µL H_2_O. Eluted samples were incubated at 72 °C for 3 min and immediately placed on ice. Each sample was added with 7 µL reverse transcription (RT) mix containing 0.75 µL H_2_O, 0.1 µL Maxima Reverse Transcriptase (Thermo Fisher Scientific, EP0741), 2 µL 5 × RT buffer (Thermo Fisher Scientific, EP0741), 2 µL Betaine (5 M, Sigma-Aldrich, B0300), 0.9 µL MgCl_2_ (100 mM), 1 µL TSO primer (10 µM), 0.25 µL RNase Inhibitor (40 U/µL). The RT reaction was incubated at 42 °C for 90 min, followed by 10 cycles of (50 °C for 2 min, 42 °C for 2 min), and heat inactivated at 70 °C for 15 min. Samples were then amplified with an addition of 14 µL PCR mix containing 1 µL H_2_O, 0.5 µL ISPCR primer (10 µM), 12.5 µL 2 × KAPA HiFi HotStart ReadyMix (KAPA Biosystems, KK2602). The PCR reaction was performed as follows: 98 °C for 3 min, 22 cycles of (98 °C for 15 sec, 67 °C for 20 sec, 72 °C for 6 min), and final extension at 72 °C for 5 min. The amplified cDNA product was purified using 0.8 × VAHTS DNA Clean Beads. Sequencing libraries were prepared using TruePrep DNA Library Prep Kit V2 for Illumina (Vazyme, TD503-02). Primer sequences: RT primer (Sangon), 5’Biotin-AAGCAGTGGTATCAACGCAGAGTACTTTTTTTTTTTTTTTTTTTTTTTTTTTTTTVN; TSO primer (Sangon), 5’Biotin-AAGCAGTGGTATCAACGCAGAGTACAT/rG//rG//iXNA_G/; ISPCR primer (Sangon), 5’Biotin-AAGCAGTGGTATCAACGCAGA*G*T.

### ChIP-seq and Library Preparation

ChIP-seq libraries were prepared as described previously^119^ with minor modifications: A total of 1 × 10^5^ FlashTag^+^ cortical cells were FACS-purified and pelleted immediately after sorting for each experiment. The pelleted cells were frozen in liquid nitrogen and stored at −80 °C until further use. Cells were resuspended in cold Nuclear Isolation Buffer (Sigma-Aldrich, NUC-101). Before immunoprecipitation (IP), cells were thawed and digested using 80 U MNase (New England Biolabs, M0247S) for 5 min at 37 °C. The digestion was stopped by adding 10% of the reaction volume of 100 mM EDTA (Invitrogen, AM9260G). Chromatin was precleared in complete immunoprecipitation buffer with 10 µL Protein A/G Dynabeads (Invitrogen, 10001D and 10003D) for 2 hours at 4 °C with rotation. Meanwhile, 10 µL Protein A/G Dynabeads were mixed with 2 µL H3K27me3 antibody (Cell Signaling Technology, 9733) or H3K27ac antibody (Abcam, ab4729) under rotation for 3 hours at 4 °C in 100 µL complete immunoprecipitation buffer. The precleared chromatin was mixed with the antibody-bound beads under overnight rotation at 4 °C. Beads were then washed twice with low salt wash buffer and twice with high salt wash buffer. DNA was then eluted for 1.5 hours at 65 °C in 30 µL ChIP elution buffer. DNA was purified using 2.2 × VAHTS DNA Clean Beads and eluted in ultrapure H_2_O (Thermo Fisher Scientific, 10977015). For H3K27ac IP, 10 mM sodium butyrate (Sigma-Aldrich, 303410) was added in cold Nuclear Isolation Buffer, complete immunoprecipitation buffer, low salt wash buffer, and high salt wash buffer. Libraries were prepared using NEBNext^®^ Ultra™ II DNA Library Prep Kit (New England Biolabs, E7645S), according to the manufacturer’s instructions. Libraries were PCR amplified for 13-15 cycles using NEBNext^®^ Multiplex Oligos (New England Biolabs, E7335S). The amplified libraries were purified using 2.2 × VAHTS DNA Clean Beads and eluted in ultrapure H_2_O.

### Hi-C library preparation

Hi-C libraries were prepared as described previously^120^ with minor modifications: 1 × 10^5^ FlashTag^+^ FACS-purified cortical cells were immediately fixed for 10 min in freshly prepared 1% formaldehyde (Thermo Fisher Scientific, 28906) in PBS (Thermo Fisher Scientific, AM9624) at room temperature with gentle rotation (300 rpm). Then the reaction was quenched in 0.2 M glycine (Sigma-Aldrich, 50046) solution for 5 min at room temperature with gentle rotation (300 rpm). The cells were washed twice with 1 ml cold 1 × PBS (centrifuged at 400 g for 5 min at 4 °C). To lyse cells, the cells were resuspended in ice-cold in situ Hi-C buffer and incubated on ice for 15 min. The lysed cells were frozen in liquid nitrogen and stored at −80 °C until further use. Chromatin was digested using 50 U of MboI (New England Biolabs, R0147S) overnight at 37 ℃ with gentle rotation. After biotin filling, proximity ligation was carried out using 17.5 U T4 DNA Ligase (Thermo Fisher Scientific, EL0012) for 4 hours at 20 °C with gentle rotation. DNA was sheared to 200-500 bp fragments using Covaris S220 sonicator with a duty factor of 10%, peak power of 140 W, and 200 cycles per burst. Each sample was sonicated for 2 cycles, 50 s/cycle. Ligation fragments containing biotin were immobilized on Dynabeads™ MyOne™ Streptavidin C1 beads (Thermo Fisher Scientific, 65001). Libraries were prepared using NEBNext^®^ Ultra™ II DNA Library Prep Kit, according to the manufacturer’s instructions. Libraries were PCR amplified for 14-16 cycles using NEBNext^®^ Multiplex Oligos. DNA was then purified using 2.2 × VAHTS DNA Clean Beads and eluted in ultrapure H_2_O.

### Library QC and sequencing

Before sequencing, libraries were quantified by qPCR, and the size distribution was assessed using Agilent 2100 Bioanalyzer. Libraries were then sequenced 150 bp paired-end run on the HiSeq X Ten (Illumina) or NovaSeq platforms (Illumina).

### Single-cell RNA-seq pre-processing

Raw fastq reads were trimmed by Cutadapt (version 1.18)^121^ to remove adapter and low-quality sequences. Reads were aligned to the mouse genome (mm10) using STAR (version 2.5.3)^122^. RSEM (version 1.3.0)^123^ with GENCODE gene annotation file (GRCm38.m23) was used to count gene expression.

### Single nucleus ATAC-seq pre-processing

Raw fastq reads were trimmed by Cutadapt (version 1.18) to remove adapter and low-quality sequences. Reads were aligned to the mouse genome (mm10) using Bowtie2 (version 2.3.3.1)^124^ with the following parameters: -t -q -N 1 -L 25 -X 2000 --no-mixed --no-discordant. Reads that have alignment quality < Q30, mapped to chrM, or overlapped with ENCODE blacklisted regions (https://sites.google.com/site/anshulkundaje/projects/blacklists) were discarded.

Duplicates were removed using Picard (version 2.20.4; http://broadinstitute.github.io/picard/) and SAMtools (version 1.13)^125^. Open chromatin region peaks were called on merged single cell and bulk ATAC-seq data for each time point using MACS2 peak caller (version 2.2.7.1)^126^ with the following parameters: -nomodel -nolambda -call-summits. Peaks from all samples were merged. Peak summits were extended by 250 bp on each side and used to define accessible regions^127^. Peaks outside ± 5 kb window from any TSS were defined as distal regulatory regions^128^.

### ChIP-seq data processing

Raw fastq reads were trimmed by Cutadapt (version 1.18) to remove adapter and low-quality sequences. Fastq files were aligned to the mm10 reference genome using Bowtie2 (version 2.3.3.1) with -X 2000 option. Reads that have alignment quality < Q30, improperly paired, or aligned to ENCODE blacklisted regions were discarded. Duplicates were removed using Picard (version 2.20.4) tools. ChIP-seq signal for each sample were normalized to Reads Per Kilobase per Million mapped reads (RPKM). Peaks were called on individual samples using MACS2 (version 2.2.7.1) with the following parameters: --nomodel –nolambda. Differential peaks were identified using “DESeq2” analysis in “DiffBind” package (version 2.14.0; https://bioconductor.org/packages/release/bioc/html/DiffBind.html) in R. For H3K27me3, differential peaks were identified by absolute log2 fold change > 0.4 and P value < 0.05. For H3K27ac, differential peaks were identified by absolute log2 fold change > 0.4 and P value < 0.01.

For E10-E12 H3K27me3 ChIP-seq, the average sequencing depth is 28.7M, and the average number of mapped reads is 17.4M. For E10-E12 H3K27ac ChIP-seq, the average sequencing depth is 27.7M, and the average number of mapped reads is 25.2M. All samples were sequenced at a depth comparable to previous studies, such as ENCODE standard. For example, the sequencing depth for E10 H3K27me3 ChIP-seq are 35.5M, 50.1M for each of two replicates, respectively. The sequencing depth for E10 H3K27ac ChIP-seq are 31.9M and 35.9M for each of two replicates, respectively. The number of mapped reads for E10 H3K27me3 ChIP-seq are 29.8M, 21.6M for each of two replicates respectively. The number of mapped reads for E10 H3K27ac ChIP-seq are 28.9M and 32.8M for each of two replicates, respectively.

For the later stages (E13-E15), when RGPs are heavily outnumbered by other cell types in the VZ, we obtained the profile of H3K27me3 and H3K27ac by CUT&Tag^129^. For E13-E15 H3K27me3 CUT&Tag, the average sequencing depth is 12.0M, and the average number of mapped reads is 11.3M. For E13-E15 H3K27ac CUT&Tag, the average sequencing depth is 32.2M, and the average number of mapped reads is 30.7M. All samples were sequenced at a depth comparable to previous studies^129^.

### Hi-C data processing

Raw sequencing reads were trimmed by Cutadapt (version 1.18) to remove adapter and lowquality sequences. Trimmed paired-end reads were aligned, processed and iteratively corrected using HiCPro (version 2.11.1)^130^ with default parameters. Sequencing reads were mapped to the mm10 reference genome. To recover chimeric fragments spanning ligation junctions, the ligation sequence (GATCGATC) was searched in unmapped fragments and was used as split site to separate Hi-C joined ends in the chimeric fragments. Biological replicates were pooled for each time point using HiCPro with “-s merge_persample”. Hi-C contact difference was calculated using “hicCompareMatrices” function in HiCExplorer with a 1-Mb, 40-kb or 10-kb bin size. Hi-C contact maps were visualized using “hicPlotMatrix” function in HiCExplorer (version 3.6)^131^ with a 1-Mb or 10-kb bin size. For CRE filtering (see section “CRE identification”), Hi-C data from all time points were merged using HiCPro with “-s merge_persample”. To call TADs from Hi-C contact matrix, R package spectralTAD (version 1.8.0)^132^ was used on 10-kb bin size with default parameters and kept level 1 TAD. Hi-C contact map with TAD was visualized using “hicPlotTADs” function in HiCExplorer with a 40-kb or 10-kb bin size. For E10-E15 Hi-C, the average sequencing depth is 184.1M, and the average number of mapped reads is 180.0M. All samples were sequenced at a depth comparable to previous studies^120^. For example, the sequencing depth for E10 Hi-C are 164.2M and 217.7M for each of two replicates, respectively. The number of mapped reads for E10 Hi-C are 156.6M and 213.8M for each of two replicates, respectively.

### Dimensionality reduction and cell clustering

#### Integrated ATAC and RNA

The weighted nearest neighbor (WNN) approach in Seurat v4^46^ was used to integrate the single cell ATAC and RNA information. WNN learned cell-specific “weights” for each modality, which reflects their information content and relative importance in downstream analyses. This enables the generation of a WNN graph: this graph denotes the most similar cells for each cell, based on a weighted combination of ATAC and RNA similarities. Here, we used the top 20 PCs in RNA and the top 20 PCs in ATAC as the input to “FindMultiModalNeighbors” function to generate the WNN graph. The WNN graph was then used as input to build uniform manifold approximation and projection (UMAP) with “RunUMAP” function with parameters: nn.name = “weighted.nn”. To identify cellular clusters, we used “FindClusters” function with parameters: graph.name = “wsnn”, algorithm = 3, resolution = 3.

#### RNA only

Dimension reduction and clustering analysis with RNA information were performed using the Seurat (version 4.0) bioinformatics pipeline. We first created a “Seurat object”, which includes all cells and all genes. To remove sequencing depth biases between cells, we normalized and scaled Transcripts Per Kilobase Million (TPM) counts using “NormalizeData” (normalization.method = “LogNormalize”, scale.factor = 1000000) combined with “ScaleData” function. We then determined the most variable genes using “FindVariableFeatures” function with following settings: selection.method = “vst”, nfeatures = 2000. Next, we identified statistically significant principal components in the top 2000^31^ highly variable genes and selected the top 19 principal components as input to UMAP dimensional reduction using the “RunUMAP” function. To identify cellular clusters, we used “FindClusters” function with parameters: resolution = 2.6.

#### ATAC only

Dimension reduction and clustering analysis with ATAC information were performed using ChromVAR (version 1.8.0)^49^ and Seurat (version 4.0) bioinformatics pipeline. First, ChromVAR was used to quantify single-cells accessibility variation by aggregating accessible regions containing a specific TF motif. Cells with fewer than 5,000 read pairs or less than 0.25 reads-in-peak ratio (fraction of reads in peaks, FRiP) were filtered from further analyses. To facilitate multi-modal analysis, the single-cells accessibility of each motif was then used as input to build a new assay in the “Seurat object”. We identified statistically significant principal components in highly variable motifs (variance > 1.5) using the “RunPCA” function. Dimensionality reduction was performed on the top 9 components of PCA using the “RunUMAP” function. To identify cellular clusters, we used “FindClusters” function with parameters: resolution = 1.

### Pseudo-time inference

The pseudo-time inference was performed with Slingshot (version 1.4.0)^133^ on all identified RGP cells. Briefly, a pseudo-time trajectory was computed for RGP cells on WNN-based UMAP projection using “getLineages” and “getCurves” functions in Slingshot. The left end of trajectory contained cells from E10 time point and was marked as the beginning of pseudo-time. For all cells, pseudo-time was calculated as the distance of a cell to the left end along the trajectory using “slingPseudotime” function.

### CRE identification

First, Cicero (version 1.4.4)^55^ was used to identify co-accessible peaks using the following parameters: cutoff 0.35 for co-accessibility score and *k* = 50, where k is the number of cells in k-nearest neighbors to aggregate prior to the calculation of co-accessibility scores. The co-accessibility score to each pair of peaks was assigned by Cicero using Graphical Lasso model with the penalty of genomic distance. The putative CRE was defined as the identified co-accessible peaks that located in the ± 500 kb window around each annotated TSS^55^. Next, Signac (version 1.10)^134^ function “LinkPeaks” was used to identify peak-gene associations in cis. To reduce the noise and sparsity of raw single cell data, aggregated fragments count matrix from Cicero was used as input for Signac. Signac computed Pearson correlation coefficient between RNA expression of a gene and the accessibility of peaks within ± 500 kb window of its TSS. The peak-gene links were defined by significant peak-genes associations (absolute correlation > 0.6, P < 0.05, one-sided z-test). Finally, we merged the CRE identified in Cicero and Signac for each gene. To obtain high-fidelity CRE, we further filtered CRE-gene association supported by topological contact (normalized Hi-C contact frequencies > 1) in merged Hi-C data from all time points. As co-accessibility was computed over all time points, we therefore aggregated Hi-C data from all time points.

### TF motif accessibility

First, FIMO (MEME suite v5.0.4)^135^ was used to identify TF motif occurrences in CRE using a first-order Markov background model, a P value cutoff of 10^-5^, and position weight matrices (PWMs) from the mouse HOCOMOCO motif database (v11)^136^. Then, we employed chromVAR to calculate TF-motif accessibility in each cell. Briefly, for each TF motif, we counted total fragments in all accessible peaks that were identified containing the motif occurrence by FIMO to generate a raw cell-motif count matrix. The cell-motif matrix was normalized by cell size factor using the “estimateSizeFactors” function in Cicero to correct for differences in read depth. Motif accessibility was then computed as z-score across all cells.

### Chromatin potential

Chromatin potential was calculated as previously described^36^. First, for gene *j* in cell *i*, we obtained its expression in RNA space (in TPM) and its expression in ATAC space, defined as averaged accessibility (in Fragments Per Kilobase Million (FPKM)) over distal and proximal CRE associated with gene *j*. Next, we calculated smoothed profiles of each cell, by averaging ATAC expressions and RNA expressions over its k-nearest neighbors (k-NN, *k* = 10). For each smoothed cell profile, we searched for its k-NN (*k* = 10) in smoothed RNA space (*XR*_*i,k*_) using its smoothed chromatin profile in ATAC space (*XA*_*i*_). This approach identifies the potential future RNA profiles of cell *i*, reflected by the average of *XR*_*i,k*_ over *k*, based on its current chromatin profile. Therefore, the direction of cell *i* to its k-NN defines the flow of the chromatin potential. To obtain a UMAP view of chromatin potential, we divided the UMAP space into a 20 × 20 grid and computed grid-representative arrows which were scaled to averaged chromatin potentials of all cells within each grid.

### Reference regulatory links

The reference regulatory links were obtained by two conventional methods: (1) we manually curated TF-target regulatory links that are supported by experimental evidence, obtaining 364 reference regulatory links (Table S2). (2) we selected 9 TFs that exhibited large variance in pseudo-time dynamics, overexpressed each TF in RGPs by in utero electroporation, and identified differentially expressed (DE) genes (P value < 0.05, absolute log2 fold change > 0.4), obtaining a total of 1,722 TF-DE gene pairs as reference regulatory links (Table S2). The details of plasmid preparation, in utero electroporation, library preparation, and data analysis are as follows:

### Plasmids and TF overexpression

For TF overexpression RNA-seq experiments, a *pCAG-IRES-tdTomato* plasmid backbone was selected which contains a CAG promoter for constitutive transcription and an internal ribosome entry sites (IRES) to express a reporter tdTomato on the same transcript. We amplified cDNA of *Lef1*, *Deaf1*, *E2f5*, *Xbp1*, *Lhx2*, *Hbp1*, *Mybl2*, *Hey1*, *Pbx2* and inserted each TF cDNA into *pCAG-IRES-tdTomato* right before IRES. In-utero electroporation was performed as previously described^137^. In brief, timed pregnant CD-1 mice were injected with 2 µg/µl plasmid DNA mixed with Fast green (Sigma-Aldrich, F7252) into the lateral ventricle, and five 50 ms pulses of 30-40 mV with a 950 ms interval were delivered across the uterus with two 9 mm electrode paddles positioned on either side of the head (BTX, ECM830).

### Bulk RNA-seq library preparation for TF overexpression experiments

For each overexpression experiment, batches of 100 tdTomato and FlashTag double-positive cells were sorted into 10 µL lysis buffer, containing TCL lysis buffer (Qiagen, 1031576) added with 10% 2-Mercaptoethanol (Sigma-Aldrich, M6250-100ML), and then centrifuged by 700 g for 1 min. The tdTomato was a marker for TF expression, and the FlashTag was used to enrich RGP cells^45^. The RNA was purified from lysed cells by 2.2 × VAHTS RNA Clean Beads and used as input to a modified SMART-seq2 protocol for scRNA-seq library preparation, as described in the section “scAnR-seq, scRNA-seq for separated cytoplasm content”.

### Processing of overexpression bulk RNA-seq

Raw fastq reads were trimmed by Cutadapt (version 1.18) to remove adapter and low-quality sequences. To enable the alignment of the tdTomato sequence in plasmid, the reference genome (mm10) and gene annotation files (GRCm38.m23) were modified with the addition of tdTomato sequence and annotation. Reads were aligned to the modified mouse genome using STAR (version 2.5.3). RSEM (version 1.3.0) with the modified GENCODE gene annotation file was used to count gene expression. Differential expression was estimated using edgeR package (version 3.28.1)^138^, and genes with absolute log2 fold change > 0.4 and P value < 0.05 driven by TF overexpression were included to build reference regulatory links.

### Multi-modal inference methodology

#### Generation of prior matrix

A prior matrix is a #genes by #TF binary matrix where the matrix entry (*i, j*) denotes whether there is an occurrence of the motif of TF *j* in CREs associated with the target gene *i*. The prior matrix represents the potential regulatory relation between TFs and target genes on the basis of putative TF binding on target CRE. Specifically, the prior matrix *P* is defined as

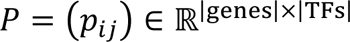

where *P*_*i*_ = 1 if the motif of TF *j* is identified in the CRE region of the target gene *i*, otherwise *P*_*i*_ = 0. We used FIMO (MEME suite v5.0.4) to search for TF motif occurrences in CREs, with the following settings: a first-order Markov background model, a P-value cutoff of 10^-5^, and PWMs from the mouse HOCOMOCO motif database (v11).

#### Regulatory links inference

We adopted the LASSO-StARS framework^60^, which regularizes false positive discoveries using LASSO fitting, to infer TF-target regulatory links:

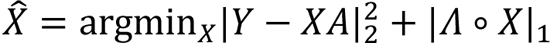

Here, the output *X* ∈ ℝ^|genes|×|TFs|^ is the statistical significance matrix of TF-target regulatory links to be inferred. The inputs are as follows:

*Y* ∈ ℝ^|genes|×|samples|^ is the expression matrix of target genes*. Y* takes one of two forms: (1) levels of gene expression and (2) change rates of the gene expression levels. Each form then takes one of the following transformations: (1) raw scale, (2) z-score, (3) logistic regression. The (2) and (3) were employed to stabilize variance. The parameter for logistic transformation was identified to be 1.0. This resulted in a total of 6 combinations of forms and transformations, which were designed to capture gene expression dynamics that exhibit diverse scales of variability.

*A* ∈ ℝ^|TFs|×|samples|^is the TF activity matrix. *A* is estimated by adapting published methods^60^. Briefly, *A* takes one of three forms: (1) TF expression level, (2) TF motif accessibility, and (3) estimated TF activity based on the expression level of known TF target genes^139^. For (1), *A* takes the expression matrix of all TFs, and then takes variance stabilizing transformations similar to that for *Y.* For (2), *A* takes the TF motif accessibility matrix generated in the section “TF motif accessibility”, and then takes variance stabilizing transformations similar to that for *Y*. For (3), *A* is obtained by solving *T = PA*, where *P* is the prior matrix, and *T* is the expression matrix of known TF target genes.

Λ ∈ ℝ^|genes|×|TFs|^ is a matrix of nonnegative penalties, and ∘ denotes Hadamard product in Λ ∘ *X*. Each element ΛΛ*_i,j_* in ΛΛ determines the strength of penalty for inferring a regulatory link between TF *j* and gene *i* on the basis of whether a prior evidence of TF *j* motif is identified in the CREs of the gene *i*. Briefly, Λ_*i,i*_ = *b* × λ if *P*_*i*_ = 1, and Λ_*i,i*_ = λ if *P*_*i*_ = 0 where, λ sets the penalty: the larger the value is, the stronger a penalty is applied. An optimal λ is searched by a data-driven selection method^60^. *b* ∈ (0.1, 0.25, 0.5, 1.0) sets the bias that weights the prior evidence during fitting: when *b* = 0.1, a stronger penalty is differentially placed on the links without prior evidence. Of note, previous studies have found that some TF may regulate target genes that contain no cognate binding motif of those TFs^60^, thus *b* > 0 is to permit a more flexible inference of regulatory relation. The Λ matrix is employed by the LASSO-StARS framework to reduce overfitting for regulatory links without prior evidence.

In total, 64 combinations of *Y*, *A*, Λ were obtained to estimate the statistical strength of regulatory links between TFs and target genes that exhibit diverse scales of variability and evidence of cognate binding motifs.

### Consolidation of regulatory links

We consolidated regulatory links inferred in the 64 combinations of *Y*, *A*, Λ into an overall regulatory relation. The consolidation procedure was derived from boosting procedure used in the machine learning community. We first divided reference TF-target regulatory links into training and testing datasets in a 4:1 ratio. Then, we consolidate regulatory links in mini-batches: (1) We initialized an empty output matrix *X*_0_ that does not contain any TF-target regulatory links. X_0_ denotes the overall regulatory relation. (2) In each iteration *t* (*t* > 0), we selected the top 1,000 regulatory links in each individual *X*_*i*_ (*i* = 1, ⋯,64) (not present in *X*_*t*−1_) based on their statistical strengths and added them into *X*_*t*−1_ to yield 64 new *X*^∽^_*i,t*_(*i* = 1, ⋯,64). Then, we calculated the number of overlapped regulatory links between the training dataset and regulatory links in each *X*^∽^_*i,t*_ and updated *X*_*t*_ as the *X*^∽^_*i,t*_ with the highest overlapped number. (3) Repeat step-(2) until the overlapped number is converged. (4) As the reference TF-target regulatory links comprised of both direct and indirect regulatory links, we also considered the indirect links in the consolidation process. To achieve this, we first obtained *X*_0_ as above. Then, we performed a second round of iterations, and in steps (2) and (3) of each iteration *t*, *X*_*t*_ was updated as the *X*^∽^_*i,t*_ with the highest number of overlapped regulatory links between the training dataset and the indirect regulatory links in each *X*^∽^_*i,t*_.

### Evaluation of the overall regulatory relation

To examine the overall regulatory relation in the consolidated network, we performed the following two evaluations: (1) we calculated the percentage of reference regulatory links in the test dataset that were consolidated in the overall regulatory relation. The regulation relation with high performance should capture a high proportion of reference regulatory links in the test dataset. (2) As the predicted target genes in high-performance regulation relation should be enriched in the differentially expressed (DE) genes due to TF over-expressions, we accordingly calculated the enrichment score (P value using Fisher’s exact test) between TF-DE gene pairs and overall regulatory relation using the “fisher.test” function in R. The TF-DE gene pairs were obtained as described in the section “Reference regulatory links, Processing of overexpression bulk RNA-seq”. Briefly, differential expression was estimated using the edgeR package (version 3.28.1), and genes with absolute log2 fold change > 0.25 and P value < 0.05 driven by TF overexpression were used as TF-DE gene pairs. TF-DE gene pairs in the training dataset were excluded from the enrichment calculation.

### Alternative inference methods

To compare the performance of Multi-modal inference and alternative inference methods, we selected the following gene-regulatory approaches that are widely used. (1) RNA - (∂ RNA) / (∂ t) (Lasso): This approach infers regulatory links by fitting RNA expression data (response variable) given temporal differential RNA expression (explanatory variable) using LASSO fitting. We used only RNA expression data as input and applied the above inference method (see the section “Multi-modal inference methodology, Regulatory links Inference”) to infer regulatory links. (2) RNA - (∂ RNA) / (∂ t): This approach infers regulatory links by fitting RNA expression data (response variable) given temporal differential RNA expression (explanatory variable) using regression trees. We used the dynGEINE3^62^ algorithm with RNA expression data as input and with default parameters. (3) RNA+motif - RNA: This approach first obtains putative TF-target regulatory links by searching TF motif in CREs of target gene, and subsequently fitting target gene RNA expression data (response variable) given TF RNA expression data (explanatory variable) using Spearman correlation analysis. The motif scan in CREs of genes was done using FIMO (MEME suite v5.0.4) with the following settings: a first-order Markov background model, a P value cutoff of 10^-5^, and PWMs from the mouse HOCOMOCO motif database (v11). A TF-target regulatory link was identified if the TF motif was identified in CRE of the gene and correlation between the TF expression and the gene expression was higher than 0.5. (4) RNA - RNA: This approach infers regulatory links by fitting target gene RNA expression data (response variable) given TF RNA expression data (explanatory variable) using correlation network analysis. We used weighted correlation network analysis (WGCNA, version 1.69)^64^ algorithm to infer regulatory links with RNA expression as input and with default parameters.

### Regulatory links inference on co-variance permutated data

To assess the reliability of Multi-modal inference approach, we inferred and consolidated regulatory links based on co-variance permutated data. We first permutated samples for each gene, so that the co-variance in the gene expression matrix was largely diminished after permutation. Then for each dataset, we applied Multi-modal inference approach (see section “Multi-modal inference methodology, Regulatory links Inference”).

### PCA and NMF module analysis

The PCA module analysis was performed using the “pca” function in Matlab. The highly-variable modules were selected based on the variance explained (> 1% variance of PC1). The NMF module analysis was performed using the “nmf” function of NMF package (version 0.23.0)^140^ in R, with options: method = “snmf/r”, seed = “nndsvd”. The module number in NMF analysis was searched with the number of highly-variable PCA modules as guidance to cover the major expression dynamics on the basis of the variance explained.

### Short-time Fourier transform

Short-time Fourier transform (STFT) was used to examine how time-localized frequency varies over time. We computed STFT for each PCA module using the MATLAB function “spectrogram” across a total 361 (real data) or 500 (simulation data) samples along pseudo-time with options: hamming window, window length = 80 samples. Then we obtained matrices of the power spectrum by time for each PCA module. To identify temporal changes in the power spectrum, we computed the delta power along the time axis, which is defined as the difference in the spectrum between adjacent time points. The sudden changes in the power of high-frequency components align with major changes in the temporal dynamics of PCA modules. We max pooled delta power of all PCA modules to visualize temporal dynamics in a single plot.

### Activity of TF-target regulatory links

To assess the activity of TF-target regulatory links over the course of multiple transitions, we first divided pseudo-time into 17 consecutive time stages using sliding windows along pseudo-time. Then, the activity of each regulatory link was quantified as the strength of concordant changes in the linked target and TF within each time stage, which was defined as:

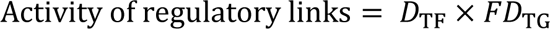

where *D*_TF_ represents the change of motif accessibility and expression for each TF in each time stage, and *FD*_TG_ is absolute log2 fold change of gene expression between the first and the second half of a time stage. *D*_TF_ = 1, if expression or motif accessibility of a TF was differential in a time stage. Otherwise, *D*_TF_ = 0.1. to represent low level of dynamics. Differential motif accessibility or differential expression was analyzed using Wilcoxon rank-sum test and identified by P value < 0.05, absolute log2 fold change > 0.65 (or absolute log2 fold change > 0.25).

Finally, the activity of regulatory links was normalized by z-score transformation. Regulatory links with activity higher than 2 were determined as active regulatory links (regulatory links with strong activity). To identify clusters of regulatory links according to their activity, we first identified statistically significant principal components using “RunPCA” and then selected all principal components for cluster analysis. The resolution of “FindClusters” function was set as 0.4. Next, the temporal boundaries that differentiate activities of regulatory links of neighboring clusters were obtained by maximizing likelihood ratio, e.g., using the function of “hclust” in R, and were used to partition stages along pseudo-time.

### Single-cell local residual analysis

We first obtained the local residuals of each gene by regressing single-cell gene expression against pseudo-time using the “RegressOutMatrix” function in the Seurat package (version 4.0)^46^. The raw single-cell gene expression is known to be corrupted by high false zeros, often termed dropout noise. To compensate for dropout noise in raw single-cell data, we computed de-noised local residuals for each gene in each single cell by smoothing the residuals over k-NN (*k* = 10) cells. To evaluate temporal coupling between a TF and gene pair, we calculated the Pearson correlation coefficient between the local residuals in TF and in the target gene in all time stages.

### Simulation with non-temporal or temporal regulatory links

#### Generation of synthetic transcription regulation network (TRN)

In our simulations, we generated TRN as follows: (1) the degree of each gene, i.e., number of its regulating TFs, is drawn from a Poisson distribution Pois(δ). (2) The probability of each TF to be selected as the regulator of a gene is proportional to: *p*(*j*) ∝ *i*^−τ^, then the degree distribution of TFs follows a scale-free distribution^141^: *P*(*k*) ∝ *k*^−(1+1/τ)^. The number of genes is *M* = 1,000 for all simulations, and the first 100 genes were set as TFs, = 100.

#### Dynamic model

Here, we modeled the dynamics of gene expressions using the Chemical Langevin equation (CLE)^142,143^, which has been employed in the GeneNetWeaver (GNW)^144,145^ and single-cell expression simulator (SERGIO)^71^. The expression time-course of gene-*ii* is modeled by the following ODE:

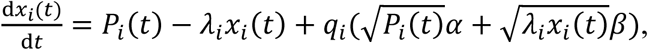

where *x*_*i*_(*tt*) is the expression of gene-*i* at time *t*. The regulators of gene-*i* are encoded in the corresponding TRN, and their influence on gene-*i* is reflected by the production rate *P*_*i*_(*t*). λ_*i*_ is the decay rate, and *q*_*i*_ is the noise amplitude in the transcription of gene-*i*. α and β are two parameters characterizing two independent Gaussian white noise processes.

The production rate *P*_*i*_(*t*) is expressed as the sum of contributions from each of its regulators:

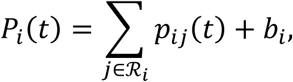

where ℛ_*i*_ is the set of all regulators of gene-*i*, and *b*_*i*_ is the basal production rate of gene-*i*. *p*_*i*_(*t*) measures the regulatory effect of gene-*j* on gene-*i*, which is modeled as a Hill function in terms of the expression of regulator gene-*j* (which is a TF):

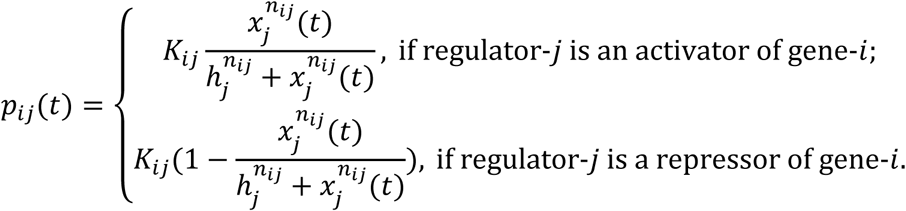

Here, *x*_*i*_(*t*) is the expression of regulator-*j* at time *t*, *K*_*i*_ is the maximum contribution of regulator-*j* to target gene-*i*, *n*_*i*_ is the Hill coefficient, and ℎ_*i*_ is the regulator expression that produces the half-maximum regulatory effect, respectively.

#### Parameterization

In all the simulations, we set λ_*i*_ = 0.8, *b*_*i*_ = 0, α = 0 and β = 0 to study the dynamics of stage transition. We set the Hill coefficient *n*_*i*_ = 2 and drew ℎ_*i*_ from the normal distribution of mean 2 and standard deviation 4: *N*(2,4) bounded between 1 and 10. We drew *K*_*i*_ from the uniform distribution *u*(−1,1). The initial expression of each gene was drawn from the uniform distribution *u*(0,1), and the ODE was integrated using Euler’s method with time step *dt* = 0.01, and the total simulation time *t* = 6,000.

#### Generation of simulation data

1. To simulate the gene expression data with non-temporal regulatory links, we used one global TRN and simulated for 5,000 steps. Specifically, a stimulus was added to drive the TFs with the highest degree (*L* = 20) in the global TRN. We included an initial 1,000 simulation steps with the stimulus set to 0 to allow the gene expressions to reach steady states. After that, we simulated gene expression profiles with a sustained stimulus with expression as 1. (2) To simulate the gene expression data with temporal regulatory links, we used five stage-TRNs extracted from the global TRN and simulated 1,000 steps for each stage. Specifically, a stimulus was added to drive TFs with the highest degree (*L* = 20) in the TRN of each stage in a continuous simulation, where the expression of each gene in the last step of a stage was the initial expression in the first step of the next stage.

### Transition-wise analysis of regulatory links

To perform the transition-wise analysis of regulatory links, we first divided pseudo-time into 6 cell states, which we denoted as E10, E11, E12, E13, E14, E15, as the cell states along pseudo-time were highly correlated with the RGP temporal identity and sampling. Accordingly, the 5 cell-state transitions were denoted as E10-E11, E11-E12, E12-E13, E13-E14, E14-E15. To identify the active regulatory links in each transition, we determined TFs and target genes that displayed salient changes (differential motif accessibility or differential expression) during each transition. These salient TFs were identified as the TFs showing significant differential motif accessibility or differential expression between the first and the second half of a transition, which were identified by Wilcoxon rank-sum test, P value < 0.05, absolute log2 fold change > 0.65 for motif accessibility or absolute log2 fold change > 0.25 for differential expression. The salient target genes were identified as genes displaying significant differential expression between the first and the second half of a transition, which were identified by Wilcoxon rank-sum test, P value < 0.05, absolute log2 fold change > 0.25. The regulatory links between these salient TFs and target genes in the overall regulatory relation comprise the active subset of regulatory links in each transition.

### Randomly shuffled overall regulatory relation

To shuffle overall regulatory relation, we randomly removed TF-target regulatory links and added back the same number of random TF-target regulatory links. The percentage of random removal and replacement ranged from 5% to 40% to systematically evaluate the robustness of transition hub structure or upstream TFs.

### Identification of transition hub TFs

Transition hub TFs were defined as those TFs with significantly higher number of regulatory links, referred to as degree, in the original regulation relation than that in random networks. To identify transition hub TFs, we employed permutation test for each TF in each transition. We first calculated the number of regulatory links for each TF in active subset from the original regulation relation and from 20 regulation relation in random networks. The generation of random networks was similar as described in the section “Regulatory links inference on co-variance permutated data”. Then, we randomly shuffled regulatory links to perturb scale-free distribution in the generated network. Next, for each TF, we calculated the statistical significance of its degree in the distribution of degrees of all TFs in each permutation using one-sample Student’s t-test. The transition hub TFs were identified as TFs with P values < 0.05 and FDR < 0.25, which means that TFs had P < 0.05 in at least 150/200 permutations. We further sorted these transition-hub TFs based on their degree, and the top 25 transition-hub TFs were referred to as top hub TFs in a transition. For transitions with less than 25 transition hub TFs, top hub TFs refer only to the transition-hub TFs.

### Biological process enrichment analysis

We performed functional enrichment analysis using the “clusterProfiler” R package (v3.14.3)^146^ with default parameters (P value < 0.05). The functional annotations were obtained from Kyoto Encyclopedia of Genes and Genomes (KEGG) pathway^147^ and terms in Gene Ontology (GO)^148^.

### Enrichment analysis of target genes in TAD

We obtained TADs in Hi-C data merged from all time points, as described in the section of “Hi-C data processing”. For each TF, we employed permutation test to calculate the enrichment score of its targets in TAD. Specifically, we generated a background distribution by randomly shuffling the position of TADs in the genome 200 times. Then, the enrichment score of targets clustering in TAD was calculated using one-sample Wilcoxon rank-sum test to quantify the statistical significance of number of target genes in real TAD in the distribution of numbers of target genes in randomly shuffled TAD.

### Gene set enrichment analysis (GSEA)

GSEA algorithm was used as described^149^. We performed GSEA analysis to calculate enrichment score and P value using “GSEA” function from “clusterProfiler” package (v3.14.3) in R. Specifically, the enrichment score represents difference between the observed rankings and expected rankings under a random walk distribution, as described^149^. The statistical significance level of the enrichment score is determined by the P value in Kolmogorov-Smirnov test.

### Luciferase reporter assay

#### Plasmids and cell transfection

To validate the regulation by CREs and their cognate TFs, luciferase reporter assay was performed as described previously^150,151^ with minor modifications. To construct plasmids for functional analysis of CREs, a *pGL4.23* (Promega, E8411) was selected as plasmid backbone which contains a mini promoter to express firefly luciferase gene Luc2. The CRE regions were amplified from mouse genomic DNA and inserted into the *pGL4.23* plasmid backbone right before the mini promoter. To express the cognate TFs, a *pCAG-IRES-tdTomato* plasmid backbone was selected which contains a CAG promoter for constitutive transcription and an internal ribosome entry sites (IRES) to express tdTomato on the same transcript. The cognate TFs were amplified from mouse genomic DNA and inserted into *pCAG-IRES-tdTomato* before IRES. HEK293T cells were cultured in 24-well plates and transfected with 240 ng *pGL4.23-empty* or pGL4.23-CRE, 240 ng *pCAG-IRES-EGFP* or *pCAG-TF-IRES-EGFP* and 40 ng *pGL4.73* (Promega, E6911), which encodes the expression of Renilla luciferase gene (hRluc) as an internal control for normalization. Cell transfection was performed as follows. First, HEK293T cells were seeded into 24-well plates (Corning, 3524) one day prior to transfection at a density of 200,000 cells per well. Cells were transfected using polyethylenimine (PEI; Polysciences, 24765-1). For each well of a 24-well plate, 3 µL PEI (1 mg/mL) or 500 ng plasmids was separately mixed with 24 µL OptiMEM (Thermo Fisher Scientific, 31985062) and incubated for 10 min at room temperature. Then, we combined the two mixtures and incubated mix for another 25 min at room temperature to prepare the transfection mix. Transfection mix was added dropwise to each well, and then the plate was shaken gently. Cells were cultured as described in the section “Cell culture”.

#### quantitative PCR (qPCR)

Expression of Luc2 and hRluc was quantified using qPCR. Cells were collected at 48 hours post transfection and then the total RNA was extracted using HiPure Total RNA Plus Kit (Magen, R411103). The total RNA was reverse transcribed using HiScript II Q RT SuperMix for qPCR (+gDNA wiper) (Vazyme, R223-01) to generate cDNA. qPCR was subsequently performed on the cDNA using NovoStart^®^SYBR qPCR SuperMix plus (Novoprotein, E096-01B) with gene specific qPCR primers (Table S5). The quantified levels of Luc2 transcripts were normalized to hRluc.

### Single-cell transcriptomics and phenotype analysis of *Hbp1* and *Foxk1*

#### Plasmids and in utero electroporation for Hbp1 and Foxk1 in single-cell transcriptome analysis

*pCAG-IRES-tdTomato, pCAG-Foxk1-IRES-tdTomato*, and *pCAG-Hbp1-IRES-tdTomato* plasmids were constructed for TF overexpression. The plasmid backbone and the procedure for in utero electroporation was the same as described in the section “Reference regulations, Plasmids and TF overexpression”.

#### Single cell RNA-seq library preparation

The library preparation for *Hbp1* or *Foxk1* overexpression scRNA-seq experiment was similar as described in the section “Reference regulations, Bulk RNA-seq library preparation for TF overexpression experiments” with the modification: tdTomato and FlashTag double-positive single cell was sorted into 5 µL lysis buffer.

#### Single cell RNA-seq processing

The processing of raw fastq reads, alignment, and reads counting was the same as described in the section “Reference regulations, Processing of overexpression bulk RNA-seq”. Dimension reduction and clustering analysis was performed similarly as described in the section “Dimensionality reduction and cell clustering, RNA only”. For cluster analysis in *Foxk1* overexpression data, the resolution of “FindClusters” function was set as 2.5. For cluster analysis in *Hbp1* overexpression data, the resolution of “FindClusters” function was set as 1.5. The RGP cell clusters were identified by the positive expression of known RGP markers *Pax6* and *Sox2*^9^. The RGP cell clusters whose majority were *Foxk1* or *Hbp1* overexpressing cells were denoted as RGP_*Foxk1* or RGP_*Hbp1* respectively, and the other RGP cell cluster was denoted as the control cluster of RGP. Differentially expressed genes between clusters were identified using the “FindMarkers” of Seurat with options: P value < 0.05 and absolute average log2 fold change > 0.25.

#### Plasmids and in utero electroporation for Foxk1 in phenotype analysis

For overexpression experiments, *pCAG-IRES-EGFP* and *pCAG-Foxk1-IRES-EGFP* plasmids were constructed. The plasmid backbone was similar as described in the section “Reference regulations, Plasmids and TF overexpression”, except that EGFP was used as reporter to facilitate imaging. For knockdown experiments, *pLL3.7-EGFP*, *pLL3.7-shFoxk1-EGFP*, and *pLL3.7-shFoxk2-EGFP* plasmids were constructed. The pLL3.7 plasmid contains an mU6 promoter for the expression of inserted shRNA and a separate CMV promoter that drives the expression of EGFP. shRNA sequences against *Foxk1* or *Foxk2* were designed as follows: *Foxk1* shRNA (ATGAAAACTACAGAAGCC), *Foxk2* shRNA (GTGACTCGAGTGGCCTGGC). All sense and anti-sense oligos were purchased from BGI (Shen Zhen, China), and annealed oligos were cloned into the HpaI and XhoI sites of the *pLL3.7*^152^. The procedure for in utero electroporation was the same as described in the section “Reference regulations, Plasmids and TF overexpression”.

### Immunohistochemistry and imaging

Embryonic mice were perfused with PBS (pH 7.4), followed by 4% paraformaldehyde (PFA) in PBS, and brains were post-fixed with 4% PFA for 6 hours at 4 °C. Coronal sections with 40 µm thickness were obtained using a vibratome (Leica, CM3050) and subjected to immunohistochemistry. Brain sections were blocked for 1 hour in 10% donkey serum containing 0.5% Triton X-100 at room temperature. Primary antibody incubation was subsequently performed for 48 hours at 4 °C, and then washed three times with PBS + 0.1% triton X-100, followed by secondary antibody incubation at room temperature for 2 hours. The following primary antibodies were used: chicken anti-GFP (Aves Labs, GFP-1020; 1:500 dilution), rabbit anti-PAX6 (BioLegend, 901301; 1:500 dilution), sheep anti-PAX6 (R&D Systems, AF8150; 1:500 dilution), mouse anti-TUBB III (Sigma-Aldrich, T8660; 1:500 dilution), rabbit anti-Ki67 (Abcam, AB16667; 1:100 dilution), rabbit anti-N-cadherin (BD bioscience, 610920; 1:100 dilution), rabbit anti-c-JUN (Cell Signaling, 9165; 1:500 dilution), rabbit anti-TCF4 (Abcam, ab217668; 1:500 dilution), mouse anti-SOX2 (Proteintech, 66411-1-Ig; 1:500 dilution). The following secondary antibodies were used: donkey anti-chicken IgY (H + L) 488 (Jackson ImmunoResearch, 703-546-155; 1:500 dilution), donkey anti-mouse IgG (H + L) 555 (Thermo Fisher Scientific, A-31570; 1:500 dilution), donkey anti-rabbit IgG (H + L) 647 (Thermo Fisher Scientific, A-31573; 1:500 dilution), donkey anti-Sheep IgG (H + L) 488 (Thermo Fisher Scientific, A-11015; 1:500 dilution), donkey anti-Sheep IgG (H + L) 647 (Thermo Fisher Scientific, A-21448; 1:500 dilution), donkey anti-mouse IgG (H + L) 488 (Thermo Fisher Scientific, A-21202; 1:500 dilution), donkey anti-rabbit IgG (H + L) 555 (Thermo Fisher Scientific, A-21432; 1:500 dilution). Nuclei were counterstained with DAPI (Thermo Fisher Scientific, D1306; 1:1,000 dilution). Images were obtained as z-stacks using a confocal microscope (Olympus, FV3000) with 20X air objective lens. Z-stack images were max projected and stitched using Fluoview (version 4.2, Olympus). Imaris (version 9.0.1, Oxford Instruments) was used to manually count cells. ImageJ (Fiji) (1.52p, NIH) was used to quantify fluorescence intensity. The figures of typical areas were prepared in ImageJ (Fiji) (1.52p, NIH) or Photoshop (Adobe).

### Knockdown efficiency evaluation in HEK293T

#### Cell transfection

To evaluate the knockdown efficiency of *Foxk1* shRNA in HEK293T cells, the shRNA plasmid (*pLL3.7-shFoxk1-EGFP*) or scrambled control plasmid (*pLL3.7-EGFP*) was co-transfected with the TF overexpression plasmid (*pCAG-Foxk1-IRES-EGFP*) at a 1:1 ratio. To evaluate the knockdown efficiency of *Foxk2* shRNA in HEK293T, shRNA plasmid (*pLL3.7-shFoxk2-EGFP)* or scrambled control plasmid (*pLL3.7-EGFP*) was co-transfected with TF overexpression plasmid (*pCAG-Foxk2-HA-IRES-EGFP*) at a 1:1 ratio. The *pCAG-Foxk2-HA-IRES-EGFP* plasmid overexpresses FOXK2 fused with an HA tag and was constructed as described in the section “Single-cell transcriptomics and functional analysis of *Hbp1* and *Foxk1*, Plasmids and in utero electroporation for *Foxk1* in functional analysis”. Cell transfection was performed as described in the section “Luciferase reporter assay, Plasmids and cell transfection”.

#### Western blot

Cells were collected with 1 × sample loading buffer (Mei5bio, MF145-10) at 48 hours after transfection, vortexed for 10 s, and boiled for 20 min at 95 °C. Whole-cell lysate was separated by SDS-PAGE electrophoresis and then transferred to a PVDF membrane (Bio-Rad, 1620177) under 100V for 100 min on ice. The membrane was blocked in 5% nonfat milk for 1 hour and subsequently immunoblotted with primary antibodies overnight at 4 °C. The following primary antibodies were used in this study: rabbit polyclonal anti-FOXK1 antibody (Sino Biological, 101153-T10; 1:1,000 dilution), mouse monoclonal anti-HA antibody (BioLegend, #MMS-101P; 1:2,000 dilution) and mouse monoclonal anti-α-Tubulin antibody (Santa Cruz Biotechnology, sc-8035; 1:10,000 dilution). The membrane was washed three times with TBST (Monad, CR10401S) for 5 min and then incubated with secondary antibodies for 1 hour at room temperature. The following secondary antibodies were used in this study: goat anti-rabbit IgG-HRP secondary antibody (Sino Biological, SSA004; 1:10,000 dilution) and goat anti-mouse IgG-HRP secondary antibody (Sino Biological, SSA007; 1:10,000 dilution). The signal was detected using High-sig ECL Western Blotting Substrate (Tanon, 180-5001).

### Knockdown efficiency evaluation in RGPs

#### In utero electroporation

To evaluate the knockdown efficiency of *Foxk1* shRNA in RGPs, the shRNA plasmid (*pLL3.7-shFoxk1-EGFP*) or scrambled control plasmid (*pLL3.7-EGFP*) was electroporated with TF overexpression plasmid (*pCAG-Foxk1-HA-IRES-tdTomato*) at a 1:1 ratio. To evaluate the knockdown efficiency of *Foxk2* shRNA in RGPs, shRNA plasmid (*pLL3.7-shFoxk2-EGFP)* or scrambled control plasmid (*pLL3.7-EGFP*) was electroporated with TF overexpression plasmid (*pCAG-Foxk2-HA-IRES-EGFP*) at a 1:1 ratio. The plasmids were constructed as described in the section “Single-cell transcriptomics and functional analysis of *Hbp1* and *Foxk1*, Plasmids and in utero electroporation for *Foxk1* in functional analysis”. The procedure for in utero electroporation was described in the section “Reference regulations, Plasmids and TF overexpression”.

#### Immunohistochemistry and imaging

The procedures for the collection of brain sections and immunostaining were described in the section “Immunohistochemistry and imaging”.

### Functional silencing of specific CRE using deactivated Cas9-KRAB system

#### Plasmids and in utero electroporation

We first generated *EF1α-MCP-KRAB-T2A-tdTomato-U6-deadsgRNA-2xMS*2 construct, which contains an EF1α promoter to express MCP-KRAB and tdTomato reporter and a separate U6 promoter that drives the expression of truncated sgRNA-2xMS2. The truncated sgRNA (Table S5) only guides Cas9 targeting without activating Cas9 for cleavage^153^. The sgRNA backbone incorporates MS2 binding sites, which recruit MCP fused KRAB to silence CREs at the sgRNA targeting site. This system is referred to as the deactivated Cas9-KRAB system. To functionally silence specific CREs, we used in utero electroporation to introduce the deactivated Cas9-KRAB system together with targeting sgRNA into the cortex of Cas9 mice. The procedure for in utero electroporation was described in the section “Reference regulations, Plasmids and TF overexpression”.

#### Bulk RNA-seq library preparation and qPCR experiments

The library preparation for analysis of CRE functional silencing was described in the section “Reference regulations, Bulk RNA-seq library preparation for TF overexpression experiments”. qPCR was used to analyze the changes in the target gene expression following CRE silencing and was performed as described in “Luciferase reporter assay, qPCR” with gene specific qPCR primers (Table S5). The quantified levels of target gene transcripts were normalized to *Actb*.

#### Bulk RNA-seq analysis of CRE functional silencing

To examine if *Pygo2* or *Nek6* CRE functional silencing could affect RGP transition states, we calculated the enrichment score (P value and odds ratio using Fisher’s exact test) of genes altered by turning off specific CREs in DE genes during the E12-E13 transition by using “fisher.test” function in R. Briefly, genes altered by turning off specific CREs were obtained as described in the section “Reference regulations, Processing of overexpression bulk RNA-seq” with following modifications: differential expression was estimated using DESeq2 package (version 1.26)^154^, and genes with absolute log2 Fold change > 0.25 and adjusted P value < 0.05. The identification of DE genes during the E12-E13 transition is described in the section “Transition-wise analysis of regulatory links”.

#### Identification of upstream TFs

To identify upstream TFs, we first calculated *AN*_TF_ of each TF, defined as the average of the number of regulatory links of downstream TF targeted by the TF.

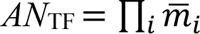

where *m*∽_*i*_ is average number of regulatory links of downstream TFs targeted in each transition *ii*.

The numbers of regulatory links of downstream TFs were normalized as fractions of the maximum in each transition. We identified upstream TFs based on three criteria: (1) The upstream TF regulates TFs (*AN_TF_* > 0) in all transitions. (2) The upstream TF is not listed among genes strongly induced by cell preparation procedures^155^. (3) The upstream TF displays salient changes (differential motif accessibility or differential expression) in more than one transition.

Differential expression was analyzed using the Wilcoxon rank-sum test, with P value < 0.05, absolute log2 fold change > 0.25. Differential motif accessibility was analyzed using the Wilcoxon rank-sum test, with P value < 0.05, absolute log2 fold change > 0.65.

#### Identification of upstream miRNAs

To identify upstream miRNAs, we first calculated *AN*_miRNA_ of each miRNA, defined as the average of the number of regulatory links of downstream TF targeted by the miRNA (*AN*_miRNA_).

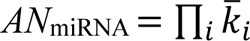

where *kk*^∽^_*ii*_ is the average number of regulatory targets of downstream TFs regulated by a miRNA in each transition *i*. The numbers of regulatory links of target TFs were normalized as fractions of the maximum in each transition. The miRNA-TF regulatory links were obtained from miRDB database (version 6.0)^116^. We identified upstream a miRNA based on two criteria: (1) the upstream miRNA regulates target TFs (*AN*_miRNA_ > 0) in all transitions; (2) The miRNA is expressed in cortex (counts > 1). The miRNA expression data was obtained from a publicly available cortex miRNA-seq database GSE142253^156^.

#### ChIP-seq enrichment analysis of motif occurrence

To examine ChIP-seq signal enrichment at CRE, publicly available ChIP-seq data was downloaded from GEO Accessions (GSE66961, GSE104247, GSE99818, GSE63282, GSE78720, GSE111657, GSE146961, GSE117997 and GSE127913). The processing of downloaded fastq reads and alignment was the same as described in the section “ChIP-seq data processing”. ChIP-seq and background signals were calculated on individual samples using MACS2 (version 2.2.7.1) with following parameters: --bdg --keep-dup all. Specifically, we obtained ChIP-seq signals as pileup reads from ChIP-seq sample and the background signals from dynamic background read distribution estimated from ChIP-seq sample by MACS2, as previously described^157^. Then we examined the difference between the ChIP-seq and background signals in genomic windows centered on each motif occurrence within CREs. The motif position was identified using FIMO (MEME suite v5.0.4) with the following settings: a first-order Markov background model, a P value cutoff of 10^-3^, and PWMs from the mouse HOCOMOCO motif database (v11). The statistical significance of the difference between max ChIP-seq signal and background signal at the motif occurrence was assessed using “ppois” function in R with the parameter lower.tail = F.

### Simulation with global temporal regulator

#### Global temporal regulator

We introduced an additional TF node to model the regulatory effect of the global temporal regulator., i.e., it can regulate *L* different TFs. We set *L* = 20 and drew the maximal contributions of global temporal regulator to these *L* different TFs from the uniform distribution *u*(−*K*_0_, *K*_0_). Those *L* TFs are chosen to be the TFs with the highest degree unless otherwise stated. Simulations were performed with *K*_0_ drawn from {1, 2, 5, 20} and yielded quantitively comparable staged patterns.

#### Simulating expression data with global temporal regulator

We first generated the TRN using the procedure discussed in “Simulation with non-temporal or temporal regulatory links, Generation of synthetic transcription regulation network (TRN)”. The dynamic model and parameterization were the same as described in the section “Simulation with non-temporal or temporal regulatory links”. The simulation was initialized with 1,000 steps with the expression of the global temporal regulator set to 0 to allow the gene expressions to reach steady states. Then, to examine the effect of patterns of expression trajectory of the global temporal regulator on the stage transition, we simulated the gene expression profiles of the global temporal regulator using three different progressively increasing expression patterns as follows *x*_lr_ = *m*_0_ *m* log(*y*): (1) normal (*m* = 1) (2) Over-expression (*m* = 2) and (3) Knockdown (*m* = 0.5). We chose *y* to be 5,000 evenly spaced values in the interval [1, 1.5], and each value corresponds to a time step during simulation.

#### Human single cell RNA-seq data analysis

Expression matrix (TPM) of human cortical development was collected from GSE104276. Dimension reduction and clustering analysis was performed similarly as described in the section “Dimensionality reduction and cell clustering, RNA only”. The resolution of “FindClusters” function was set as 1.8. The RGP cell clusters were identified by the expression of known RGP markers PAX6 and SOX2^158^. The expression of each NFI genes in the RGP cluster was computed as cluster average for each embryonic development stage.

## SUPPLEMENTARY FIGURE

**Figure S1-1:**
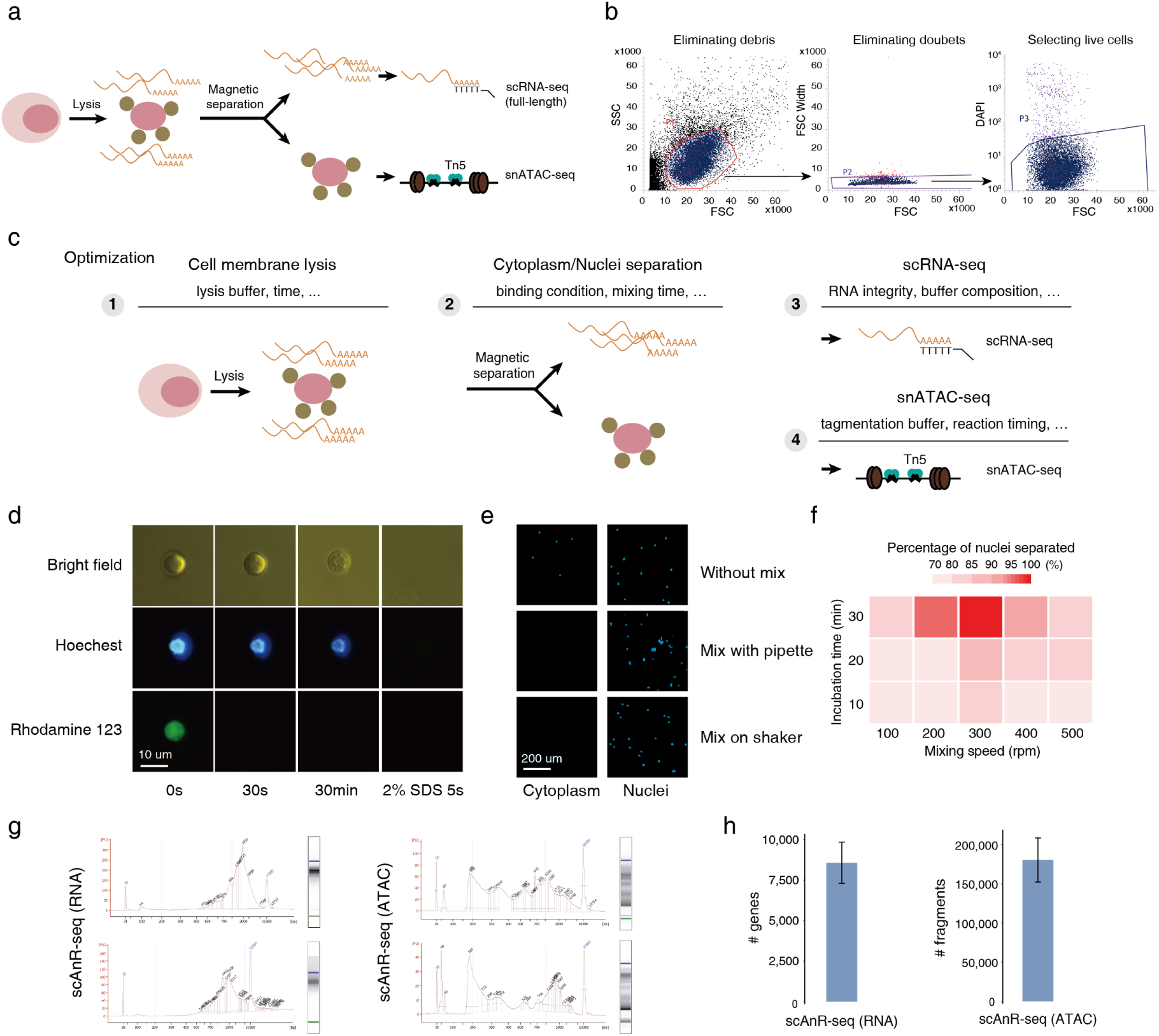
scAnR-seq optimization. **(a)** Schematic of scAnR-seq. **(b)** Representative FACS plots depicting the isolation of live cells from single cell suspension of the ventricular zone of cortex. **(c)** Schematic for optimization of scAnR-seq. **(d)** Intact nucleus during selective lysis of cell membrane; Rhodamine 123, dye impermeable to cell membrane; Hoechest, nuclear DNA staining. **(e)** Separation of cytoplasmic and nuclear contents. The nuclei were pre-stained with Hoechest (blue), extracted by nucleus-binding beads, and imaged to examine cytoplasmic and nuclear separation. Three methods for beads extraction are labeled. **(f)** Heatmap showing cytoplasmic and nuclear separation efficiency under combinatorial optimization for mixing speed and incubation time. **(g)** Representative DNA size distribution of pre-amplified RNA-seq (left) and ATAC-seq library (right) from a split single cell. **(h)** Quantification of genes (left) and chromatin fragments (right) captured by scAnR-seq. An average of 8,579 genes and 180,477 ATAC accessible fragments were detected. Shown are mean ± std.

**Figure S1-2:**
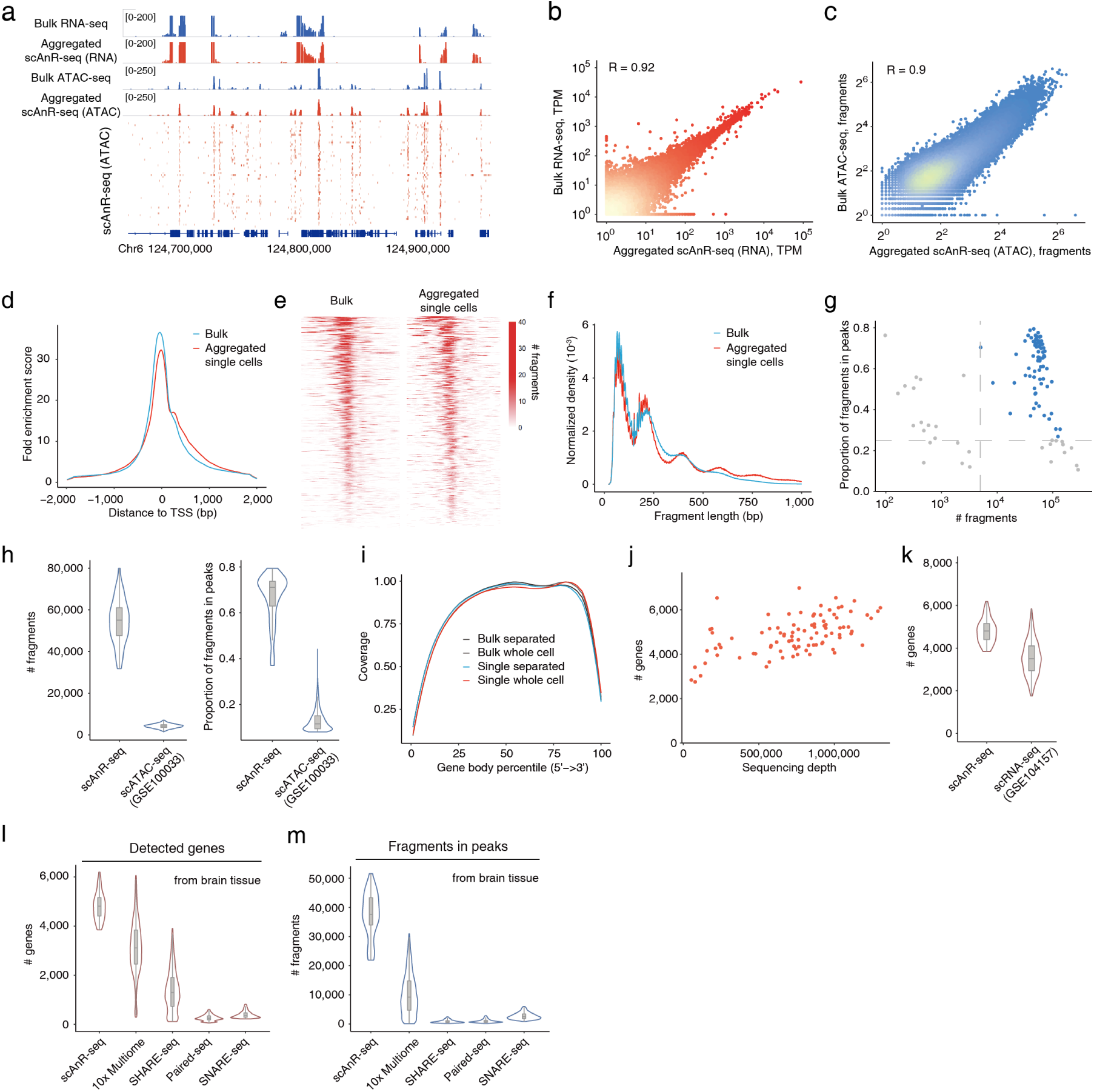
scAnR-seq significantly outperforms leading technologies. **(a)** Matched bulk and aggregated scAnR-seq profiles of cells from brain tissue. **(b and c)** scAnR-seq accurately represents both omics, showing genes (b) and chromatin fragments (c) from aggregated scAnR-seq highly correlated with bulk RNA-seq in cells from brain tissues. TPM and fragments counts were added a pseudo count of 1. **(d)** Strong enrichment of scAnR-seq (ATAC-seq portion) fragments around TSSs (32-fold in aggregate). **(e)** Enrichment of chromatin accessibility signals around TSSs for bulk and aggregated single cell profiles. **(f)** Insert-size distribution of scAnR-seq (ATAC-seq portion) fragments, exhibiting characteristic ATAC-seq distribution patterns. **(g)** In-peak ratio versus total captured fragments in single cells. **(h)** Violin plot showing distribution of total fragments and in-peak ratio in scAnR-seq, significantly surpassing previous technology (GSE100033). P value < 3.8×10^-35^ for total fragments and P value < 2.0×10^-34^ for in-peak ratio. Wilcoxon rank-sum test. **(i)** Distribution of mRNA reads along gene body, showing full length coverage. **(j)** The number of genes detected and sequencing depth in scAnR-seq. **(k)** Violin plot showing distribution of gene counts (TPM>0) in scAnR-seq, comparable to stand-alone Smart-Seq2 experiments (GSE104157). **(l-m)** Number of detected genes (L) or chromatin fragments in peaks (M) for scAnR-seq (this study) significantly surpass those from 10x Multiome (https://www.10xgenomics.com/resources), SHARE-seq^36^, Paired-seq^37^ or SNARE-seq^35^.

**Figure S1-3:**
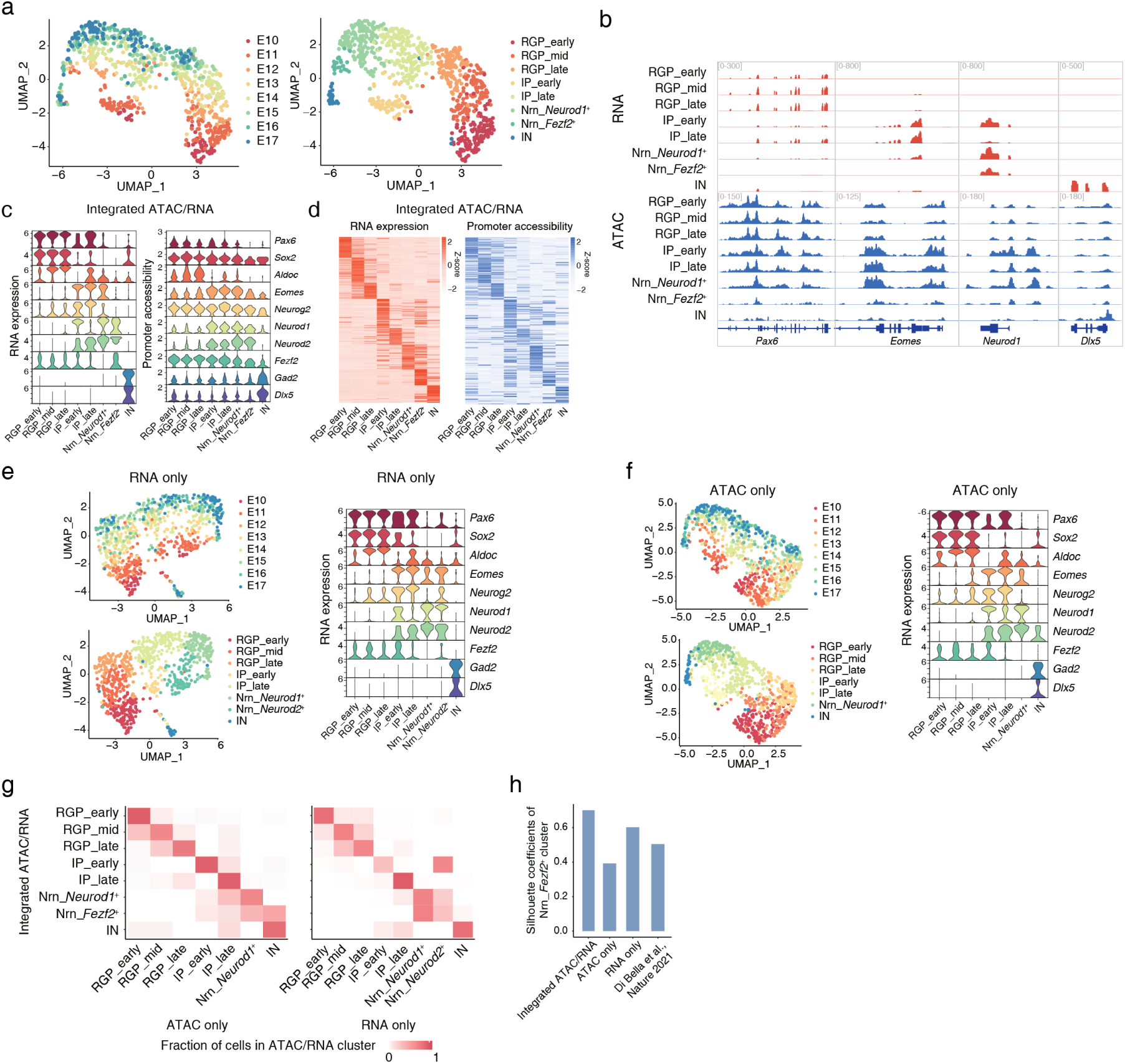
Integrated ATAC and RNA captured additive gain of cellular information. **(a)** Single cells on uniform manifold approximation and projection (UMAP) coordinates defined by integrated RNA and ATAC. Cells are colored by sampling-time (left) or cell type assignment (right), n = 817. **(b)** Aggregated gene expression and chromatin accessibility profiles for each cell cluster at loci of representative marker genes. **(c)** Violin plots showing the markers of cell types assigned by integrated RNA and ATAC. Left: RNA expression; Right: promoter accessibility. **(d)** Heatmaps showing cell-type specific genes. Matched gene expression levels (left) and promoter accessibility (right). **(e)** Left: UMAP coordinates defined using only RNA data. Cells are colored by sampling time (top) or cell type assignment (bottom). Right: Violin plots showing the markers of cell types assigned using only RNA data. **(f)** Left: UMAP coordinates defined using only ATAC data. Cells are colored by sampling time (top) or cell type assignment (bottom). Right: Violin plots showing the markers of cell types assigned using only ATAC data. **(g)** Heatmaps showing overlap in cell-type assignment between clusters defined by ATAC only (left) or RNA only (right) and integrated ATAC and RNA. **(h)** Silhouette coefficients of Nrn_*Fezf2*^+^ clusters in UMAP defined by integrated ATAC/RNA, ATAC only, RNA only, and a public RNA dataset^27^.

**Figure S1-4:**
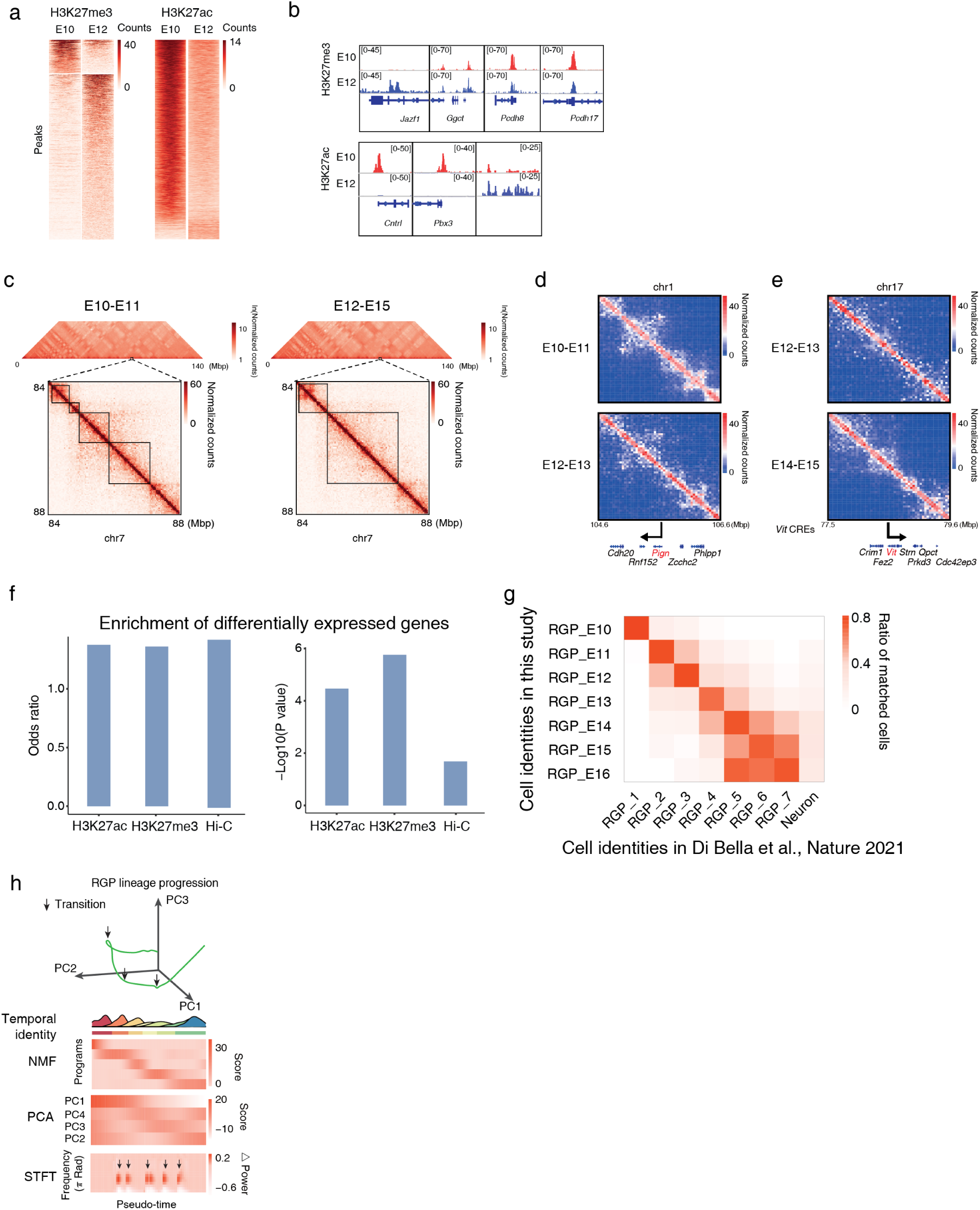
Pseudo-time alignment of scAnR-seq profiles recapitulate RGP progression. **(a)** Heatmaps showing differential H3K27me3 (left) or H3K27ac (right) ChIP-seq signal. **(b)** IGV view of differential H3K27me3 (top) or H3K27ac (bottom) ChIP-seq signals at representative loci. **(c)** Heatmaps showing normalized Hi-C contact counts in 1 Mb bins (top) and 40 kb bins (bottom) in chromosome 7. **(d)** *Pign* regions with decreased looping intensity during E12-E13 compared to E10-E11. **(e)** *Vit* regions with increased looping intensity during E14-E15 compared to E12-E13. **(f)** Differentially expressed genes along pseudo-time enriched in differential peaks of H3K27ac, H3K27me3, and differential Hi-C contact. Fisher’s exact test. **(g)** Cell type consistency for RGPs from scAnR-seq and a large-scale public dataset^27^. **(h)** Top: trajectory of RGP transitions in 3-dimensional PCA space. Middle: RGP temporal identity, gene program score from NMF, and PC component score from PCA analysis. Bottom: STFT analysis quantification of transitions along pseudo-time. Delta power was taken as the difference in STFT power spectrum between consecutive windows. Arrows point to spikes in frequency components.

**Figure S2-1:**
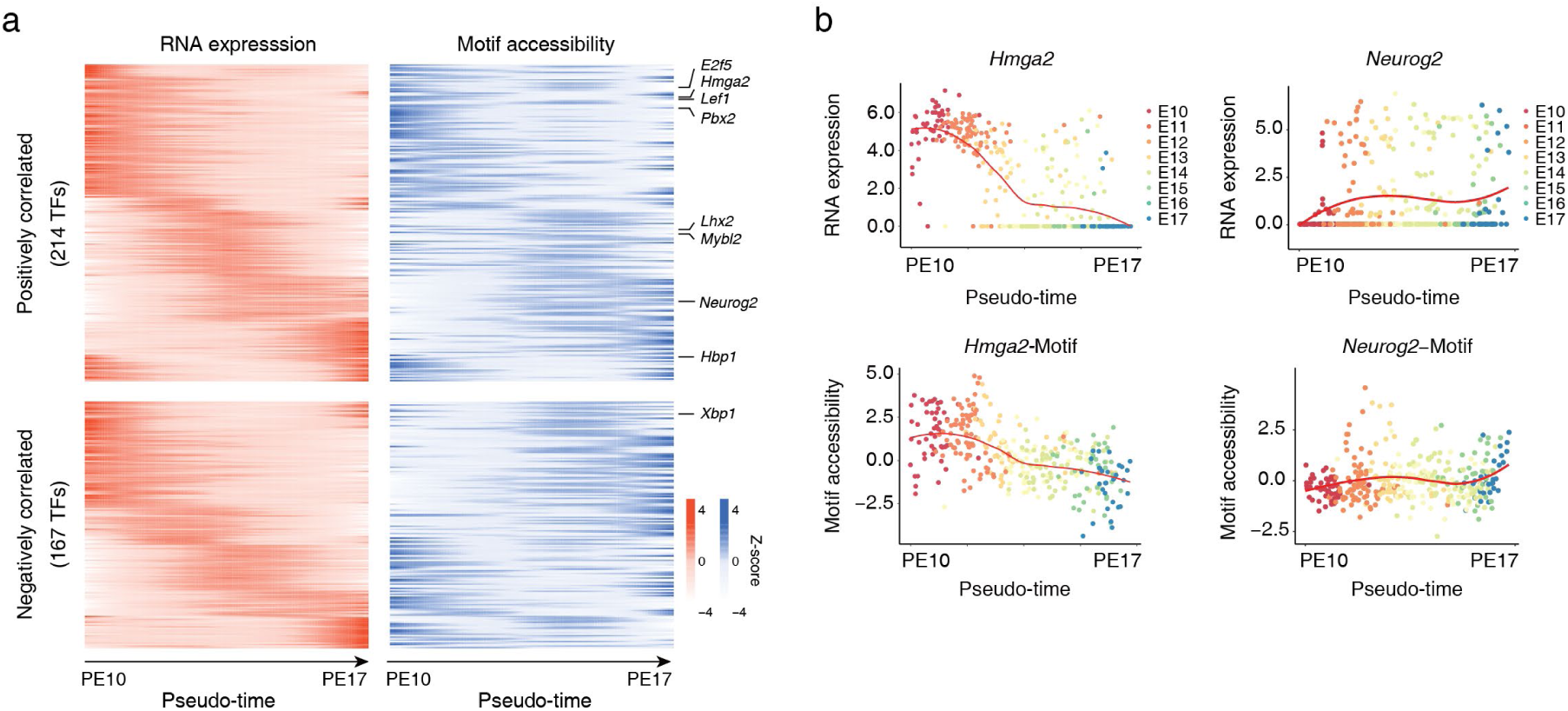
Globally coordinated dynamics in motif accessibility. **(a)** Clustering of TF expression (left) and motif accessibility (right) dynamics. Single cells ordered by pseudo-time. **(b)** Representative examples of coordinated TF activity and expression dynamics. Scatter plot showing gene expression (top) and TF motif accessibility (bottom) of *Hmga2* (left) and *Neurog2* (right) across pseudo-time during RGP development. Red line: linear regression fitting.

**Figure S2-2:**
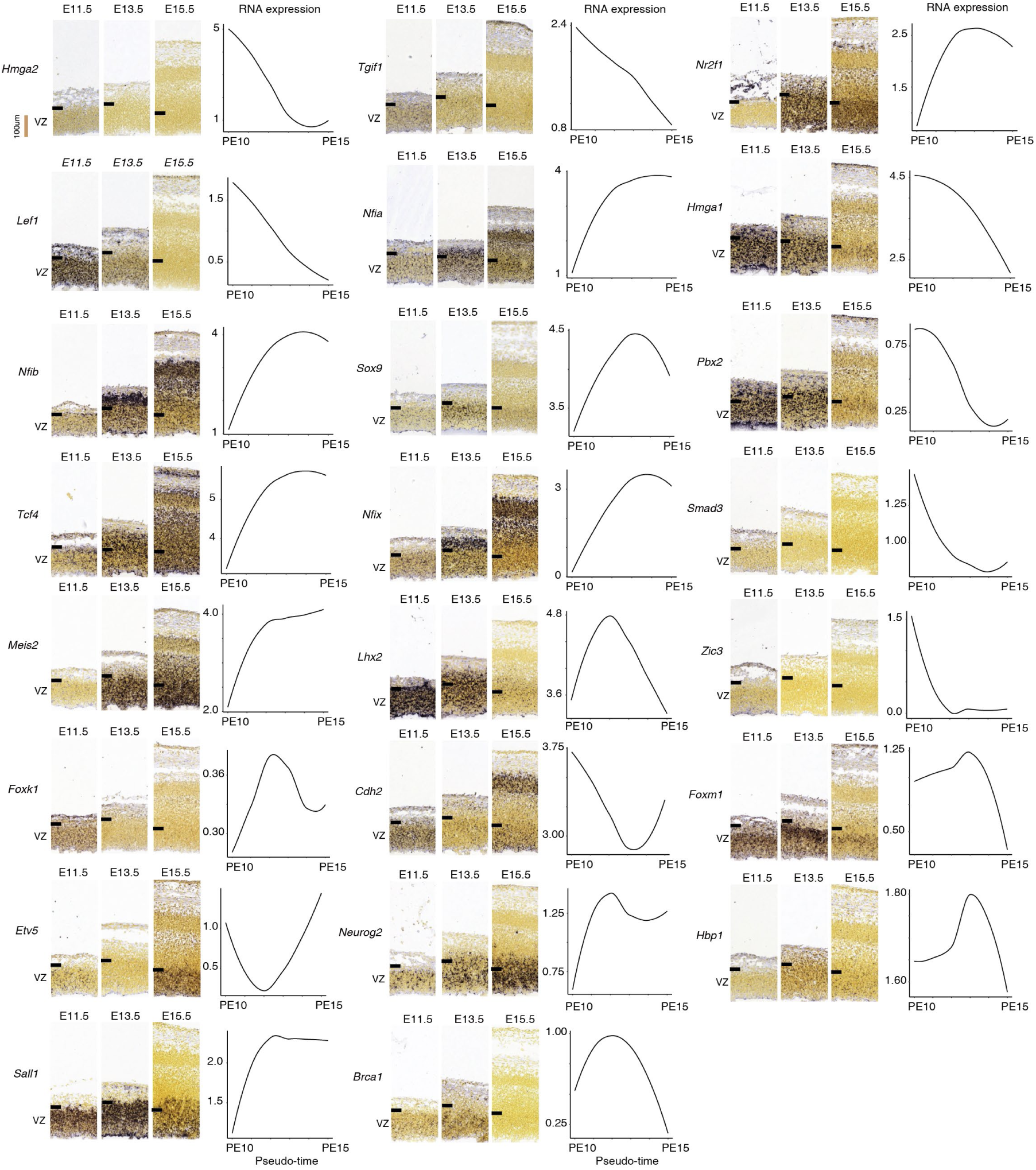
In situ hybridization temporal series. For each panel, left: TF gene symbol; middle: RNA in situ hybridization (ISH) of TF in mouse cerebral cortex at three developmental points (E11.5, E13.5, E15.5); source of ISH: Allen Developing Mouse Brain Atlas (https://developingmouse.brain-map.org/); right: RNA expression in scAnR-seq (black curve) covering the same development period. Tick mark indicates VZ basal border.

**Figure S2-3:**
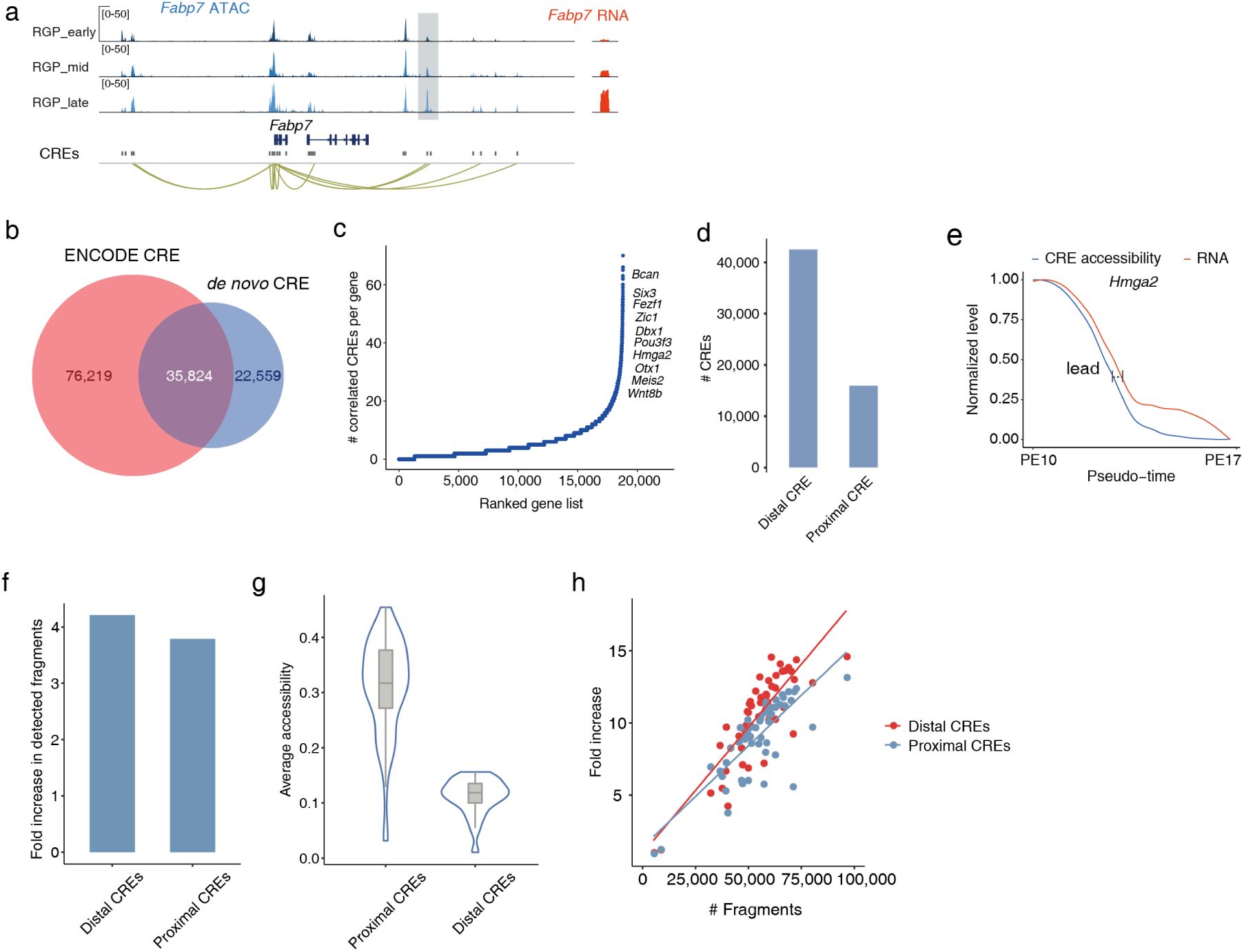
Globally coordinated dynamics in CREs. **(a)** Representative example of CRE and target gene association. Genome browser view of the *Fabp7* locus. Yellow loops: gene-CRE associations. Grey box: representative distal CREs of *Fabp7* showing accessibility changes correlated with *Fabp7* expression. **(b)** Venn diagram showing CREs overlapping with the ENCODE brain enhancer database^59^. **(c)** Distribution of the number of CREs associated with each gene, within 500 kb of the transcription start site (TSS). Known genes related to cortical development are highlighted. **(d)** The number of distal CREs (>500 bp from the TSS) is higher than that of proximal CREs (within 500 bp of the TSS). **(e)** CRE accessibility (blue) and gene expression (red) dynamics of *Hmga2* during its down-regulation, showing a modest lead in the CRE dynamics. **(f)** Fold increase in detected fragments in distal CREs by scAnR-seq compared to 10x Multiome is greater than that observed in proximal CREs. **(g)** Violin plot showing that average accessibility of distal CREs is lower than that of proximal CREs. Box inside violin plot denote 25th, 50th, and 75th percentiles; whisker length represents 1.53 interquartile ranges. **(h)** Fold increase in detected fragments of distal CREs with increasing sequencing depth is greater than that of proximal CREs.

**Figure S2-4:**
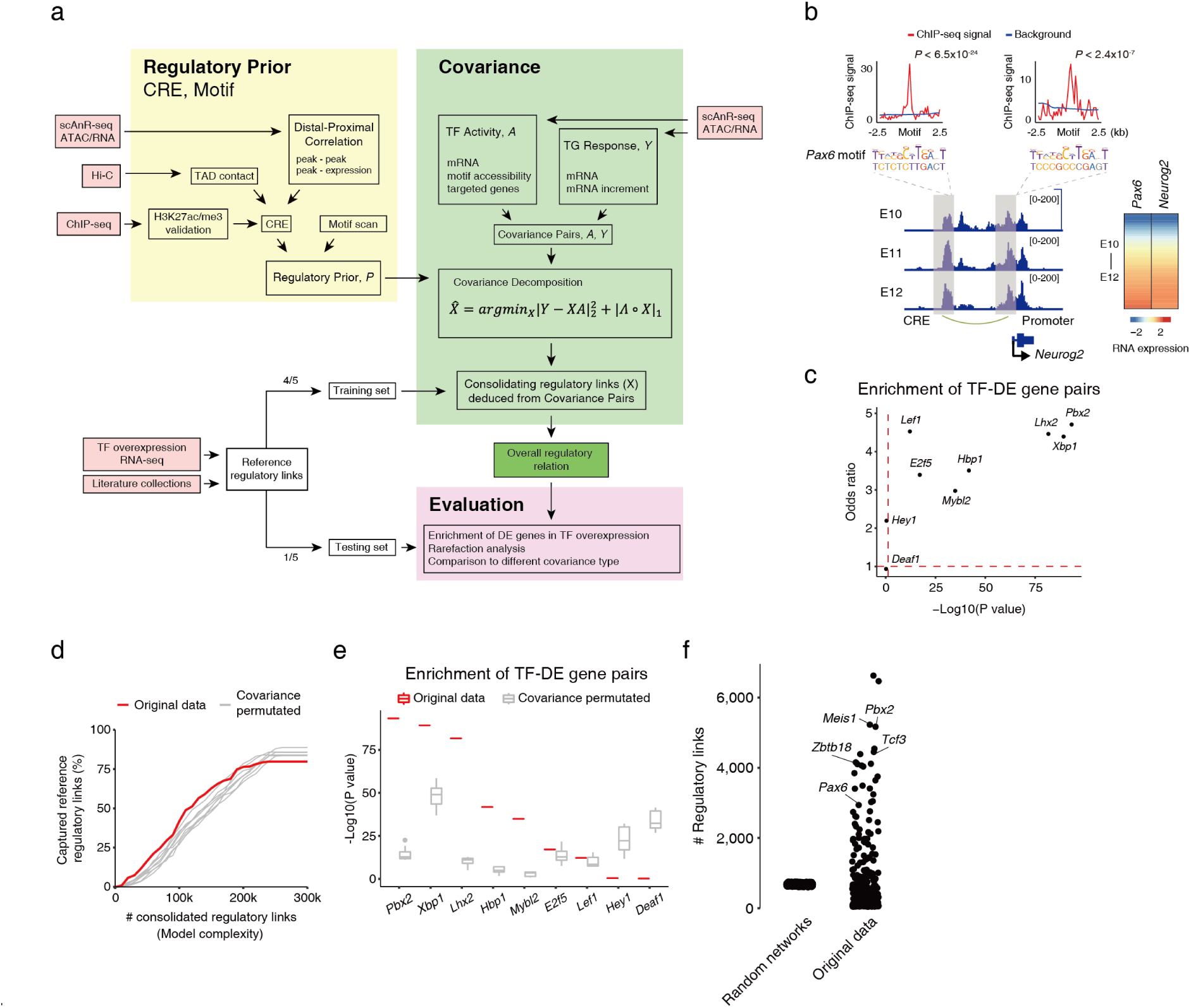
Multi-modal inference of regulatory links. **(a)** Overview of the multi-modal inferences, consolidation, and evaluation. We obtained reference regulatory links, by two conventional methods: (i) we selected 9 TFs that exhibited large variance in pseudo-time dynamics, overexpressed each TF in RGPs, and identified differentially expressed (DE) genes that were strongly induced (P value < 0.05, absolute log2 fold change > 0.4), obtaining a total of 1,722 TF-DE gene pairs as reference regulatory links; (ii) we manually curated TF-gene regulation supported by experimental evidence in the literature, obtaining 364 TF-gene pairs as reference regulatory links. **(b)** Overall regulatory relation recapitulated reference regulatory links, illustrated by *Pax6* to *Neurog2* regulatory link. Shown are co-varying dynamics in expression and chromatin accessibility of *Pax6* and *Neurog2*. Grey boxes indicate identified *Pax6* motif instances in the *Neurog2* proximal and distal CREs. The identified motif instances are valid *Pax6* binding sites, showing enrichment of *Pax6* ChIP-seq signal (GSE66961)^107^ in brain tissue. **(c)** Enrichment of differentially expressed genes induced by TF overexpression in TF target genes predicted by the overall regulatory relation. Fisher’s exact test. **(d)** Rarefaction analysis showing percentage of reference regulatory links captured versus number of consolidated regulatory links inferred from original data versus covariance permutated data (n = 20 trials). **(e)** Box plot showing enrichment of differentially expressed genes induced by TF overexpression in TF target genes predicted by the overall regulatory relation inferred from original data versus covariance permutated data (n = 20 trials). Fisher’s exact test. **(f)** Distribution of numbers of linked target genes per TF; The random networks were inferred from covariance permutated data (n = 200 trials; Methods.

**Figure S3-1:**
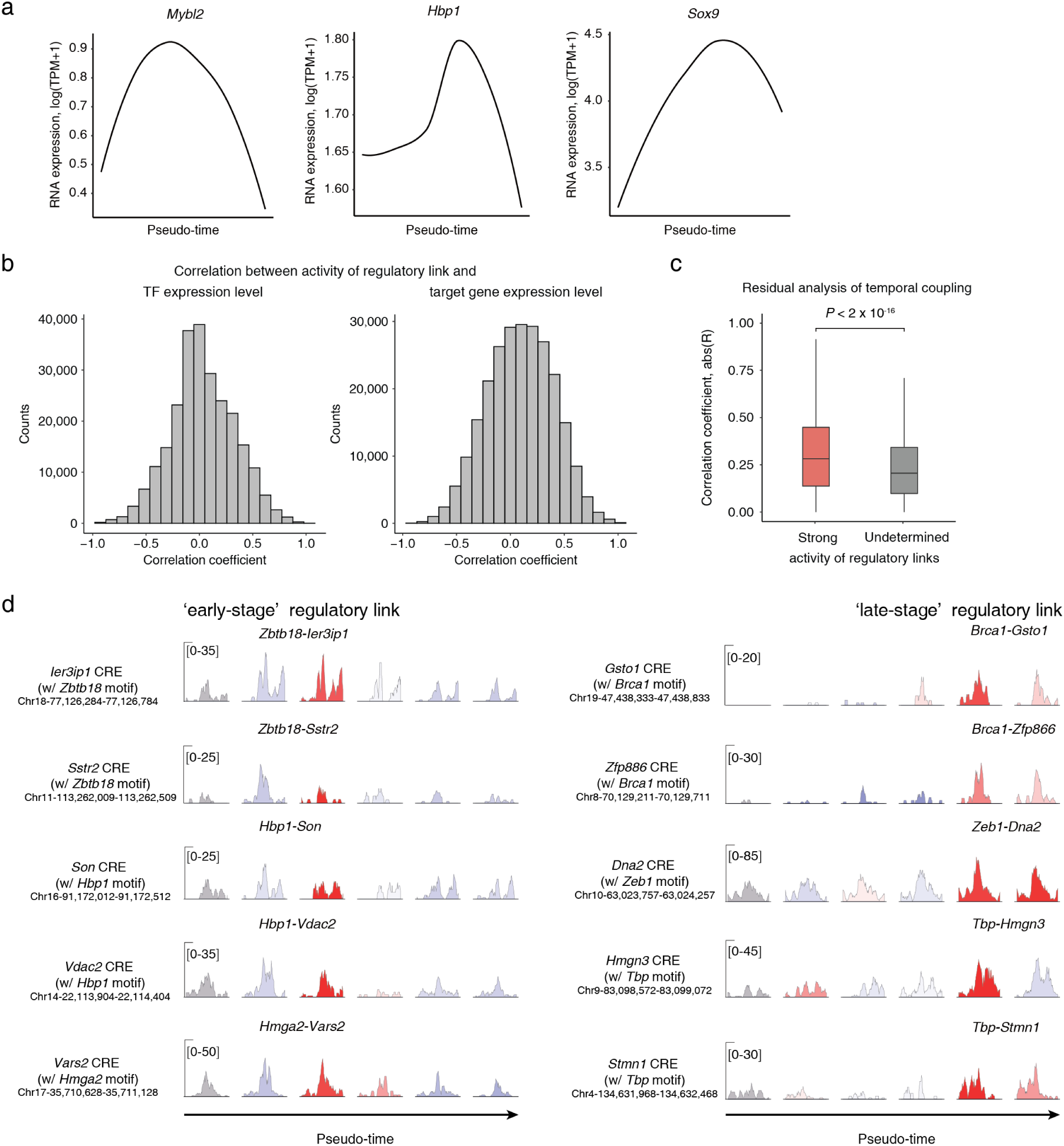
Characterization of activity of regulatory links. **(a)** Gene expression of *Mybl2*, *Hbp1*, *Sox9* along pseudo-time. **(b)** Distribution of Pearson correlation coefficient between activity of regulatory links and the corresponding TF (left) or target gene (right) expression level. **(c)** Box plot showing higher correlation between single cell residual variability of TFs and targets when the regulatory link exhibits strong activity. Activity of regulatory links was undetermined when TFs and genes exhibited little changes in expression (Methods). **(d)** More representative IGV track examples showing that accessibility increase at a CRE that links TF and target precedes strong activity of the regulatory link. Peaks are colored by activity strength of the regulatory link.

**Figure S4-1:**
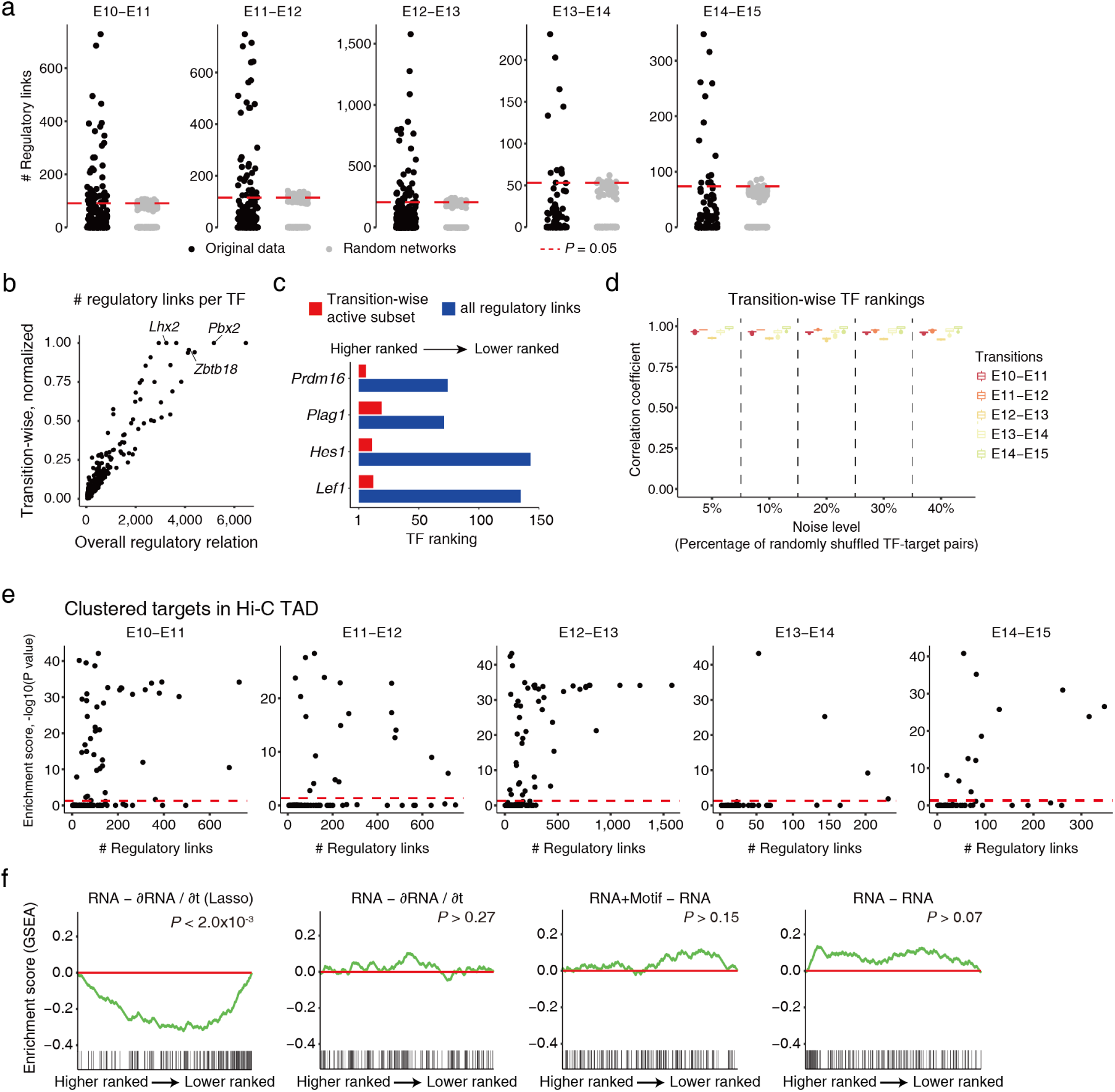
Structural, epigenetic, and functional characterization of transition-wise TF hierarchy. **(a)** Distribution of number of associated regulatory links per TF in transition-wise analysis. The random networks were inferred from covariance permutated data (n = 200 trials; Methods). Permutation test. **(b)** Top-ranked TFs in the overall regulatory relation were generally also top-ranked TFs in transition-wise analysis. Numbers of regulatory links by TF were normalized as fractions of the maximum in transition-wise analysis and max pooled across all transitions. **(c)** Bar plots showing *Prdm16*, *Plag1*, *Hes1* and *Lef1* promoted in transition-wise TF rankings. **(d)** Correlation of transition-wise TF rankings under perturbation, where TF-target links of the indicated percentages were randomly shuffled. Error bar indicates standard deviation in random trials, n = 50 trials. **(e)** Scatter plot showing the enrichment of linked targets in TAD (Methods) versus number of regulatory links by each TF in transition-wise analysis. Red dashed line indicates the significance level of P value = 0.05. Enrichment estimated by comparing targets in TAD and in randomized TAD, one-sample Wilcoxon rank-sum test. **(f)** GSEA running score in the gene set of regulators involved in cortical developmental defects (Table S4) for TFs ranked in regulation relation inferred based on RNA expression, as in Figure 3f. Kolmogorov-Smirnov test.

**Figure S4-2.**
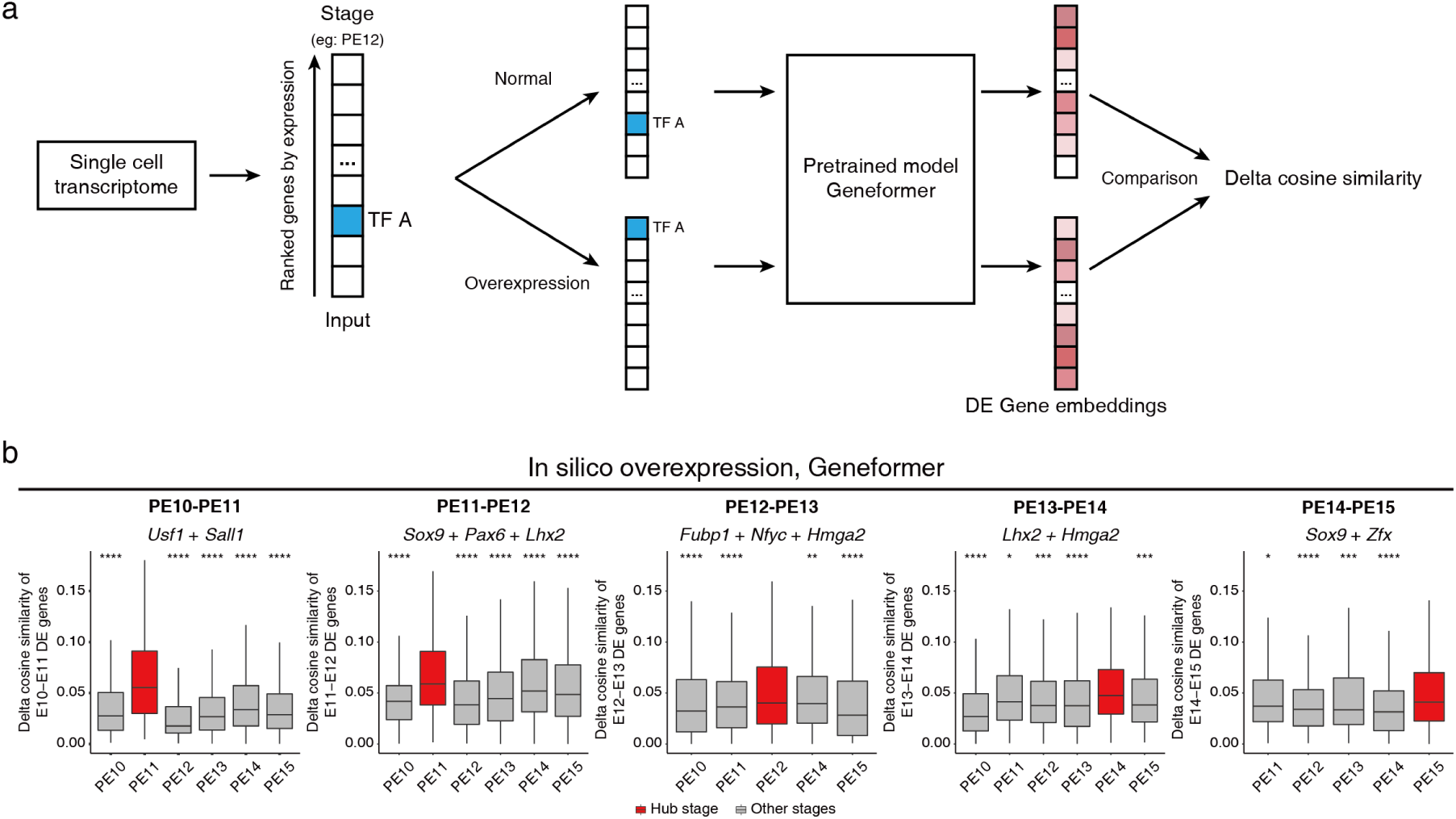
The hub TFs regulate developmental state-specific gene expression programs.**(a)** Diagram illustrating perturbation of hub TFs and analysis of their impact on state-dependent expression programs using GeneFormer. GeneFormer is a foundational biological model trained on 30 million single cell transcriptomic profiles (*46*). The input to GeneFormer consists of genes ranked by their expression levels. For the control group, genes are ranked based on single cell gene expression profiles specific to a RGP temporal state. For the perturbation group, the ranking of the hub TFs corresponding to that state are elevated to simulate upregulation of these hub TFs. These ranked genes are input into GeneFormer to obtain gene embeddings. Since the pretrained GeneFormer model captures regulatory relation among genes, upregulation of hub TFs affects the expression of other genes, reflected in changes in the gene embeddings. To quantify the magnitude of this impact, the delta cosine similarity between the gene embeddings of the control and the perturbation groups is calculated. **(b)** Using the method described in (A), the impact of hub TF upregulation on state-dependent gene expression was calculated for their corresponding states (red boxes) and non-corresponding states (gray boxes). Box plots display the delta cosine similarity across different single cells of the same temporal state. From left to right, shown are results for the perturbation of hub TFs correspond to states progressing from earlier to later stages.

**Figure S5-1:**
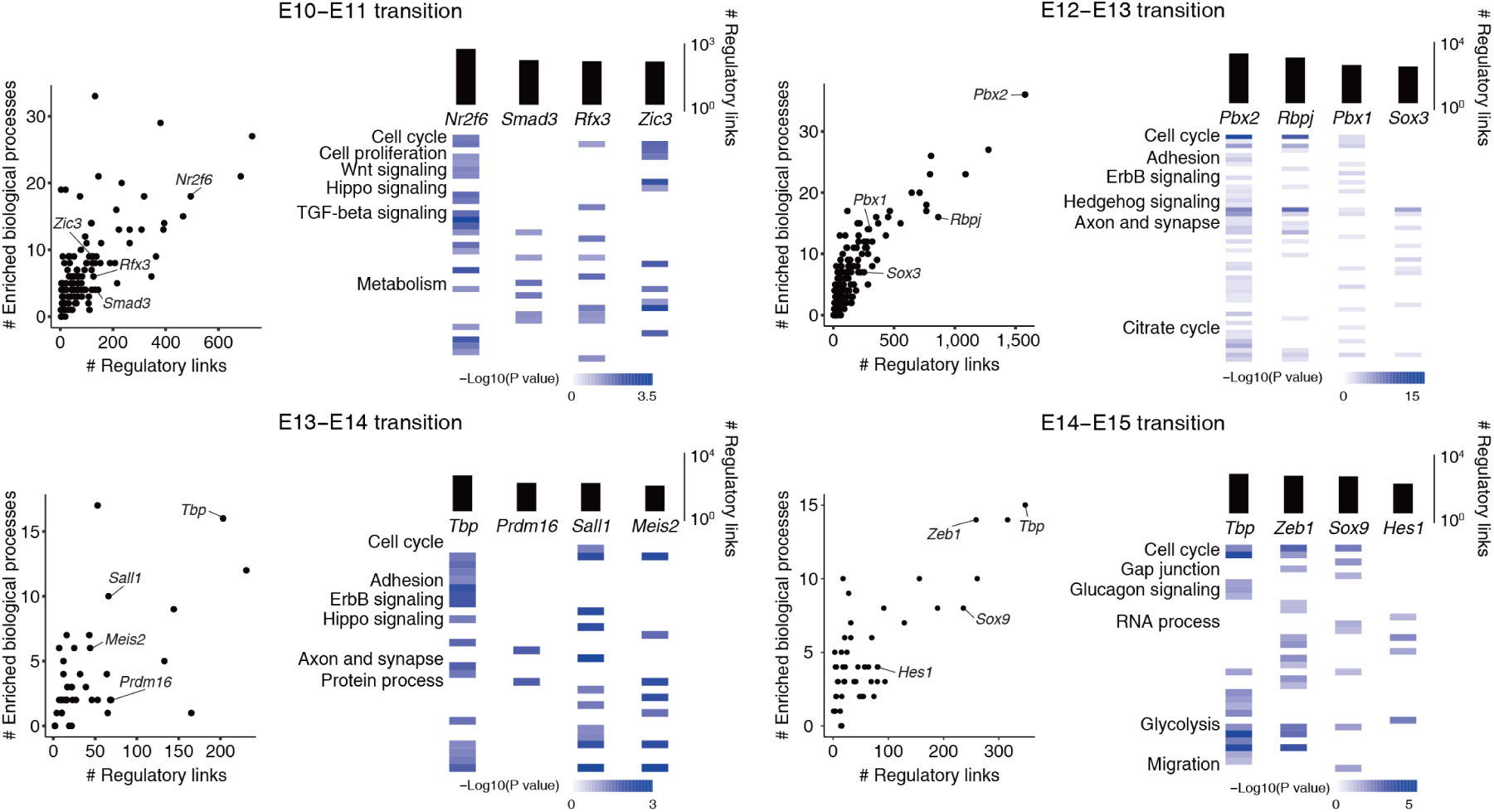
Hierarchical functional coordination. For each panel, left: scatter plot showing the number of enriched KEGG and GO biological processes by TF direct targets versus the number of regulatory links in indicated transitions. Right: heatmaps showing the enrichment score of KEGG and GO biological processes by direct targets of indicated TFs. Representative processes are provided. Top: bar plots showing the number of regulatory links.

**Figure S5-2:**
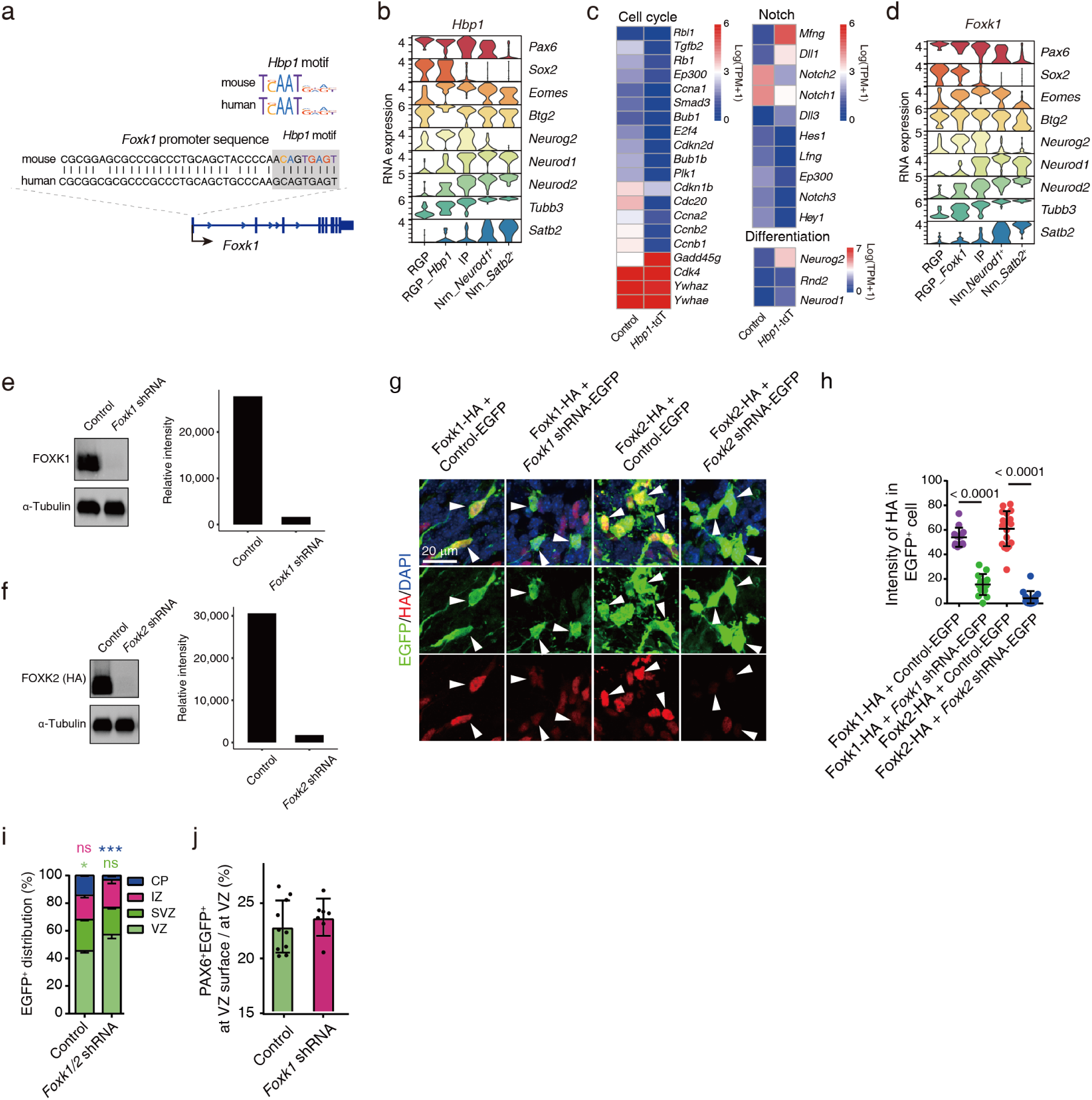
Coordination of key process by *Hbp1* and *Foxk1* in the transition-wise TF hierarchy. **(a)** Top: conservation of mouse and human *Hbp1* motif. Bottom: conservation of mouse and human *Foxk1* promoter sequence containing the identified *Hbp1* motif instance. Related to Figure 5c, shown is human HBP1 ChIP-seq signal at the identified motif instance. **(b)** Violin plots showing the markers of cell types assigned in *Hbp1* overexpression single-cell data integrated with control single-cell data. **(c)** Heatmaps showing the differentially expressed genes in cell cycle, Notch signaling pathways, and differentiation^51,108,109^ induced in *Hbp1* overexpressing RGPs. **(d)** Violin plots showing the markers of cell types assigned in *Foxk1* overexpression single-cell data integrated with control single-cell data. **(e)** Knockdown efficiency of *Foxk1* shRNA was evaluated by western blot in a cell line overexpressing *Foxk1*. Left: anti-FOXK1 and anti-ɑ-Tubulin western blot of control and *Foxk1* shRNA sample. Right: quantification of FOXK1 protein abundance in left. ɑ-Tubulin was used as an internal normalization control. **(f)** Knockdown efficiency of *Foxk2* shRNA was evaluated by western blot in a cell line overexpressing *Foxk2*-HA. Left: anti-HA for FOXK2 and anti-ɑ-Tubulin western blot of control and *Foxk2* shRNA sample. Right: quantification of FOXK2 protein abundance in left. ɑ-Tubulin was used as an internal normalization control. **(g)** Knockdown efficiency of *Foxk1* or *Foxk2* shRNA was evaluated by immunofluorescence in RGPs overexpressing *Foxk1*-HA or *Foxk2*-HA. Representative coronal sections of dorsolateral telencephalon triple stained using anti-GFP (green), anti-HA (red) for FOXK1 or FOXK2, and a DNA dye (blue) of control, *Foxk1* and *Foxk2* shRNA sample. **(h)** Quantification of immunofluorescence intensity for FOXK1 or FOXK2 in EGFP^+^ cells in (g). **(i)** Quantification of the distribution of EGFP^+^ cells in Figure 5i. Data are shown as mean ± SEM. *P=0.0185, ***P=0.0002; Student’s t-test; N.S., not significant. **(j)** Increase in the percentage of PAX6^+^EGFP^+^ cell on the ventral surface in EGFP^+^ cells in *Foxk1* knockdown samples.

**Figure S6-1:**
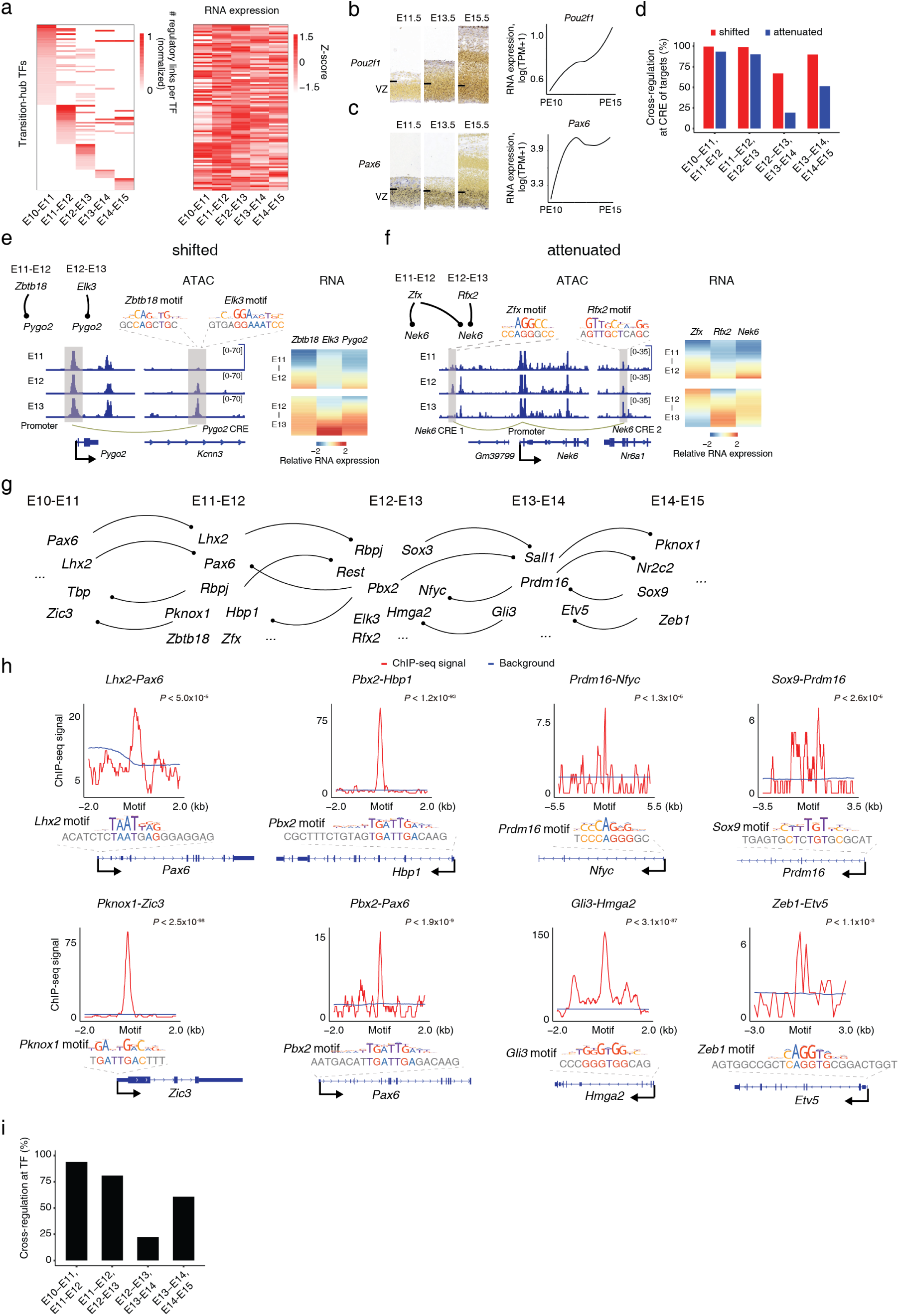
Cascade of sequential hub TFs cross-regulated at network level. **(a)** The transition hub TFs do not exhibit expression patterns in strict temporal windows. Left, heatmap showing numbers of regulatory links by TFs normalized as fractions of the maximum in transition-wise analysis. Right, heatmap showing the TF expression in each transition. **(b and c)** Expression of transition hub TFs, *Pou2f1* (b) and *Pax6* (c), are not in strict temporal windows. left: RNA in situ hybridization (ISH) of indicated TFs in mouse cerebral cortex at three developmental points (E11.5, E13.5, E15.5); source of ISH: Allen Developing Mouse Brain Atlas (https://developingmouse.brain-map.org/); Tick mark indicates VZ basal border. Right: RNA expression in scAnR-seq (black curve) covering the same development period. **(d)** Percentages of shifted or attenuated expression dynamics with cross-regulation at CREs of targets by transition hub TFs of adjacent transitions. **(e)** Shifted expression dynamics of *Pygo2*, cross-regulated by *Zbtb18* and *Elk3*, transition hub TFs of adjacent transitions. (Right) Initially, *Pygo2* expression was up-regulated in concert with an increase in *Zbtb18* expression. In the next transition, *Pygo2* expression was decoupled from that of *Zbtb18* and continued increase, along with a concomitant increase in *Elk3* expression. (Left) the accessibility of cognate CRE at the *Pygo2*, indicating *Pygo2* regulation shifted from *Zbtb18*-mediated to *Elk3*-mediated regulation. **(f)** Attenuated expression dynamics of *Nek6*, cross-regulated by *Zfx* and *Rfx2*, transition hub TFs of adjacent transitions. (Right) *Nek6* and *Zfx* expression both increased initially, but were then decoupled in the next transition, when the *Rfx2*-specific CRE at *Nek6* became increasingly accessible (Left). This change in chromatin accessibility likely counteracted *Zfx2* down-regulation, leading to an attenuated *Nek6* expression dynamics and hence a stable expression level. **(g)** Model of cascading of transition hub TFs. **(h)** Cross-regulation at CREs of transition hub TFs outside of their hub stages. For each regulation pair, the gene model and occurrence of a motif of the regulator at a target CRE are shown. The identified motif occurrences are valid binding sites, showing enrichment of regulator ChIP-seq signal (GSE99818^110^; GSE63282^111^; GSE78720^23^; GSE111657^112^; GSE146961^113^; GSE117997^114^; GSE127913^115^) in a cell line (for *Pknox1*, *Sox9*) or in tissue (for *Lhx2*, *Pbx2*, *Prdm16*, *Gli3*, *Zeb1*). **(i)** Extending the analysis in (d) for attenuated expression dynamics of transition hub TFs outside of their hub stages.

**Figure S6-2:**
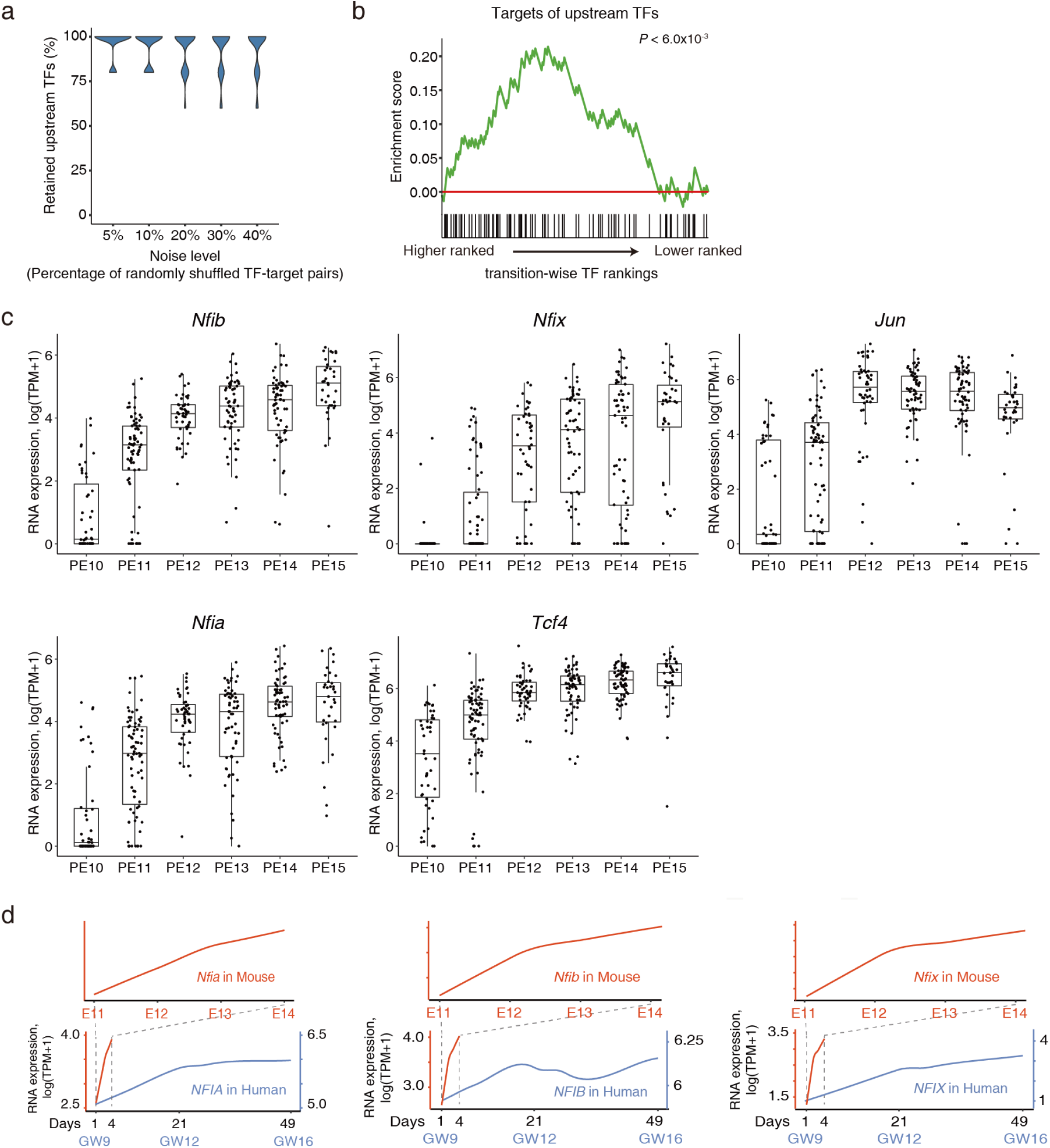
Progressively increasing pattern of global temporal regulators along pseudo-time. **(a)** Violin plot showing percentage of retained common upstream TFs under perturbation, where TF-target pair of the indicated percentage were randomly shuffled. Error bar indicates standard deviation in random trials, n = 50 trials. **(b)** Gene set enrichment analysis (GSEA) running enrichment score for targets of the common upstream TFs, based on transition-wise TF rankings, max pooled across all transitions. P < 6.0 × 10^-3^. Kolmogorov-Smirnov test. **(c)** Gene expression of global temporal regulators along pseudo-time, related to Figure 6b. **(d)** Progressively increasing dynamics of Nfia, Nfib or Nfix expression in mouse (red) or human single-cell RNA-seq data (blue; GSE104276) (Zhong et al., 2018) along aligned cortical development. Gestational weeks (GW); Embryonic day (E).

**Figure S6-3:**
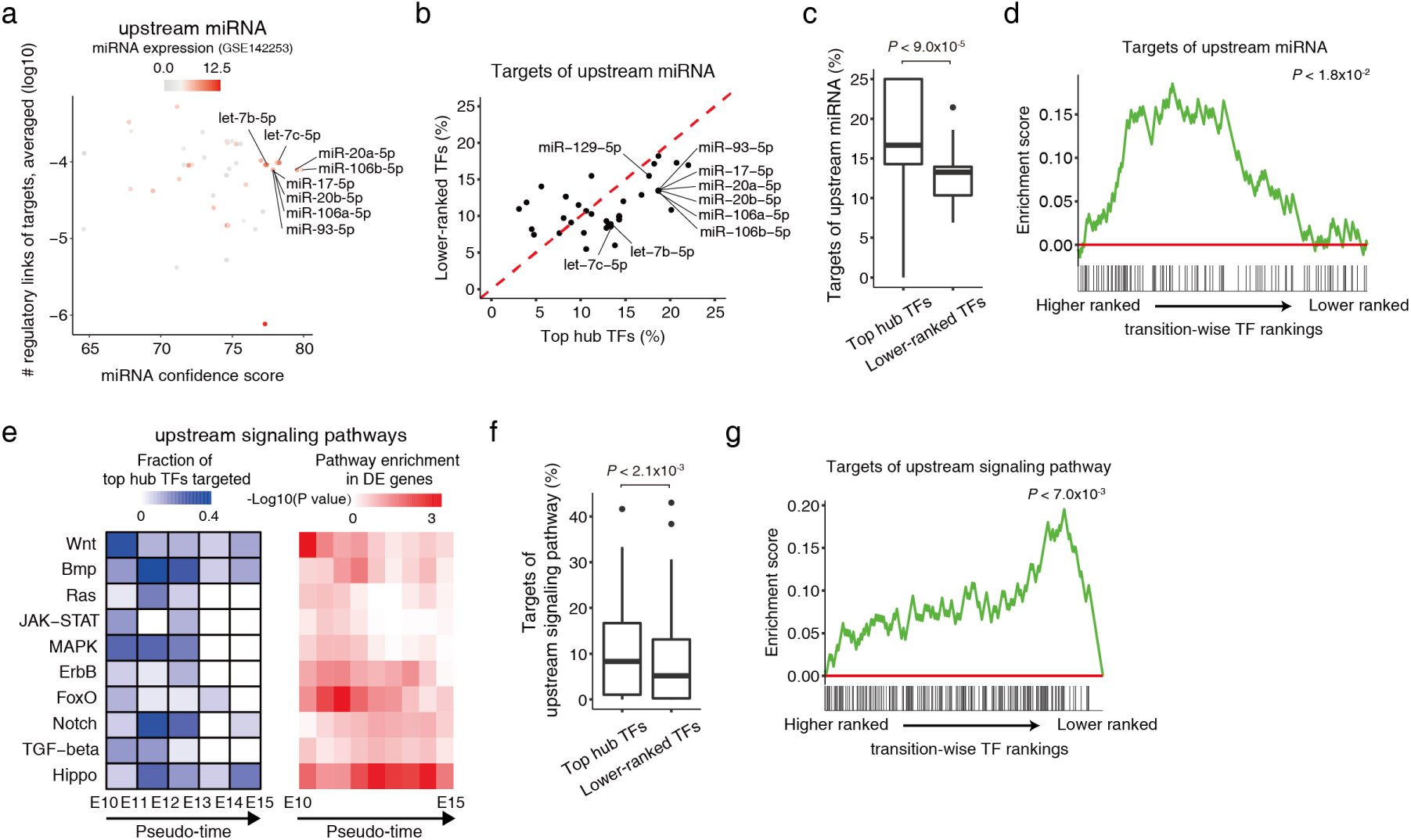
Functional characterization and expression of additional global temporal regulators. **(a)** Scatter plot showing the average number of regulatory links of targets versus the confidence of cognate sites in targets per upstream miRNA^116^. The numbers of regulatory links of targets by each upstream miRNA were normalized as fractions of the maximum in transition-wise analysis and averaged across all transitions. Known miRNA regulators in cortical development are highlighted. **(b)** Scatter plot showing the average percentage of top transition-hub TFs versus lower-ranked TFs regulated by upstream miRNA. The upstream miRNA are labeled as in (a). Percentage averaged across transitions. **(c)** The upstream miRNA that are highlighted in (a) regulate a higher percentage of top transition-hub TFs than that of lower-ranked TFs. P < 9.0 × 10^-5^. One-sided paired Wilcoxon rank-sum test. Box plots denote 25th, 50th, and 75th percentiles. **(d)** Gene set enrichment analysis (GSEA) running enrichment score for targets of upstream miRNA that are highlighted in (a), based on transition-wise TF rankings, max pooled across all transitions. P < 1.8 × 10^-2^. Kolmogorov-Smirnov test. **(e)** Left: heatmap showing the fraction of top transition-hub TFs targeted by upstream signaling pathways along pseudo-time. Right: enriched upstream signaling pathways in differentially expressed genes along pseudo-time (Methods). **(f)** The upstream signaling pathways regulate a higher percentage of top transition-hub TFs than that of lower-ranked TFs. P < 2.1 × 10^-3^. One-sided paired Wilcoxon rank-sum test. Box plots denote 25th, 50th, and 75th percentiles. **(g)** GSEA running enrichment score for targets of upstream signaling pathway, based on transition-wise TF rankings, max pooled across all transitions. P < 7.0 × 10^-3^. Kolmogorov-Smirnov test.

---

**Supplementary Table S1.** scAnR-seq optimization parameters

**Supplementary Table S2.** Reference regulatory links

**Supplementary Table S3.** Transition-wise TF rankings

**Supplementary Table S4.** TFs associated with cortical developmental defects

**Supplementary Table S5.** sgRNA and primer sequences

## Notes

### Competing Interest Statement

The authors have declared no competing interest.

